# SPLASH: a statistical, reference-free genomic algorithm unifies biological discovery

**DOI:** 10.1101/2022.06.24.497555

**Authors:** Kaitlin Chaung, Tavor Z. Baharav, George Henderson, Ivan N. Zheludev, Peter L. Wang, Julia Salzman

## Abstract

Today’s genomics workflows typically require alignment to a reference sequence, which limits discovery. We introduce a new unifying paradigm, SPLASH (Statistically Primary aLignment Agnostic Sequence Homing), an approach that directly analyzes raw sequencing data to detect a signature of regulation: sample-specific sequence variation. The approach, which includes a new statistical test, is computationally efficient and can be run at scale. SPLASH unifies detection of myriad forms of sequence variation. We demonstrate that SPLASH identifies complex mutation patterns in SARS-CoV-2 strains, discovers regulated RNA isoforms at the single cell level, documents the vast sequence diversity of adaptive immune receptors, and uncovers biology in non-model organisms undocumented in their reference genomes: geographic and seasonal variation and diatom association in eelgrass, an oceanic plant impacted by climate change, and tissue-specific transcripts in octopus. SPLASH is a new unifying approach to genomic analysis that enables an expansive scope of discovery without metadata or references.

**One-sentence summary:** SPLASH is a unifying, statistically driven approach to biological discovery from raw sequencing data, bypassing alignment.

## Introduction

Genomics is now foundational to biology, ecology and medicine, and as sequencing databases grow, so too does the opportunity to leverage them for discovery. How can this data best be analyzed to reveal regulation and function?

Traditionally, bioinformatic pipelines start by assigning genomic positions to reads via alignment to a reference genome. While alignment has served reasonably well for decades, it has many limitations. For less studied organisms, references are often incomplete and or partially misassembled; and for many organisms, there is no reference. Reliance on references risks missing sequences that are not present in the reference, a major problem even in the intensely studied human genome: for example, genetic variants associated with under-studied populations are poorly represented in current references^1^, undermining the impact of genomic studies and exasperating health disparities. Some diseases such as cancer are almost defined by their changes from the reference, and vary even within a single tumor, so reference-based approaches are inaccurate approximations of the underlying biology. Finally, the enormous diversity of viral and microbial genomes, and their constant adaptation^2, 3^ makes it infeasible to define a complete set of references. Practically, alignment to references is computationally slow, limiting the scale of genomic inference.

Another challenge for reference-based methods is dealing with repetitive elements: aligners are forced to assign mapping positions to reads that simply do not have a unique alignment. Repetitive regions comprise ∼54% of the human genome^4^ and may be important to analyze, but are often just ignored because alignment is difficult or impossible. Further, many genes have sequence-similar paralogs, which pose similar alignment issues.

Testing statistical hypotheses after alignment is also problematic. For reads that do map, the complex algorithms underlying sequence alignment, which may have ad hoc and hidden assumptions, are difficult to model; thus statistical statements about differences (from SNPs to splicing) are difficult if not impossible to obtain in closed form. Permutation-based methods do not solve this problem: they are capable of giving 10-fold underestimates of the false discovery rate (FDR)^5, 6^. Even simple tasks such as calling allele-specific expression in well-studied genomes are fraught with statistical imprecision introduced during the alignment step^7^.

As an alternative paradigm, we considered if there could be a shared formulation that would allow signals of biological interest to be detected with statistical methods, directly from sequencing data (without a reference genome), unifying seemingly disparate sets of biological questions (currently addressed with custom and distinct bioinformatic approaches). For example: does a set of patients harbor distinct viral strains? Are alternative mRNA transcripts specific to distinct cell types or conditions? Does RNA expression and processing differ with geography or season? Notice that the answer to each involves studying how sequences vary within a set of samples. As we show, this idea can be made formal, and answers can be obtained with an approach that uses references as an interpretive guide rather than a critical step in statistical inference.

Here, we introduce a simple unifying paradigm for genomic analysis: SPLASH (**S**tatistically **P**rimary a**L**ignment **A**gnostic **S**equence **H**oming). SPLASH is applied directly to sequencing reads without the use of a reference genome. It relies on a simple formalization of sequence variation (short stretches of varying sequences, “targets”, adjacent to short stretches of a constant sequence, “anchors”) that captures many biologically important events. The distribution of anchor-target pairs are cataloged and the differences in their distribution between the samples is tested statistically at a very early stage in the pipeline. This simultaneously detects variation of myriad forms: single-nucleotide polymorphisms, alternative exon splicing, DNA rearrangement or transposition, and more. With today’s paradigm, a separate algorithm would be required to detect each type of variation. Instead, with SPLASH, if the user is interested in specific types of variation they can focus on that with post-hoc processing and analysis; alternatively, SPLASH may – and in many cases does – expand horizons to unexpected types of variation. In the results below, we demonstrate a snapshot of SPLASH’s wide possibilities for discovery, first in human samples: encompassing viral strain variation, alternative isoforms in single cells, B and T cell antigen receptor diversity. We also show that SPLASH is easily applied to less studied organisms: in lemur, to find antigen receptor diversity; in octopus, to detect tissue-specific splicing, processing and expression; and in eelgrass, to uncover allelic variation, splicing, and diatom association. We believe this demonstrates SPLASH’s potential to discover meaningful sequence variation, with or without the aid of reference genomes, across many biological questions and organisms.

## Results

### SPLASH is a *k*-mer based, statistics-first approach to identify sample-dependent sequence variation

The fundamental concept of SPLASH is anchors and targets (Figure 1A). An “anchor” is simply any particular *k*-mer of sequence in a read (*k* = 27 by default, but adjustable). Every *k*-mer a fixed offset downstream (*R*, which may be zero) is called a “target”. Note that targets are always defined relative to an anchor, and a given anchor may have multiple associated targets. Anchors with more than one target can report on nearly all sequence variation of interest: the simplest case is where two targets differ at a single position, as in mutations, allelic differences or RNA editing. It also can capture insertions and deletions, alternative splicing and isoform usage, mobility of transposons, gene rearrangements and fusions, antibody/T cell receptor somatic variation, and more.

**Figure 1.**
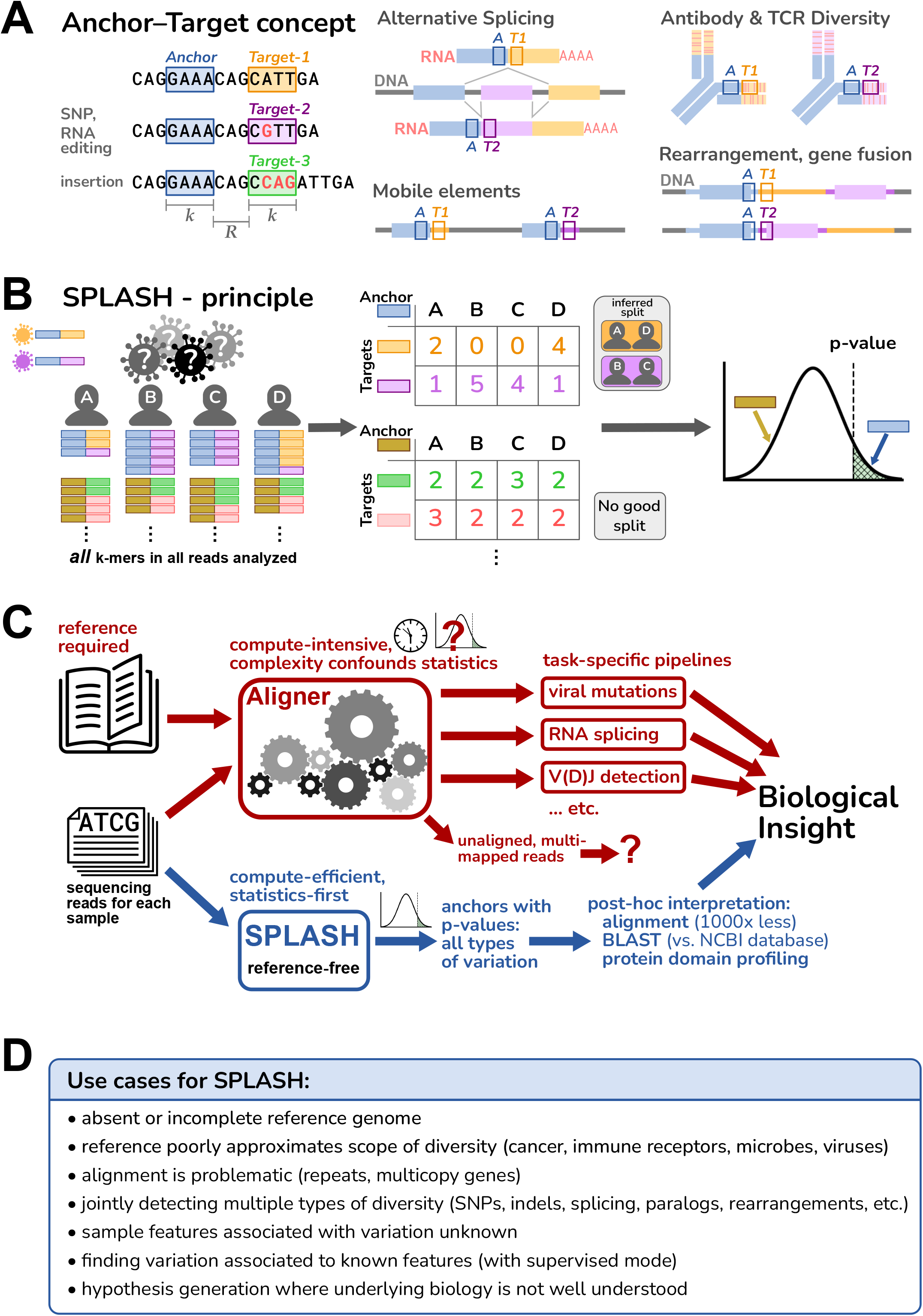
Overview of SPLASH. **A.** An anchor is a sequence of length *k* (*k*-mer) in a read; its target is defined as the *k*-mer that follows it after a fixed offset of length *R*. An anchor may occur with different targets, and this can capture many types of variation, ranging from single-nucleotide changes to alternative exon splicing; other examples are depicted schematically, with anchor as a blue box and targets as orange or violet boxes. **B.** SPLASH compiles a table for each anchor, where the columns are samples and the rows are targets, and entries are the respective occurrence counts. In its default unsupervised mode, SPLASH iterates over multiple random splits of the samples, calculating a test statistic that measures the deviations between each sample’s target distribution and the average target distribution over all samples, searching for the most discriminating split. For the best split identified, SPLASH reports a *p*-value bound (which is non-asymptotic, controlling Type I error for any number of observations). Anchors with low *p*-values have target distributions that vary significantly between samples. **C.** Standard alignment-based pipelines are limited by the need for a reference genome, are compute-intensive, and difficult to model statistically due their complexity. SPLASH was designed to detect variation directly from raw reads with rigorous statistics, is very compute-efficient, and can detect many kinds of variation at once. **D.** Some considerations of when SPLASH may be most useful, which reflect its design characteristics.

The fundamental goal of SPLASH is to detect variation among a set of sequencing samples, and it requires only a FASTQ file for each sample. The definition of a “sample” is dictated by the biological question: each sample could be from different cells, different tissues, different individuals, or different mixed populations (e.g. metagenomics). Samples might differ by conditions as well – different times or treatments, and also in other features (cell-type, geographic location, etc.); we refer to these generically as “metadata”.

The principle of SPLASH is depicted in Figure 1B (and discussed in detail in Note S2 and Figure S1). Conceptually, SPLASH steps through all positions in all reads of all samples, recording all anchor-target pairs. To save compute time, SPLASH can run in a mode that only analyzes anchors at a subset of read positions, e.g., every fifth position (as used in this work). SPLASH compiles a separate counts table for each anchor, with a column for each sample and row for each target; each entry is the counts of a given target in a given sample; classically, this is a contingency table. Compiling these tables requires only a single pass through the raw sequences and doesn’t involve alignment to a reference, so it is computationally efficient. Importantly, we have developed a highly flexible test statistic that captures the desire to find relatively discrete groups of samples with differing variation, and that controls false positives even for low numbers of observations. This statistic admits a rigorous closed form *p*-value bound, which is thus fast to compute (unlike resampling methods). For each anchor, SPLASH calculates a *p*-value bound for the null hypothesis that the observed target frequencies in the samples all come from the same distribution, i.e., that there is no underlying variation of targets between the samples. A low *p*-value for an anchor implies that the samples differ significantly in which targets they contain.

We emphasize that SPLASH does *not* need predefined sample groups (though it can also operate in such a “supervised” mode). Indeed, for all the results presented here, we used SPLASH in its default unsupervised mode. For each anchor, SPLASH tries many random splits of the samples and the targets, retains the one which minimizes the *p*-value, and reports the corresponding effect size for this grouping (next paragraph). Although perhaps surprising, this random process is designed to and does detect the presence of strongly patterned target variation among the samples, if it exists (Figure S3, Note S3).

SPLASH also calculates for each anchor an “effect size” that ranges from 0 to 1, with 0 meaning that the target distribution is the same between the two groups, and 1 meaning that the targets found between the two groups are disjoint. Effect size does not require that target distributions of all samples within a group are similar, just that they are different from the other group; thus, effect size can be high even when there are more than two natural groups. Anchors with effect sizes closer to 1 have target variation that is more discrete across the samples, and thus are more amenable to biological investigation.

To aid in biological interpretation it can be useful to have longer sequence context than just the anchor and target. Thus, SPLASH also can generate a “consensus” for each anchor, in each sample. Consensuses are longer sequences (up to the length of a read) assembled from the raw reads of a given sample, looking at every occurrence of an anchor and extending its sequence base by base as long as the reads show a consensus (see Methods). Mapping consensuses to protein sequences can identify protein domains – protein domain profiling – a powerful, reference-free attribution of biological function. Consensuses can also be aligned to sequence databases or reference genomes; aligning only the consensus sequences for significant anchors reduces the typical computational load by 1000-fold compared to usual approaches that align all reads.

Figure 1C diagrams some of the differences between traditional alignment-first approaches and SPLASH; Figure 1D outlines interesting use cases for SPLASH. These guided us in our initial explorations with SPLASH, which are described below. The datasets analyzed were simply the first ones we chose, and may not be the best or most informative. While SPLASH has some adjustable parameters, we did not attempt to “tune” these (indeed, SPLASH seems robust to a range of parameters, Note S3); all analyses were run with the same settings, in unsupervised mode (blind to metadata). Despite this, we found that SPLASH performed well across a variety of datasets, and in all cases found significant patterns of sequence variation (*q*-values for anchors, and binomial *p*-values quantitating the visually evident spreads in plots, are given in the Note S7).

### SPLASH identifies strain-defining and other mutations in SARS-CoV-2 *de novo*

Viral genomes have high mutation rates, including quasispecies^2^. For example, the emergence of SARS-CoV-2 was followed by multiple surges caused by variant strains, over the course of just two years. This is an ideal setting for the application of SPLASH: viruses are always mutating, but out of a sea of mutations, scientists, clinicians and public health officials want to identify those showing consistent variation between patient groups.

We applied SPLASH in unsupervised mode to two SARS-CoV-2 datasets, both viral amplicon Illumina sequencing of nasopharyngeal swabs from infection-positive patients, taken from times when the dominant viral strains were Delta or Omicron. The samples from South Africa^8^ (Nov to Dec-2021, 70 samples) represent the rapid rise of Omicron during its first outbreak. The samples from France^9^ (Dec-2021 to Feb-2022, 106 samples) represent cases of co-infection by more than one strain; the authors of the original study provided as metadata the assignment of the primary and secondary viral strains for each case (though not used by SPLASH).

SPLASH finds many significant anchors with low corrected *p*-values (<0.05) and high effect-sizes (>0.5), directly from sequencing reads (250 for South Africa dataset, 252 for France). High effect sizes are expected for anchors that partition samples by strain. To test if SPLASH recovers strain-defining and other variation, we examined the subset of significant anchors that perfectly map to a reference strain (Original, Delta, Omicron BA.1 or BA.2; defining mutations taken from CoVariants.org^10^), and call an anchor “strain-defining” if it has at least two targets with >5% abundance, at least one of which perfectly matches to a reference strain; by definition the other target is distinct.

We compare to a control set of anchors, those that are most abundant across all the reads; this tests that SPLASH’s findings are not solely due to sequence abundance. In the South Africa dataset, 98% (126/128) of SPLASH anchors that mapped perfectly were strain-defining, compared to 7/201(3.5%) in the control set (hypergeometric *p*-value <1.7E-79). In the France dataset, 100% (39/39) of SPLASH anchors were strain-defining, compared to 8.4% (21/250) of the control anchors (hypergeometric *p*-value <2.6E-33). Nearly all the control anchors have only a single abundant target (with the remaining low abundance targets representing either private mutations or sequencing errors). Thus SPLASH, though blind to strain reference sequences and sample metadata, detects strain differences with high precision.

Figure 2A-C shows exemplary strain-defining mutations identified by SPLASH; we focus on the Spike protein (S gene), where the majority of the mutations were located. Figure 2A shows an anchor that distinguishes at the major lineage level: one target has no mutations (relative to the Original SARS-CoV-2 reference) and is consistent with Delta; the other target has the mutation K417N, found in all Omicron strains (but not in Delta or Original). Target abundances across samples are consistent with the strain assignment metadata. Figure 2B shows an anchor that discriminates sub-lineages: one target has no mutation, consistent with Delta; the second target has the 3-nt deletion NL211I and the 9-nt insertion R214REPE which are Omicron BA.1 specific; the third target has the mutation V213G which is Omicron BA.2 specific. Figure 2C shows an anchor that detects mutations not in our references. One target has a pair of mutations, N679K and P681H, found in all Omicron strains. The other targets all have S:P681R, a Delta-specific mutation, but two targets additionally encode Q677H (by different mutations). Mutations at Q677 may provide a selective advantage: they have arisen independently multiple times on different lineages^11, 12^, and Q677H in several strain backgrounds enhanced infectivity, syncytia formation, and resistance to neutralizing antibodies in pseudotype assay 13.

**Figure 2.**
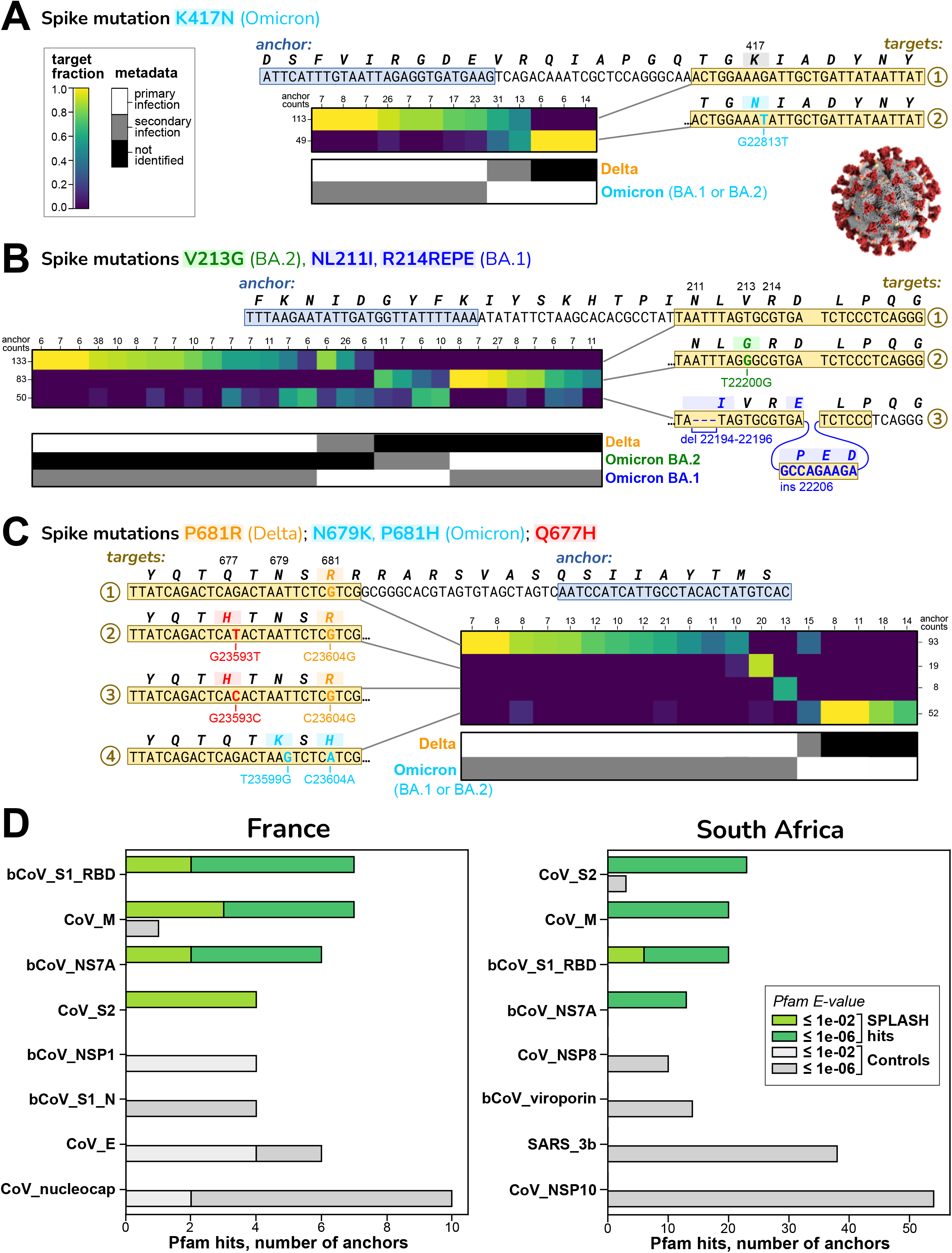
SPLASH identifies strain-defining and other variation in SARS-Cov2. In A-C, sets of targets that distinguish between SARS-CoV-2 strains are shown; all are in the spike protein (S) gene. Each heatmap has columns for different patients and rows for different targets; the coloring indicates the fraction of the given target observed in the given patient. Summary anchor counts are given for rows and columns, simply to give context of how many observations are summarized. Also shown is a map of the categorical metadata of what strains (primary and secondary) were identified in each patient in the original study; this data was not used by SPLASH, but there is evident agreement between the expression in the heatmap and the strain assignment in the metadata. We give binomial *p*-values to quantify the distinctions in the plots (per Note S7). **A.** Mutation K417N, identified by SPLASH in target 2, distinguishes at the major strain level: it is not found in Delta but is in all Omicron (both BA.1 and BA.2 sub-strains). Patients classified as Delta all express target 1; two patients co-infected with Delta and Omicron show both targets. (*p* = 6.4E-07) **B.** An anchor with three targets identified by SPLASH distinguish at the sub-strain level: target 1 with no mutations matches Delta, target 2 with V213G is specific for BA.2, and target 3 has both a deletion mutation (NL211I) and insertion mutation (R214REPE) characteristic of BA.1. Target 1 and 2 associate inversely with Delta and BA.2; target 3 is more mixed, due to all samples with this anchor expressing some level of BA.1. (*p* = 1.0E-13) **C.** An anchor with four main targets identifies mutations that are not associated to a specific strain: targets 2 and 3 encode Q677H (with different mutations) together with the Delta-specific mutation P681R, and each target is predominant in a different patient. Target 1 has only P681R and lacks Q677H. Target 4 has the Omicron-specific mutations N679K and P681H. SPLASH can identify complicated mutation patterns. (*p* = 4.9E-12) **D.** Protein domain profiling in SARS-CoV-2. The top four protein domain types found by Pfam for translated extended sequences for SPLASH significant anchors (green bars) and Control anchors (highest abundance but generally without significant *p*-values; gray bars) are shown. S1 Receptor binding domain (RBD) and S2 domain, known to be under strong selective pressure, show high variation by SPLASH in both datasets. Other abbreviations used in the Pfam short-names are: bCoV = beta-coronavirus; CoV = coronavirus, nucleocap = nucleocapsid N = N-terminal domain, SARS = Severe acute respiratory syndrome coronavirus; M, NS7A, NSP1, NSP8, 3b, NSP10 are viral protein names.

SPLASH results can also be analyzed completely without a reference genome by examining the coding potential of its nominated sequences.The anchor sequence is extended by a consensus approach, translated *in silico* to amino acid sequences (in all six reading frames) and matched against a database of protein domain models such as Pfam^14^; the best Pfam hit is assigned to that anchor. Protein profiles that are more frequently associated with significant SPLASH anchors, compared to control anchors, are candidates for proteins with important patterns of variation. SPLASH protein profiles for SARS-CoV-2 were overall extremely different from controls (chi-squared test *p*-values: France <3.7E-7, South Africa <2.5E-39) (Figure 2D). The top four SPLASH protein domains in both datasets were beta-coronavirus receptor-binding domain (RBD; within S1 region of spike protein), coronavirus S2 domain (within spike protein), coronavirus M (matrix/membrane) protein, and coronavirus ORF7a protein. By contrast, the top four control domains for each dataset were completely different from these, and also different between datasets. The SPLASH protein domains are significantly enriched when considered individually as well; for example, in the South Africa dataset for spike S2 domain: 23 SPLASH vs 3 control hits, *p* = 2.9E-6 (corrected hypergeometric *p*-value). The spike protein is the major site of antigenic variation in coronaviruses, as it is a principal focus of the immune response; the RBD is well known as a target for natural and therapeutic neutralizing antibodies^15^, but in addition about 50% of natural anti-spike antibodies are directed against the S2 domain^16^.

We also carried out SPLASH protein domain profiling on an unrelated virus, rotavirus^17^ (102 samples). The domains enriched over controls were rotavirus VP3 and NSP3 proteins (Figure S2A). These two proteins have roles in blocking host innate immunity^18^. Thus, variation in viral protein domains interacting with the immune system may be a recurring theme in SPLASH protein profiling of viral strains.

In summary, SPLASH finds patterns of variation in SARS-CoV-2, including those characteristic for strains, without requiring reference sequences or metadata (though these obviously help with interpretation); the methodology should be generally applicable to other viruses. More broadly, since SPLASH analyzes variation without any predetermined reference or template, it may be useful in surveillance for new strains or even completely new pathogens, and to cluster patients directly from raw sequencing samples.

### SPLASH identifies regulated expression of paralogs and HLA in single cell RNA-seq

Today’s approaches to single cell bioinformatics require references and custom bioinformatic pipelines that struggle to quantify the extent of (post-)transcriptional regulation of gene expression which includes splicing and highly similar sequence variants, such as paralogs. We tested if SPLASH can unify discovery in the realm of single cell sequencing generated with the Smart-Seq2 (SS2) protocol^19^, which provides broad transcript coverage.

Our first testbed was human macrophage and capillary cells from the Human Lung Cell Atlas^20^. We previously established that these cell types have regulated alternative splicing in MYL6, a light chain subunit of myosin motor protein, so that serves as a positive control^21^. As expected, among SPLASH’s most significant anchors are ones reporting this MYL6 alternative splicing, splicing in or out of a cassette exon (Figure S4A). Surprisingly, other SPLASH top anchors also involved different myosin light chain subunits, MYL12A and MYL12B. These two genes are paralogs with highly similar coding regions (95% nucleotide, 98% amino acid identity for human).

Nevertheless, SPLASH finds targets that specifically distinguish them. They show that macrophages express more MYL12A, while capillary cells express more MYL12B (Figure 3A). This same pattern is observed in two different individuals. Not much is known about these two genes, but they are known to show differential expression in rat tissues^22^, and there is evolutionary conservation in mammals, birds, and reptiles of two adjacent MYL12 paralogs within a syntenic region (e.g. human; rat^22^; *Gallus gallus*, NCBI Gene IDs 396284 and 770011; *Chelonia mydas*, Gene IDs 102938771 and 102937279). Besides the small number of amino acid differences between the paralogs, there may also be an important functional role for nucleotide sequence differences, as has been demonstrated for another pair of highly similar cytoskeletal paralogs, beta- and gamma-actin^23^.

**Figure 3.**
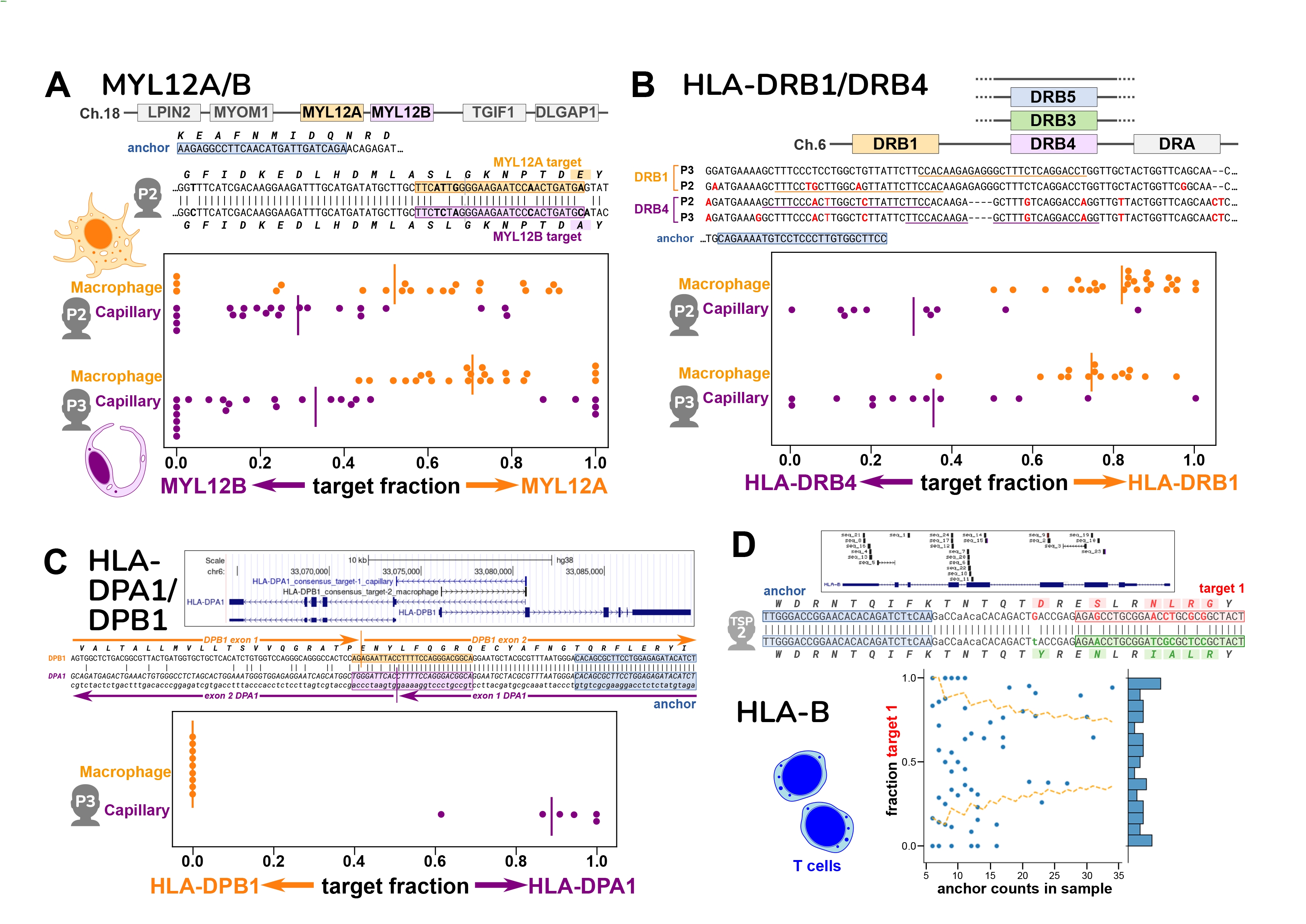
Cell-type specific expression of paralogs and HLA from single-cell data. Figures A-C show spread plots, with each dot representing the relative isoform expression in a single cell; a bar marks the average relative expression across all the cells. **A.** Human MYL12A and MYL12B lie adjacent on chromosome 18; this region is syntenic in mammals, chickens, and reptiles. The two genes are very similar in the coding region, as seen in the sequence alignment that also shows the locations of the anchor and two targets for individual P2 (anchor is distinct for P3). Macrophages express relatively more MYL12A and capillary cells more MYL12B, based on target fractions; this result is consistent in two different individuals. **B.** The HLA-DRB locus occurs as several different haplotypes, all of which contain a DRB1 gene but differ in presence of a paralog (DRB3, DRB4, DRB5, or none; hg38 reference has DRB5). The anchor and targets lie in the somewhat similar 3’ untranslated regions of DRB1 and DRB4; however the two individuals have distinct alleles at both DRB1 and DRB4. Macrophages express mainly DRB1 and capillary cells mainly DRB4, based on target fractions. **C.** The HLA-DPA1 and HLA-DPB1 genes overlap in a head-to-head arrangement as shown in the UCSC Genome Browser plot, which also shows the BLAT alignment of the consensus sequences for the DPA1 and DPB1 anchors. These lie on opposite strands of the genome. This is also depicted in the alignment of the two consensuses (which represent mRNA); the targets are best assigned to opposite strands, and the location of splice junctions corroborates the strandedness. An anchor reporting simultaneously on DPA1 and DPB1 was only found for individual P3; its targets show that macrophages exclusively express DPB1, while capillary cells express DPA1. **D.** T cells from one individual showed a large number of anchors across the polymorphic HLA-B gene, as depicted in the UCSC Browser plot. We investigated one HLA-B anchor, which lies in exon 2. The hg38 reference is the B*07:02 allele, whereas alleles of this individual match best by BLAST to B*08 and B*51 (consensus 1 and 2). In the alignment, differences from hg38 in the anchor and lookahead region are shown in lowercase. Individual T cells show a wide range of expression ratios between the two HLA-B alleles (different target 1 fractions), as shown in the scatterplot. The yellow lines mark a 98% confidence interval for a distribution based on the population average (confidence depends on observation depth = anchor counts); the observed pattern differs (binomial *p* = 1.73E-25), and some cells express almost exclusively one allele over the other.

In the same data, SPLASH finds cell type-specific expression of genes in the major histocompatibility complex (MHC), known as HLA in humans. HLA is the most polymorphic region in the human genome, with the most known disease associations; the polymorphism of HLA class I and class II proteins is intimately tied to their function in antigen presentation for adaptive immunity^24^. Due to the high levels of polymorphism, the region is challenging for representation in a reference genome and for read-mapping pipelines. SPLASH identifies an anchor and targets that distinguish between the highly similar 3’ untranslated regions (UTRs) of HLA-DRB1 and HLA-DRB4 (class II beta-chains) (Figure 3B). Macrophages express mainly DRB1, while capillary cells mainly express DRB4. The same pattern is found in two different individuals, who carry different alleles at DRB1 and DRB4 (as seen in the alignment). Macrophages are “professional” antigen-presenting cells and constitutively express HLA class II. Not all endothelial cells express class II *in vivo*, however most human capillary cells do^25^; endothelial MHC expression may be strongly cytokine-dependent^26^. Thus, class II expression in macrophages and capillary cells is likely to be regulated differently. Five major haplotypes have been identified at the HLA-DRB locus: all contain a DRB1 gene, but some contain an additional functional paralog, either DRB3, DRB4, or DRB5^27^. DRB1 is extremely polymorphic (3516 alleles, in March 2023); the paralogs somewhat less so, e.g. DRB4 (236 alleles)^28^.

In one individual (P3), SPLASH also finds a remarkable anchor and two targets that report on a pair of class II genes, HLA-DPA1 and HLA-DPB1 (class II alpha and beta chains, respectively), which are unique among HLA genes in being organized head-to-head and transcribed in opposite directions. The anchor lies in sequence shared by DPB1 and a specific isoform of DPA1, while the targets lie in exons exclusive to each; the extended sequences provided by SPLASH consensuses confirm directionality as they bridge splice junctions. Macrophages in this individual express exclusively DPB1, while capillary cells express mainly DPA1 (Figure 3C). This pattern may be haplotype-specific, as we did not find a similar anchor for another individual (P2).

A final example is SPLASH’s detection of allele-specific expression of HLA-B in human T cells (from a different dataset, see next section). The class I gene HLA-B is the most polymorphic of all HLA genes (9274 alleles)^28^, and HLA-B is the gene with the most anchors found by SPLASH in human T cells (Figure 3D). Since these T cells all come from one individual, this indicates substantial variation in expression of this individual’s two HLA-B alleles at the single-cell level, as can be seen in the scatterplot (Figure 3D). Different T cells express a wide range of ratios of the two alleles, some cells expressing both alleles, but others expressing almost entirely one allele or the other (well outside the 98% confidence interval of what is expected by the average ratio). This is in keeping with a preprint that found allele-specific expression of HLA class I genes in normal breast epithelial cells, as well as in multiple myeloma cells^29^; other single-cell studies have found changes in allele-specific expression during T cell activation^30^, and identified HLA expression quantitative trait loci^31^.

Overall, SPLASH finds disparate biological features producing regulated variation at the single cell level: paralogs, splicing, and alleles, especially within the medically important and polymorphic HLA locus. This is just a small glimpse into the complexity of HLA haplotype- and cell type-specific expression patterns; it raises the possibility that disease-related HLA alleles might be expressed differently in key cell types compared to other alleles.

### SPLASH identifies B and T cell receptor diversity in human and lemur single cell RNA-seq

Adaptive immune receptors for B cells (immunoglobulin or Ig), and T cells (T cell receptor or TCR) are generated combinatorially through V(D)J recombination, and Ig is further diversified through somatic hypermutation. Rearranged sequences are absent from germline reference genomes and cannot be cataloged comprehensively due to their huge potential diversity, empirically estimated at 10^10^-10^11^ for Ig heavy chains^32^.

These genomic loci currently require manual curation due to their complexity and repetitive structure, so only a few species have high quality annotations. Existing methods to assign V(D)J rearrangements in single cells^33^ depend on annotations and so may have blindspots. Since SPLASH is designed to identify sequence diversity without need for references, we predicted that it would identify adaptive immune receptors *de novo*.

We ran SPLASH on 50 naive human B cells from peripheral blood of one individual, and separately on 128 CD4+ human T cells of another individual, taken from Tabula Sapiens, a large multi-organ dataset^34^. As a first pass, requiring no reference or alignment at all, protein profiling found that the most frequent domains matching to SPLASH anchors in B cells were Ig V-set and C1-set (variable-like and constant-like domains); these two domains were also matches in T cells (attributable to TCR) (Figure 4A). Mapping transcript gene-names to SPLASH anchors gives a similar picture: Ig light chain genes (both kappa and lambda) were strongly predominant for B cells; HLA-B (discussed above) and TCR genes (both alpha and beta) were most prominent for T cells; and these are not found among the control anchors which are up to 10-fold more abundant (Figure 4A). Significant anchors were also found in Ig heavy chains (shown in lemur, below), though fewer anchors than in Ig light chains. Ig/TCR anchors characteristically have a high diversity of targets (“entropy”, a measure reported by SPLASH), and could be identified on that basis rather than requiring reference mapping. This is expected for clonally diverse receptors, and is very evident in the clonotypic pattern (each cell expressing only its specific target) seen in the heatmaps (Figure 4B).

**Figure 4.**
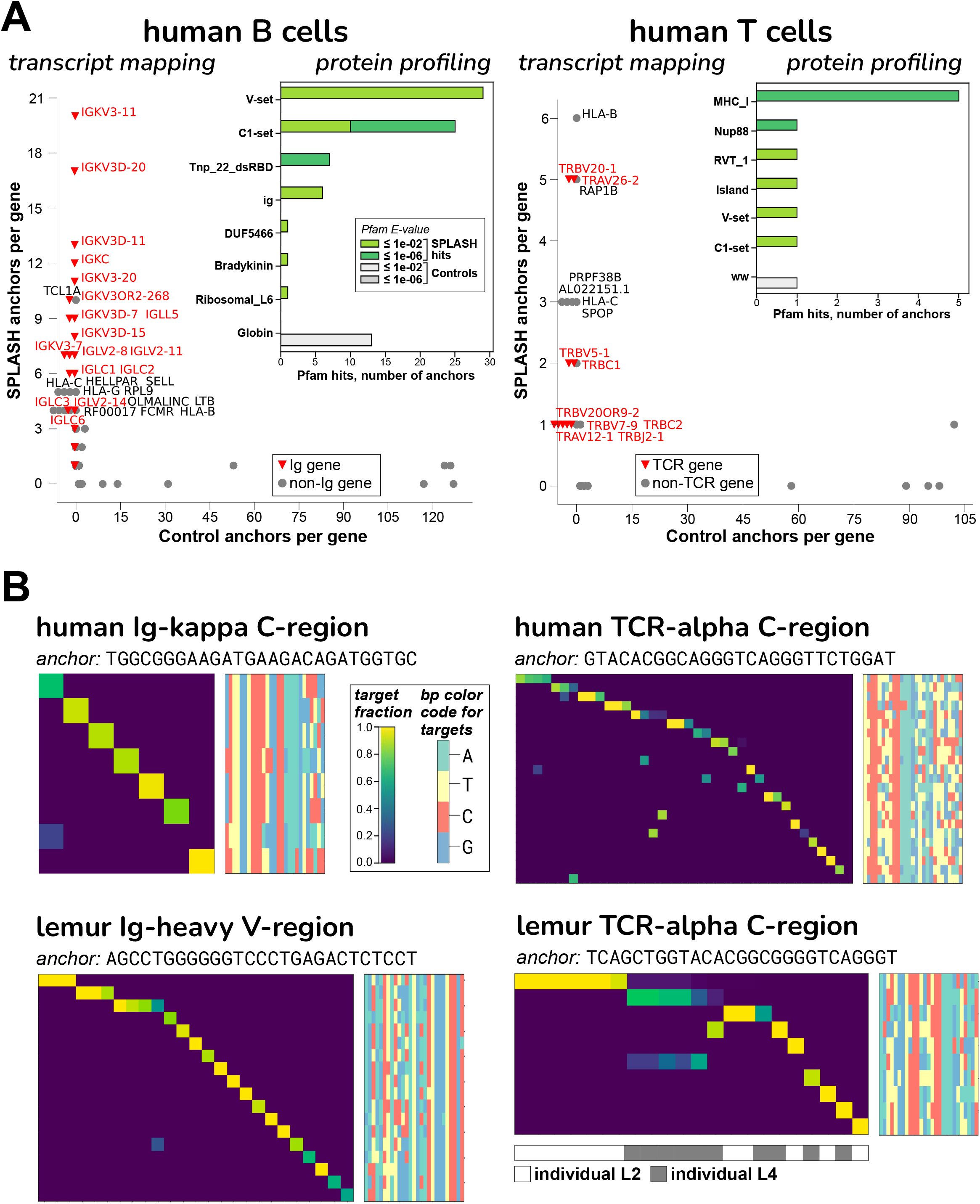
B and T cell receptor diversity from human and lemur single cell data. **A.** The “transcript mapping” plots show the number of anchors that align to a given gene name, for SPLASH on y-axis and Controls on x-axis, with immune receptor genes highlighted in red. For B cells, Ig genes (kappa = IGK and lambda = IGL) are by far the most predominant mapping among SPLASH anchors, but are not found at all in Control anchors (those with the highest counts). For T cells, TCR genes (alpha = TRA, beta = TRB) predominate, and are also not found in Controls. The inset histograms show that, in B cells and T cells, immunoglobulin-type “V-set” and “C1-set” are among the top protein domain annotations identified by Pfam on anchor consensuses, without using a reference genome. Mobile element activity is suggested by Pfam domains Tnp_22_dsRBD (“L1 transposable element dsRBD-like domain”) in B cells and RVT_1 (“Reverse transcriptase”) in T cells. **B.** Targets associated with Ig/TCR anchors are clonotypically expressed, in both human and lemur: heatmaps show that most targets (rows) are expressed only in a single cell (columns). The anchors shown have among the highest target entropy scores for their gene type (Figure S5B illustrates an extreme example with 97 targets, for lemur Ig-lambda.) Target sequences are shown as bp color-maps (rows are targets, matching those in the heatmap; columns are bp positions, colored by base), giving a quick visualization of the sequence diversity. For lemur, NKT cells were studied, and show significant shared TCR usage – see top two rows; interestingly, the shared target sequence is different in the two individuals. There is similar sharing for lemur TCR-gamma and -beta chains (Figure S5C and D).

To showcase application to non-model organisms, we also ran SPLASH on mouse lemur (*Microcebus murinus*). Mouse lemurs are primates that diverged from humans 60-75 million years ago, and have potential as a model organism^35^. The lemur reference genome is incompletely annotated, especially at loci such as Ig and TCR. While the human reference does not suffice to perform alignment-first analysis of mouse lemur transcriptomes, we find here that it is a reasonable approximation for interpreting SPLASH outputs; this may generalize, allowing SPLASH interpretation for organisms wherever a related, better-curated reference exists. From Tabula Microcebus, a multi-organ mouse lemur dataset^36^, we analyzed 111 natural killer T (NKT) cells from spleen and, separately, 289 B cells also from spleen. In both analyses, the cells came from two different individuals; for this reason, SPLASH also discovered numerous allelic differences between individuals, such as in COX2 (Figure S5A).

Our main focus was on adaptive immune receptors; similar to the human analyses, we found that SPLASH’s lemur anchors in B and NKT cells included Ig C1-set and V-set domains by protein profiling and Ig/TCR gene-names by transcript mapping (data not shown). As expected, SPLASH targets for lemur Ig heavy chain are predominantly clonotypically expressed (Figure 4B). Lemur NKT cells provide an interesting counterpoint. While there is some clonotypic diversity, it is striking that a number of cells share TCR-alpha sequences; notably, the shared target is different between the two individuals (bottom-right heatmap, top row vs. second row). We had selected NKT cells for analysis without foreknowledge of their properties. However, SPLASH has correctly characterized diversity in this case: it is known that an NKT subset expresses an “invariant” TCR-alpha chain with limited TCR-beta diversity, in humans and mice; NKT cells bridge between adaptive and innate immunity^37^. For Tabula Microcebus, NKT cells were operationally defined as co-expressing CD3E and KLRB1 (CD161)^36^; in this cell-type, SPLASH also finds shared usage in TCR-beta and TCR-gamma (Figure S5C and D).

To test if SPLASH finds diversity that may be missed by alignment-based methods, we analyzed a subset of 35 lemur B cells for which Ig light chain variable regions could not be assigned by the program BASIC^38^. BASIC uses curated human Ig sequences to assign V-D-J regions to Ig mRNAs; although it was able to assign the large majority of lemur B cells, there was a small subset for which it failed. Using a quite naive approach, we used SPLASH to find evidence of a light variable region in one of the 35 cells; we were able to reconstruct the full variable region from reads (Note S4). In two cells, there were hits to the surrogate light chain (IGLL1/IGLL5 or λ5), which associates with Ig heavy chain when there is not yet a rearranged light chain^39^. The additional yield from SPLASH was modest, but is proof-of-principle that SPLASH provides views on data distinct from traditional methods. In more recent work we have built on this capability (J. Salzman *et al.*, unpublished work).

### SPLASH applied to non-model organisms: octopus and eelgrass

To explore SPLASH’s generality, we applied it to two understudied organisms: octopus and eelgrass. Octopuses have the most complex sensory and nervous systems among invertebrates, and are unusual in having high levels of RNA editing^40^. The marine angiosperm *Zostera marina*, or eelgrass, is the most widely distributed seagrass, and its adaptation to varying conditions, especially in the face of climate change, is of great interest and only beginning to be explored at the genomic level^41, 42^.

We focused narrowly on anchors where no more than one of its targets mapped to the respective reference (Methods), that is, where reference mapping could not have detected variation, to showcase what SPLASH can uncover that would be missed by conventional approaches. However, that means that in these analyses we ignored any interesting findings of SPLASH that are consistent with the reference.

For octopus, we analyzed an RNA-Seq dataset of *Octopus bimaculoides*^43^, encompassing a variety of tissues from a single individual (N. Bellono, personal communication). We examined several anchors with high effect sizes and BLAST hits to the closely related species *Octopus sinensis* (Methods; Data S6). An interesting SPLASH anchor was found in *O. sinensis* myosin-VIIa, known as MYO7A in humans; mutations in MYO7A cause Usher syndrome, leading to deafness and blindness^44^.

Target 1 corresponds to the annotated first exon of *O. sinensis* myosin-VIIa, while target 2 represents a novel alternative first exon (not annotated in *O. sinensis*) that is expressed only in statocyst tissue (Figure 5A). Although neither target is part of the annotated *O. bimaculoides* myosin-VIIa gene, both target sequences are present shortly upstream of it in the genome; but the region containing the anchor is missing (Note S5). Incidentally, the *O. sinensis* myosin-VIIa gene also shows likely misassembly (Note S5). The statocyst-specific expression of an alternative first exon is intriguing given MYO7A’s association with Usher syndrome and deafness, as the octopus statocyst is a sensory organ for sound and balance^45, 46^, suggesting homologous gene function.

**Figure 5.**
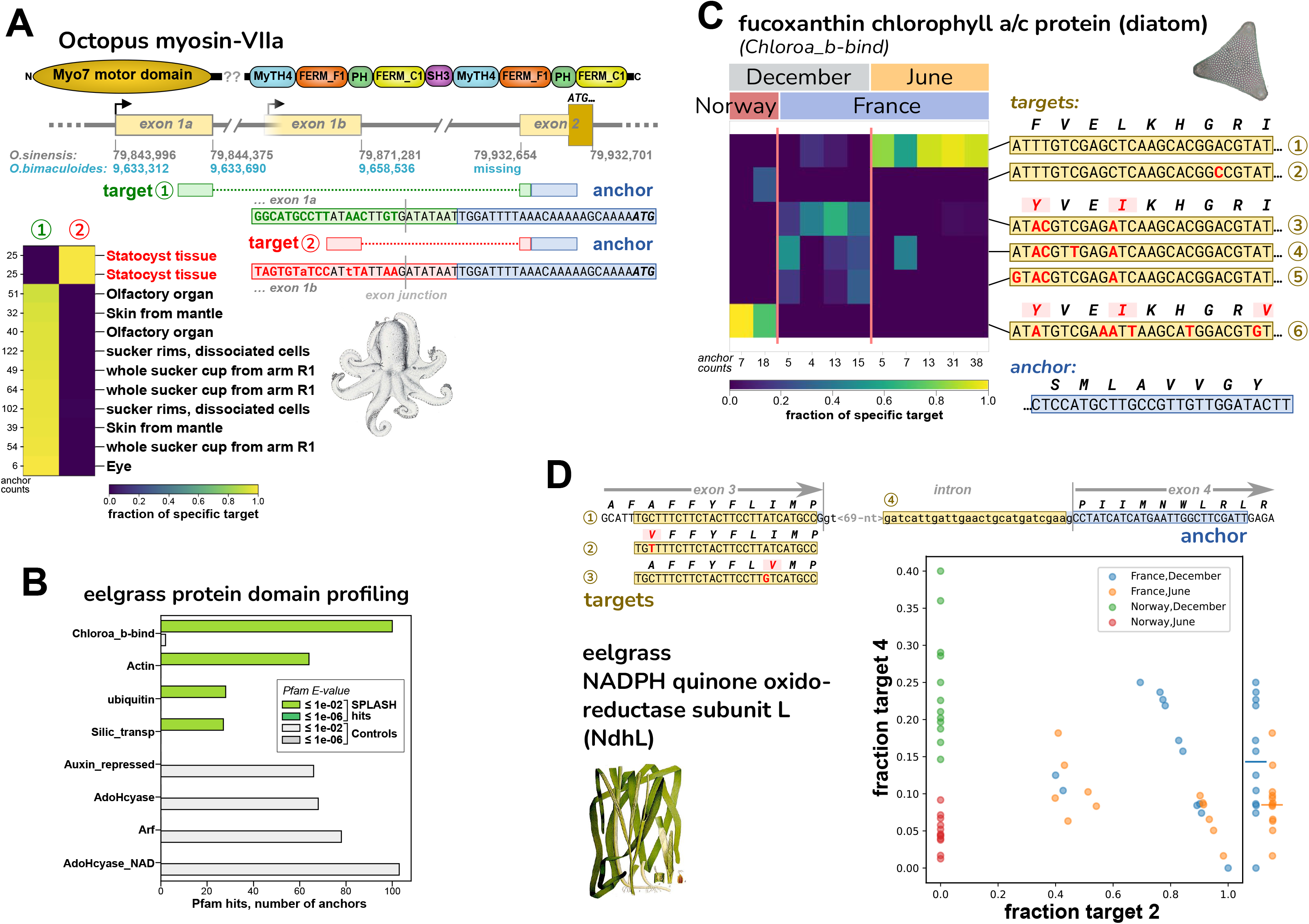
Discovery of regulated variation in non-model organisms: octopus and eelgrass. **A.** SPLASH identified alternative transcripts in the *O. bimaculoides* Myo-VIIa motor protein that are expressed mutually exclusively; the target 2 isoform is only found in statocyst, a sound-responsive tissue. The transcripts have different first exons, presumably due to alternative transcription start sites; the start codon lies in the shared second exon. The sequence of the anchor and exon 2 are missing from the current *O. bimaculoides* genome assembly, but are in the closely related *O. sinensis* genome. Sequences for targets 1 and 2 are in both genomes, but the statocyst-specific transcript is not annotated. The *O. sinensis* assembly has the Myo-VIIa gene in two inverted pieces (at the point marked by ‘??’ in protein domain schematic; Note S5). **B.** Two of the top domains identified from SPLASH anchors by protein domain profiling in eelgrass (*Z. marina*) samples are chlorophyll A-B binding protein (Chloroa_b-bind) and silicon transporter (Silic_transp); these derive from diatoms, based on BLAST protein alignment (Figure S7; Data S7). The other two top SPLASH domains, actin and ubiquitin, do not appear to derive from diatoms nor eelgrass, so may be from other epiphytic species. **C.** A Chloroa_b-bind anchor, more specifically identified by protein BLAST as a fucoxanthin chlorophyll a/c protein from diatoms (Figure S7C), has several targets that are differentially abundant: the most common target (top row) is mainly found in France/June samples; three targets that encode the same protein sequence (middle) are found in France/December samples; and one target (bottom row) is only found in Norway/December samples. **D.** An anchor from eelgrass, in the photosynthetic gene NdhL, has four major targets. Targets 1-3 are allelic coding variants (within a transmembrane region). Target 4 represents intron retention and would result in a shortened protein. The scatterplot shows that Norway samples of June vs. December (red vs. green) are perfectly segregated by the fraction of target 4 (intron retention). A similar but less marked trend is seen for France samples of June vs. December (yellow vs. blue) – at the right edge, fraction target 4 values are collapsed to one dimension, with averages marked by bars. The scatterplot also shows that Norway samples do not express target 2 at all (they express target 1 and 3, data not shown).

Other SPLASH anchor-targets did not map to the *O. bimaculoides* genome or transcriptome yet did BLAST to 3’ UTRs of annotated transcripts in *O. sinensis*, including carboxypeptidase D, Upf2, and netrin receptor/DCC (Figure S6, Data S6). For all three, each target is expressed exclusively in some tissues and not others. Although *O. bimaculoides* has annotated transcripts for these genes, in all three cases the 3’ UTR region is missing or likely incorrect in the *O. bimaculoides* genome assembly. For two of the genes, the target variation may be due to differential expression of alleles: a 13 nt deletion in carboxypeptidase D, and a short CAG repeat polymorphism in nonsense-mediated decay gene Upf2 (“regulator of nonsense transcripts 2”) (Figure S6A and B). For netrin receptor, involved in axon guidance and apoptosis, the variation SPLASH detects could be allelic but is also consistent with A-to-I RNA editing (Figure S6C). Our focus here on non-mapping anchor-targets excluded many more readily interpretable examples of regulated variation; for example, other evidence for RNA editing in numerous octopus genes (data not shown).

We also applied SPLASH to RNA-Seq data from eelgrass (*Zostera marina*), collected in two locations, Montpellier, France (Mediterranean climate) and Rovika, Norway (near-arctic climate), in two seasons (winter and summer), and during day and night^42^. Considering each anchor with its most abundant target, a large number (14,680, 5.7%) did not align to eelgrass references (Data S7). A high-level view of these is provided by protein profiling of SPLASH anchor-targets: the top hits were chlorophyll A-B binding protein domain, Actin, Ubiquitin, and Silicon transporter (Figure 5B). protein BLAST of some of the translated sequences finds that these have their best hits to a variety of organisms other than eelgrass, notably diatoms. This was a surprise to us, but in fact it has long been known that eelgrass is extensively colonized by epiphytes^47^, of which diatoms predominate^48^ and may provide as much as 71-83% of the primary production by the community^49^. For the most enriched protein domain, we investigated an anchor with high effect size (0.66); its consensus matches to “fucoxanthin chlorophyll a/c protein” (FCP) in several diatoms, for example, *Phaeodactylum tricornutum* (95% amino acid identity, 81% nucleotide identity; Figure S7C); given that the matches are imperfect, the true species of origin may not be in the NCBI database. FCPs function as light-harvesting antennae for photosynthesis^50^. This anchor has several targets (differing in their amino acid sequence), whose abundance varies by the sampling location and time of year: target 1 is strongly predominant in France in June; targets 3, 4, 5, which share the same amino acid sequence, together dominate in France in December; and target 6 is predominant in Norway in December (Figure 5C). These targets could represent different diatom species or intra-species allelic variants. The abundance of this anchor (irrespective of target) is lower in Night samples (Figure S7C), indicating circadian regulation of this diatom photosynthetic gene. Other anchors mapping to diatoms, such as ferredoxin and high mobility group box-containing protein, also show targets that segregate by location, France vs. Norway (Figure S7A and B).

One of the top anchors, although mapping to eelgrass reference, does report on novel variation. It is in the NdhL subunit of chloroplast NADPH dehydrogenase complex (NDH). Of its four most abundant target sequences, three are within exon 3 and are SNP coding variants; the fourth represents retention of the intron following exon 3 (Figure 5D). Intron retention alters the second transmembrane segment and terminates translation soon after (Note S5). The intron retention variant (target 4) is highest in winter (Figure 5D): for Norway, December (green) vs June (red) samples completely segregate by target 4 expression; the France samples show overlap, but on average December (blue) is higher than June (yellow). The scatterplot of Figure 5D also illustrates some other patterns: Norway samples do not express target 2 (instead they express target 1 and 3; data not shown); France samples have either a high fraction of target 2, or moderate (the latter also express target 1; data not shown). NDH is involved in cyclic electron transport^51^ and modulation of NDH function may affect photosynthetic efficiency and oxidative stress under varying light conditions^52^.

The above work with octopus and eelgrass are just early forays, but show that the SPLASH approach can discover novel regulated RNA splicing and isoforms, and bring to light allelic variation and communities of associated organisms. SPLASH’s statistics-first and reference-free methodology is an essential basis, which can be augmented by protein profiling and wide use of the sequence database beyond solely genomic references.

## Discussion

Genomic analysis today is performed with complex computer scientific workflows that are highly problem-specific. Here, we have shown that a single, simple statistics-first analytic framework unifies the approaches taken in disparate subfields of genomics and enables novel discovery missed by current workflows. We have achieved this by formalizing sample-dependent sequence variation in a single probabilistic model, and using this model to develop SPLASH. SPLASH frames prioritizes interpretable sequence variation as relations between sequence *k*-mers: anchors and targets; doing this required developing a new statistical test. While there is a history of using *k*-mers to detect biological variation, some statistical^53^, it was restricted to coarse-grained studies of genome composition.

Here, we provide a snapshot of SPLASH’s discoveries in very disparate subfields of genome science. When run on SARS-CoV-2 patient samples during the emergence of the Omicron variant, without strain metadata or reference genomes, SPLASH finds many anchors capturing strain defining mutations and emerging mutations, with good post-facto agreement with metadata. Using protein domain analysis, SPLASH finds that the spike protein is highly enriched for sequence variation, without using any reference sequences. This points to a broader potential for SPLASH in viral and other genomic surveillance, since emerging pathogens will likely have sequence variation missing from any reference.

In single-cell sequencing data, SPLASH is able to define differential expression between highly similar genes, including several different HLA genes and myosin light chains MYL12A and B. Like all the work presented here, SPLASH analysis is conducted in unsupervised mode; yet many of its significant anchors show cell-type regulation between macrophages and capillary endothelial cells. This testifies to the power of SPLASH’s unique statistical approach. In particular, HLA genes are highly polymorphic and traditionally difficult to analyze in genomics, yet have important associations to disease susceptibility. When applied to B and T cells of both human and mouse lemur, SPLASH automatically identifies antigen receptor genes, which are somatically assembled by V(D)J recombination, as exhibiting the most diverse variation. Lemur analysis was performed using only an approximate genomic reference (human) that diverged from lemur ∼60 million years ago. SPLASH might also be used to shed new light on other polymorphic loci including non-coding RNAs, e.g. spliceosomal variants.

To showcase SPLASH’s wide applicability, we used it on datasets from very different organisms: octopus and eelgrass. We narrowly restricted our analysis to anchor-targets that did not map to the respective genome assemblies. In octopus, we identified several tissue-regulated transcripts, in particular one in Myo-VIIa that is only expressed in statocyst. In the eelgrass dataset, SPLASH uncovered that many of the sequences come from epiphytic diatoms, and have variation that correlates with geography and season. A broader analysis, allowing mapping to the genome (as done in the human examples), would almost certainly find many more examples of regulated expression in these understudied organisms. This highlights the enormous potential in already existing datasets, and the need for tools like SPLASH to better explore them.

SPLASH should be of general interest to most genomic analyses. Users should be able to easily run SPLASH on their own samples (FASTQ files): we provide it as a containerized Nextflow pipeline to minimize installation issues, and the default parameters should work well (SPLASH is robust to a range of parameters, Note S3); it can even run on a laptop (Note S6). SPLASH will provide a large output file of significant anchors and targets. What next? First is to realize that these results are a large data reduction and distillation of the variation present in the samples. There are many ways that they can be used. The typical steps are to attach as much biological meaning to them as possible. If metadata is available, it can be correlated with anchor-targets generated in unsupervised mode; alternatively, one can use metadata to supervise SPLASH analysis. If a reference genome is available, SPLASH can use it to align anchors and targets and provide gene names. SPLASH provides a number of metrics, such as *p*-value, effect size, target entropy, average target similarity (Hamming or Levenshtein distance) which can be used to tailor the anchor list. Another useful avenue is BLAST of anchor-targets or consensus sequences against the NCBI databases (both as nucleotide, and protein after *in silico* translation); this can be very helpful especially when there is no reference genome or it is incomplete. Also useful is protein domain profiling with databases like Pfam. Ultimately, users will bring their own area expertise to bear in deciding how to best utilize SPLASH results.

Even in areas where there are existing pipelines, for example in differential alternative splicing, or antigen receptor identification, SPLASH provides a very different paradigm and may well give additional insights. And where there are no good references, it may be the only feasible option. SPLASH scales to allow discovery to keep pace with the ever-increasing sequence data from the world at large, in particular microbes and metagenomic communities; recent collaborative work has further increased the computational efficiency of SPLASH^54^. The SPLASH paradigm is expansive. It could be applied to a wide range of “omics” modalities, from DNA and protein sequencing to Hi-C and spatial transcriptomics, and more (see Note S1 “Generality of SPLASH”). The statistical ideas underlying SPLASH are also expansive: anchor-target pairs can be generalized to tensors, and higher-dimensional relations between anchors, targets, and samples can be studied; other functions for splitting and hashing targets and samples can be considered, to optimize statistical power^55^.

In summary, SPLASH illustrates the power of statistics-first genomic analysis, where references and annotations are only optionally used for *post-facto* interpretation. References are valuable for interpretation of sequence data; however, the filtering of data by alignment to references introduces statistical bias in quantification, blindspots and errors. SPLASH re-directs from the “reference-first” approach to “statistics-first”, performing statistical hypothesis tests on raw sequencing data. By this design, SPLASH is also highly efficient. SPLASH promises data-driven biological study with scope and power previously impossible.

### Limitations of the study

SPLASH can be applied to problems across diverse fields which are of great current importance (Note S1), including those previously discussed. Naturally, some problems are not directly amenable to SPLASH analysis as formulated here. The most clear are cases where quantification of sample-specific RNA or DNA abundance alone is desired (e.g., differential gene expression analysis). Additionally, SPLASH is currently unable to distinguish which biological mechanism underlies the called variation, and work in progress seeks to address this.

## Acknowledgements

We thank Elisabeth Meyer; Arjun Rustagi for extensive discussion and assistance in interpretation of viral biology; Roozbeh Dehghannasiri for selecting and collating data from HLCA/TSP/Tabula Microcebus analysis; the Tabula Microcebus Consortium (Camille Ezran and Hannah Frank) for data sharing and discussion of B and T cell receptor detection algorithms; Jessica Klein for extensive figure assistance; Michael Swift for discussion of B and T cell receptor detection algorithms; Julia Olivieri for feedback on presentation and detailed comments on the manuscript; Andy Fire for useful discussions and feedback on the manuscript; Robert Bierman for figure assistance and laptop timing; Sebastian Deorwicz and Marek Kokot for early access to code, and Aaron Straight, Catherine Blish, Peter Kim, Alex Starr and all members of the Salzman lab for feedback and comments.

## Author contributions

K.C. designed and implemented the pipeline. T.Z.B. designed, developed, and analyzed the statistics. G.H., P.L.W. provided SPLASH data analysis. I.N.Z. developed the protein domain analysis. J.S. designed and developed the statistics, and conceptualized and supervised the project. All authors analyzed data and wrote the manuscript.

## Data and materials availability

The code used in this work is available as a fully-containerized Nextflow pipeline^56^ at https://github.com/salzmanlab/nomad, commit 1b73949.

The human lung scRNA-seq data used here is accessible through the European Genome-phenome Archive (accession number: EGAS00001004344); FASTQ files from donor 1 and donor 2 were used. The FASTQ files for the Tabula Sapiens data (Smart-seq2) were downloaded from https://tabula-sapiens-portal.ds.czbiohub.org/; B cells were used from donor 1 (TSP1) and T cells from donor 2 (TSP2). The mouse lemur single-cell RNA-seq data used in this study was generated as part of the Tabula Microcebus consortium; the FASTQ files were downloaded from https://tabula-microcebus.ds.czbiohub.org. Viral data was downloaded from the NCBI: SARS-CoV-2 from France (SRP365166), SARS-CoV-2 from South Africa (SRP348159), and rotavirus (SRP328899).

The sample sheets used as pipeline input, including individual sample SRA accession numbers, for all analyses are uploaded to pipeline GitHub repository. Similarly, scripts to perform supplemental analysis can be found on the pipeline repository.

FASTP v0.23.2 was installed on 2/15/23 using bioconda. R-SPLASH v0.3.9 was installed on 2/23/23, commit 5dafdc8.

## Open-source figure attributions

Person graphic, by Tanguy Krl: https://thenounproject.com/icon/person-1218528 Virus graphic, by Nuttapon Pohnprompratahn: https://thenounproject.com/icon/virus-2198681.

Gears graphic, by Tresnatiq: https://thenounproject.com/icon/gears-1088494. Clock graphic, by sudan: https://thenounproject.com/icon/clock-5677937.

Book graphic, by Arjan: https://thenounproject.com/icon/open-book-1361747. SARS-CoV-2 virion, by Centers for Disease Control and Prevention (CDC): https://commons.wikimedia.org/wiki/File:SARS-CoV-2_without_background.png Octopus bimaculatus, by United States Fish Commission (1910): https://commons.wikimedia.org/wiki/File:Octopus_bimaculatus1.jpg

Diatom, by George Swann: https://commons.wikimedia.org/wiki/File:Diatom_4.png Zostera marina, by Carl Axel Magnus Lindman: https://commons.wikimedia.org/wiki/File:491_Zostera_marina.jpg

## Materials and Methods

### SPLASH overview

Full details of running SPLASH and its outputs can be found at https://github.com/salzman-lab/nomad. Briefly, it takes as inputs a set of FASTQ sequencing data files (one per sample). There are various parameter choices, but it should work well with the default settings. The standard output includes a table of anchors, targets, *p*-values, etc., a table of “consensus” sequences (see below), and a table of “element annotations” (see below). SPLASH can optionally also perform alignments with bowtie2 and STAR, to generate “genome annotations” and splice junction annotations (see below).

### SPLASH anchor preprocessing and parameter choices

Anchors and targets are defined as contiguous subsequences of length *k* positioned at a distance *R* = max(0, (*L* - 2 * *k*) / 2) apart, where *L* is the length of the first read processed in the dataset, and *R* is rounded to the nearest integer. If *L* = 100 and *k* = 27, then *R* = 23. We note that for a *fixed* number of anchor-target pairs, under alternatives such as differential exon skipping, larger choices of *R* have provably higher power than smaller choices, following the style of analysis in^57^. *k* = 27 is typically long enough to be assigned a unique position in a genome while having a low probability of containing a sequencing error. Anchor sequences can be extracted as adjacent, disjoint sequences or as tiled sequences that begin at a fixed step size, to reduce computational burden. For this manuscript, SPLASH was run with default parameters: with 1M reads per FASTQ file, anchor sequences tiled by 5 bp, and *k* = 27. For HLCA datasets, both read 1 and read 2 were used; for other datasets, only read 1. Extracted anchor and target sequences are then counted for each sample with the UNIX command, ′sort | uniq -c′, and anchor-target counts are then collected across all samples for restratification by the anchor sequence. This stratification step allows for user control over parallelization. To reduce the number of hypotheses tested and required to correct for, we discard anchors that have only one unique target, anchors that appear in only 1 sample, and (anchor, sample) pairs that have fewer than 6 counts. Then, we retain only anchors having more than 30 total counts after the above thresholds were applied. This approach efficiently constructs sample by target counts tables for each anchor.

### SPLASH *p*-values

SPLASH’s analysis is based on a new statistical method for analyzing contingency tables that provides *p*-values to reject the null hypothesis that row (target) counts are drawn from the same distribution for all samples. These *p*-values are closed form, meaning that no permutation, resampling or numerical solutions to complex likelihoods are required for significance tests.

We develop a test statistic *S* that has power to detect sample-dependent sequence variation and is designed to have low power when there are a few outlying samples with low counts as follows. First, we randomly construct a function *f*, which maps each target independently to {0,1}. We then compute the mean value of targets with respect to this function. Next, we compute the mean within each sample of this function. Then, an anchor-sample score is constructed for sample *j*, *S_j_*, as a scaled version of the difference between these two. Finally, the test statistic *S* is computed as the weighted sum of these *S_j_*, with weights *c_j_*(which denote class-identity in the two-group case with metadata and are chosen randomly without metadata, see below). In the below equations, *D_j,k_* denotes the sequence of the *k*-th target observed for the *j*-th sample, and *M* denotes the total number of observations of this anchor.

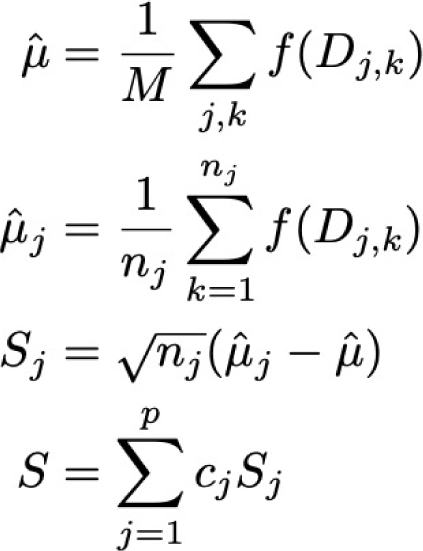

Statistically valid *p*-value bounds (non-asymptotic, providing control of Type I error for finite number of observations) are computed as:

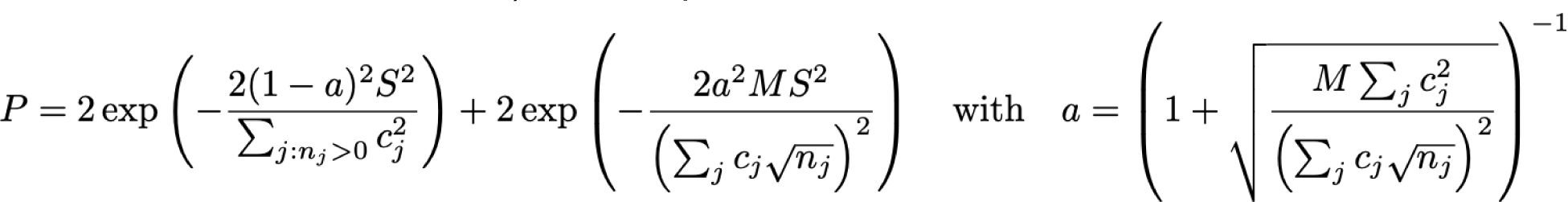

by applying Hoeffding’s inequality on these sums of independent random variables (under the null). The derivation is detailed in the Note S2.

This statistic is computed for *K* different random choices of *f*, and in the case where sample group metadata is not used or available, jointly for each of the *L* random choices of *c*, here with *K* = 10 and *L* = 50. We call the random choice of *c_j_*’s “random *c*’s” below. The choice of *f* and *c* that minimize the *p*-value are reported, and are used for computing the *p*-value of this anchor. To yield valid *p*-values we apply Bonferroni correction over the *L*×*K* multiple hypotheses tested (just *K* when sample metadata is used and randomization on *c* is not performed). Then, to determine the significant anchors, we apply Benjamini-Yekutieli (BY) correction^58^ to the list of *p*-values (for each anchor), yielding valid FDR controlled *q*-values reported throughout the manuscript implemented with the statsmodels.api.stats.multipletests functionality in python.

### SPLASH effect size

SPLASH provides a measure of effect size when the *c_j_*’s used are ±1, to allow for prioritization of anchors with large inter-sample differences in target distributions. Effect size is calculated based on the split *c* and function *f* that yield the most significant SPLASH *p*-value. Fixing these, the effect size is the absolute value of the difference between the mean function value over targets (with respect to *f*) across those samples with *c_j_* = +1 denoted *A_+_*, and the mean over targets (with respect to *f*) across those samples with *c_j_* = -1 denoted *A_-_*.

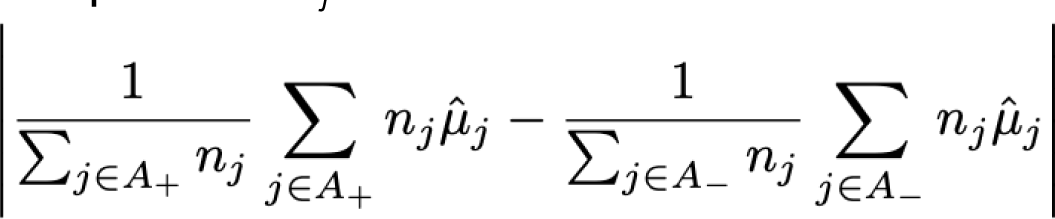

This effect size has natural relations to a simple two group alternative hypothesis. It can also be shown to relate to the total variation distance between the empirical target distributions of the two groups. These connections are discussed further in Note S2.

### Consensus sequences

A consensus sequence is built for each significant anchor for the sequence downstream of the anchor sample. A separate consensus is built for each sample by aggregating all reads from this sample that contain the given anchor. Then, SPLASH constructs the consensus as the plurality vote of all these reads; concretely, the consensus at base pair *i* is the plurality vote of all reads that contain the anchor, *i* base pairs after the anchor appears in the read (a read does not vote for consensus base *i* if it has terminated within *i* base pairs after the anchor appeared). The consensus base as well as the fraction agreement with this base among the reads is recorded. Some empirical behavior of consensuses is shown in Figure S4.

The consensus sequences can be used for both splice site discovery and other applications, such as identifying point mutations and highly diversifying sequences, e.g. V(D)J rearrangements. The statistical properties of consensus building make it an appealing candidate for use in short read sequencing^59^, and may have information theoretic justification in *de novo* assembly^60^ (Note S1).

To provide intuition regarding the error correcting capabilities of the consensus, consider a simple probabilistic model where our reads from a sample all come from the same underlying sequence. In this case, under the substitution only error model, we have that the probability that our consensus for *n* reads makes a mistake at a given location *i* under independent sequencing error rate 𝜖 (substitution only) is at most

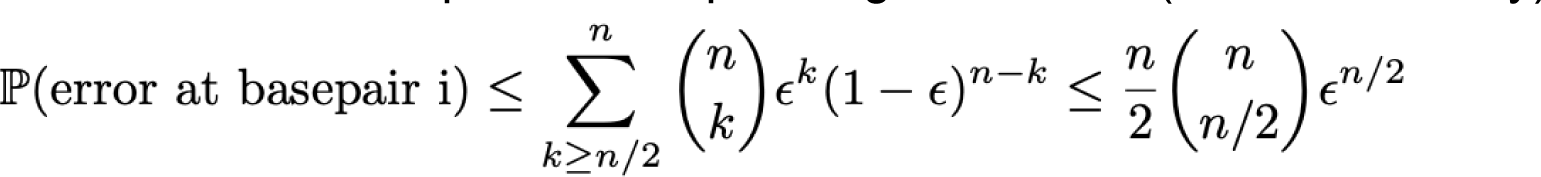

We can see that even for *n* = 10, this probability is less than 1.3E-7 for a given base pair, which we can union-bound over the length of the consensus to yield a vanishingly small probability of error. Thus, for a properly aligned read, if a base pair differs between the consensus and reference it is almost certainly a SNP.

### Element annotations

To identify false positive sequences or contextualize mobile genetic elements, anchors and targets are aligned with bowtie2 to a set of indices, corresponding to databases of sequencing artifacts, transposable elements, and mobile genetic elements^61^. In these alignments, using bowtie2, the best hit is reported, relative to an order of priority. The references used are: UniVec, Illumina adapters, grass carp (*Ctenopharyngodon idella* genome, GCA_019924925)^62^, Escherichia phage phiX174, Rfam^63^, Dfam^64^, TnCentral^65^, ACLAME^66^, ICEberg^67^, CRISPR direct repeats^68^, ITSoneDB^69^, ITS2^70^; and also the reference genome of interest for the study. (Grass carp was used as a control as it contains many artifactual Illumina adapters.) To perform these annotations, bowtie2 indices were built from the respective reference FASTAs, using bowtie2-build with default parameters. Anchors and targets were then aligned to each index, using bowtie2-align with default parameters. For each sequence, we report the alignment to the reference and the position of that alignment for each reference in the prespecified set. Anchors and targets, and their respective element annotations, are reported in the element annotation summary files.

### Genome annotations

Anchor, target, and consensus sequences can be aligned by SPLASH to reference genomes and transcriptomes, to provide information about the location of sequences relative to genomic elements. All alignments reported are run in two modes in parallel: bowtie2 end-to-end mode (the bowtie2 default parameters) and bowtie2 local mode (′-local′, in addition to the bowtie2 default parameters). To report alignments to the transcriptome, the sequences are aligned to the reference transcriptome with bowtie2, with ′-k 1′, in addition to the above parameters, to report a maximum of one alignment per sequence. If there is a transcriptome alignment, we report the alignment to the reference and the MAPQ score of the alignment. To report alignments to the genome, the sequences are aligned to the reference genome, with the same parameters above. If there is a genome alignment, we report the alignment to the reference, the strand of the alignment, and the alignment MAPQ score.

### Splice junction calls

To identify exon coordinates for reporting annotations in this manuscript, consensus sequences are mapped with STAR aligner (default settings)^71^. Gapped alignments are extracted and their coordinates are annotated with known splice junction coordinates using ‘bedtools bamtobed --split’; each resulting contiguously mapping segment is called a “called exon”. From each consensus sequence, called exons are generated as start and end sites of each contiguously mapped sequence in the spliced alignment. These ‘called exons’ are then stratified as start sites and end sites. Note that the extremal positions of all called exons would not be expected to coincide with a splice boundary; “called exon” boundaries would coincide with an exon boundary if they are completely internal to the set of called exon coordinates. Each start and end site of each called exon is intersected with an annotation file of known exon coordinates; it receives a value of 0 if the site is annotated, and 1 if it is annotated as alternative. The original consensus sequence and the reported alignment of the consensus sequence are also reported. Gene names for each consensus are assigned by bedtools intersect with gene annotations (hg38 RefSeq for human data by default), possibly resulting in multiple gene names per consensus.

### SPLASH protein domain profiles

Custom scripts were used to generate protein domain profiles. For each set of enriched anchors, homology-based annotation was attempted against an annotated protein database, Pfam^14^. For each dataset, up to 1000 of the most significant anchors (*q*-value < 0.01) were retained for the following analysis: we first generated a substring of each downstream consensus by appending each consensus nucleotide assuming both conditions were met: a minimum observation count of 10 and a minimum agreement fraction of 0.8, until whichever metric first exhibited two consecutive failures at which point no further nucleotide was added. A limit of 1000 anchors was used due to computational constraints from HMMer3 (see below). Anchors that did not have any consensus nucleotides appended were kept as is. An extended anchor was generated for each experiment in which an anchor was found. Each extended anchor was then stored in a final concatenated multi FASTA file with unique seqID headers for each experiment’s extended anchors.

To assess these extended anchors for protein homology, this concatenated FASTA file was then translated in all six frames with the standard translation table using seqkit^72^ prior to using hmmsearch from the HMMer3 package^73^ to assess resulting amino acid sequences against the Pfam35 profile Hidden Markov Model (pHMM) database. The resulting ‘raw’.tblout outputs were then processed, keeping the best hit (based on E-value) per each initial anchor, and any hits with an E-value better than 0.01 were parsed into an *_nomad.Pfam (or *_control.Pfam) file used for subsequent plotting.

All hits to the Pfam database were then binned at different E-value orders of magnitude and plotted. In each case, control assessments were performed by repeating the extension and homology searches against an equivalent number of control anchors (see below). The number of matched anchors used for SPLASH and control analysis per dataset were as follows: 201 high effect size (.5) anchors in SARS-CoV-2 from South Africa, 252 high effect size (.5) anchors in SARS-CoV-2 from France; 1000 anchors (no effect size filter) were used for rotavirus, human T cells, human B cells, *Microcebus* natural killer T cells, and *Microcebus* B cells. We note that while the number of input anchors for SPLASH and control sets are matched, it is possible to have more control protein domains in the resulting barplots, as only high E-value hits to Pfam are reported in the visualizations. Domain profiling summaries are in Data S1.

A hypergeometric test was used to give *p*-values for protein domain analysis. For a given domain, we construct the 2×2 contingency table, where the first row is the number of SPLASH hits for this domain, followed by the total number of SPLASH hits not in this domain. The second row is the mirror of this for control, where the first entry is the number of control hits for this domain, followed by the total number of control hits not in this domain. A one-sided *p*-value is computed using Fisher’s exact test, which is identically a hypergeometric test. We apply Bonferroni correction for the total number of protein domains expressed by either SPLASH or control, to yield the stated *p*-values.

Lastly it is worth noting that while only counts of the best scoring Pfam hits were assessed in this study, other information is also produced by HMMer3. In particular, relative alignment positions are given for each hit which could be used to more finely pinpoint the precise locus at which sequence variation is detected.

### Control analyses

To construct control anchor lists based on abundance, we considered all anchors input to SPLASH and counted their abundance, collapsing counts across targets. That is, an anchor receives a count determined by the number of times it appears at an offset of 5 in the read up to position R - max(0,R/2-2*k) where R is the length of the read, summed over all targets. The 1000 most abundant anchors were output as the control set. For analysis comparing control to SPLASH anchors, min (|SPLASH anchor list|, 1000) most abundant anchors from the control set were used and the same number of SPLASH anchors were used, sorted by *p*-value.

### Generation of contingency table heatmaps

To plot the anchor-target heatmaps, we exclude targets with low counts. Concretely, we by default filter out targets that occur fewer than 5 times, have less than 5% of the total counts of that anchor, and retain at most the top 10 targets, while ensuring that at least 2 targets are plotted. Then, all samples with fewer than 5 counts are discarded. For clarity of presentation, we include or remove rows corresponding to additional targets based on biological relevance.

### SARS-CoV-2 analysis

SARS-CoV-2 data was downloaded from the NCBI: France^9^ (SRP365166) and South Africa^8^ (SRP348159). Sample metadata for the France dataset was provided by the authors via personal communication, and consists of their calls of the primary and secondary infecting strains for each patient sample. We note that sample ‘WTA-022002271301_S1’ appeared to be mislabelled, appearing in the metadata file but not in the NCBI sample list. Conversely, the sample ‘Pl924-022002271301_S1472’ appears in the NCBI sample list, but not in the metadata file. Thus, we associate these labels to each other, to obtain metadata labeling for all 106 samples. We do not have information regarding which samples are replicates. We provide the NCBI sample list and the strain metadata file as Data S8.

The SARS-CoV-2 datasets used in this manuscript were analyzed with SPLASH’s unsupervised mode (no sample metadata provided). To identify high effect size anchors, a threshold of ′effect_size_randCjs′ > 0.5 was used (table S2).

For the purposes of strain-defining mutation analysis, we manually constructed “archetype” genome sequences for variant strains Delta, Omicron BA.1, and Omicron BA.2 by editing the Original (Wuhan) reference NC_045512.2 to contain all (and only) the defining mutations specified at CoVariants.org^10^; these are provided as Data S5.

To determine what SPLASH calls (and control anchors) were strain defining we perform the following. To generate SPLASH’s calls, we filter for anchors that are significant (with a BY corrected *p*-value less than .05) and have large effect size (> .5), yielding a list of *N* SPLASH-called anchors. Control anchors are generated by taking the *N* anchors with the highest counts. For each of these anchors we construct their target × sample contingency table, first filtering out all anchors with fewer than 30 counts, only 1 unique target, or only 1 unique sample, and filtering out all samples with 5 or fewer counts. Then, we discard all targets that constitute less than 5% of the remaining counts for that anchor. The remaining anchors and targets are then bowtie aligned to an index comprised of the Wuhan reference genome (NC_045512.2), and Delta, Omicron BA.1, and Omicron BA.2 archetype genomes. For this alignment, options ′-a -v 0′ were used. Then, for each set of anchors (SPLASH calls, and controls), the list is filtered to only anchors that align perfectly to at least one of the reference assemblies, further requiring that each anchor have at least one target that aligns perfectly to a reference assembly. Then for each anchor, we declare it to be strain defining if, for any of the reference assemblies, it has at least one target that maps to it and one target that does not.

### Identifying cell-type specific isoforms in single-cell data (lung macrophages and capillary cells)

The human lung scRNA-seq data used here (HLCA SS2)^20^ is accessible through the European Genome-phenome Archive (accession number: EGAS00001004344); FASTQ files from donor 1 (P2) and donor 2 (P3) generated with the Smart-seq2 protocol were used. In the analysis of HLCA SS2 data, we utilize “isoform detection conditions” for alternative isoform detection. These conditions select for (anchor, target) pairs that map exclusively to the human genome, anchors with at least one split-mapping consensus sequence, *mu_lev* > 5, and *M* > 100. *mu_lev* is defined as the average target distance from the most abundant target as measured by Levenstein distance. *M* is defined as the total number of counts in the anchor’s contingency table. To identify anchors and targets that map exclusively to the human genome, we included anchors and targets that had exactly one element annotation, where that one element annotation must be grch38_1kgmaj. To identify anchors with at least one split-mapping consensus, we selected anchors that had at least one consensus sequence with at least 2 called exons. The conditions on Levenshtein distance, designed to require significant across-target sequence variation, significantly reduced anchors analyzed (excluding many SNP-like effects). We further restricted to anchors with *M* > 100, to account for the lower numbers in macrophage cells; note that the user can choose to use a lower *M* requirement, based on input data. These isoform detection parameters were used to identify the SS2 examples discussed in this manuscript, MYL12. For HLA discussion, gene names were called using consensus_gene_mode.

While here we focus on anchors that have aligned to the human genome, we note in passing that SPLASH makes many predictions of cell-type specific RNA expression that include sequences that map to repetitive elements or do not map to the human reference: for individual P2 (respectively P3), 53% (61%), of 4010 (4603) anchors map to the human genome and no other reference; 35% (30%) map to both the Rfam and human genome; 6% (7%) have no map to any reference used for annotation which includes repetitive and mobile elements. As an example, 9 and 18 such anchors (donor 1 and 2 respectively) BLAST to MHC alleles in the NCBI database.

### Immune single-cell analysis

To study human B and T cells, we utilize Tabula Sapiens data (Smart-seq2)^34^, downloaded from https://tabula-sapiens-portal.ds.czbiohub.org/; B cells were used from donor 1 (TSP1) and CD4+ T cells from donor 2 (TSP2). Mouse lemur single-cell RNA-seq data used in this study was generated as part of the Tabula Microcebus consortium^36^; the FASTQ files were downloaded from https://tabula-microcebus.ds.czbiohub.org. B cells and natural killer T cells were analyzed separately; both were from spleen and were a mixture of individual L2 and L4. To determine the most frequent transcriptome annotation for a dataset, all significant anchors were mapped to the human transcriptome (GRCh38, Gencode) with bowtie2, using default parameters and ′-k 1′ to report at most one alignment per anchor (Data S4). Then, the bowtie2 transcript hits are aggregated by counting over anchors. The transcript hits with the highest counts over all anchors were reported. Protein domain profiling was performed as described above.

### SPLASH comparison to BASIC analysis in lemur spleen B cells

As part of the Tabula Microcebus consortium^36^, mouse lemur B cells were annotated with BASIC^38^ to identify Ig variable domains. However, BASIC was unable to identify the light chain variable domain in 35 cells. We used a simple approach to see if SPLASH could identify variable domains in these uncalled B cells. We checked the SPLASH genome annotations for these cells for anchors mapping to human “IGL” or “IGK” genes; there were only five such anchors, all to IGL, and these were found in only eight cells. For those eight cells, we retrieved the SPLASH consensus sequences for these anchors, which ranged from 2-5 per cell. Where consensuses for a cell overlapped, one was chosen, and these were submitted to BLAST against the nr/nt database. Many hits were to “immunoglobulin lambda-like polypeptide” 1 or 5 (IGLL), surrogate light chain genes that contain sequence similar to lambda J and C regions (as well as a unique N-terminal region) and so could mimic alignment to a true lambda variable domain. Therefore, BLAST alignments were checked to see whether the match could be assigned to V, J, C, or IGLL-unique regions. 4 cells matched C-region, 2 matched IGLL-unique region, and 2 had sequence beyond J-region (presumed V-region). For the latter two, we attempted to extend the consensus further into the V-region by ′grep′ in raw reads; one could not be extended as it only had adaptors adjacent to J-region sequence. The other consensus was extended through the full V-region, and its sequence is given in Note S4, along with the IGLL-unique matches.

### SPLASH for Zostera marina (eelgrass) and Octopus bimaculoides

Data was downloaded from SRP327909 (eelgrass^42^), and SRP278619 (Octopus^43^), using nf-core fetchngs run in default mode ^74^ and preprocessed with FASTP ^75^ run in default mode to mitigate false positive calls due to adapter concatenation. An updated version of SPLASH ^54^ was run with gap length=0, anchor_unique_targets_threshold=1, anchor_count_threshold=50, anchor_samples_threshold=1, anchor_sample_counts_threshold=5, and excluding anchor-targets containing poly A / C / G / T run of length 8. 500 pairs of random *c* and *f* were chosen. Fastp v0.3.9 was installed on 2/23/23 using bioconda.

Anchor-targets were selected from the SPLASH calls if they had homopolymer length ≤ 5, effect size > 0.1, and corrected *p*-value < 0.01. Element annotations were run using SPLASH commit ID 728066b; anchor-targets mapping to UniVec, Illumina adapters, SARS-CoV-2, or grass carp were removed. Anchor-targets were aligned with STAR 2.7.5 to a reference index generated from either the *O. bimaculoides* reference genome^76^ (NC_068981.1) and transcriptome^43^; or the *Z. marina* nuclear genome^77^ (v3.1, https://phytozome-next.jgi.doe.gov/info/Zmarina_v3_1) and mitochondrial and chloroplast genomes^78^ (NC_035345.1 and NC_036014.1, respectively).

For protein domain profiling, anchor-targets with no element annotation were *in silico* translated in all six frames and submitted to HMMer search of the Pfam database. For Figure 5B (eelgrass), anchors were ordered by descending number of observations; the top 200,000 were concatenated with their anchor-targets and submitted to Pfam; these anchor-targets are defined as controls. The best Pfam hit by full sequence E-value is retained for each anchor-target. For each domain, the top 4 domains hit by anchor-targets and by controls are plotted, where bar height is the number of anchor-targets that hit the domain with a full-sequence E-value < 0.01.

For BLAST analysis, we selected anchors with no more than 1 target mapping with either STAR to the reference genome or Bowtie2 element annotations, thus cases where no sample-specific variation would be detected if a reference genome were used. In eelgrass, targets were selected if their fraction exceeded 0.5 and their anchor’s effect size exceeded 0.9; in octopus, targets were selected if their fraction exceeded 0.3 and if their anchor’s effect size exceeded 0.8. The 1808 anchor-target pairs satisfying these criteria in octopus were submitted to BLAST with parameters:

-db nt -evalue 0.1 -task blastn -dust no -word_size 24 -reward 1 -penalty -3

-max_target_seqs 4

BLAST hits were merged into SPLASH output, with an indicator variable for whether the sequence was queried. For octopus anchor-targets, 1061/1808 had a BLAST hit (max E-value 0.028). There were 288 hits to octopus, of which 281 were annotated as from *O. sinensis* and 7 from *O. bimaculoides*. Selected sequences mapping to *O. sinesis* were further analyzed. For eelgrass, 1606/4081 had a BLAST hit (max E-value 0.028).

SPLASH output, merged with Pfam and BLAST analyses, are in Data S6 and Data S7 (octopus and eelgrass, respectively).

## Supplemental Information

**Document S1.** Figures S1 to S7; Notes S1 to S7.

**Data S1.** Protein domain profiling output tables.

**Data S2.** Tables for SARS-CoV-2 and single-cell analyses: significant anchors, anchor statistics, and *c_j_*’s used for each anchor.

**Data S3.** Additional summary tables for macrophage and capillary single-cell analyses: significant anchors, their targets, anchor statistics, anchor and target reverse complement information, highest priority element annotations for anchors and targets, anchors annotations, and consensus annotations.

**Data S4.** Tables for human and lemur immune single-cell analyses: significant anchors, and their genome and transcriptome annotations.

**Data S5.** Artificial genome sequences with defining mutations for SARS-CoV-2 Delta, Omicron BA.1, and Omicron BA.2 strains.

**Data S6.** Table for *Octopus bimaculoides* analysis: significant anchors, their targets, anchor statistics, STAR mapping annotations, Pfam results, and BLAST results.

**Data S7.** Table for *Octopus bimaculoides* analysis: significant anchors, their targets, anchor statistics, STAR mapping annotations, Pfam results, and BLAST results.

**Data S8.** SARS-CoV-2 metadata.

## Document S1

**Figures S1 to S7**

**Figure S1.** SPLASH computations.

**Figure S2.** Rotavirus protein domain profiling.

**Figure S3.** Effectiveness of SPLASH randomized sample splitting.

**Figure S4.** Additional details for single cell analyses.

**Figure S5.** Lemur COX2 allelic detection, additional Ig/TCR anchors.

**Figure S6.** O. bimaculoides 3’ UTR anchors show tissue-specific expression.

**Figure S7.** Diatom anchors in eelgrass samples show variation with location/season or Day vs Night.

### Supplemental Figure Legends

**Notes S1 to S7**

**Note S1.** Generality of SPLASH.

**Note S2.** SPLASH statistical inference.

**Note S3.** SPLASH is robust to parameter choices and effective without metadata.

**Note S4.** Lemur lamba light chain and surrogate light chains found by SPLASH.

**Note S5.** Octopus and eelgrass analyses, additional notes.

**Note S6.** SPLASH runs on a laptop.

**Note S7.** Anchor and target sequences, q-values, and binomial p-values.

## Supplemental Figure Legends

**Figure S1.**
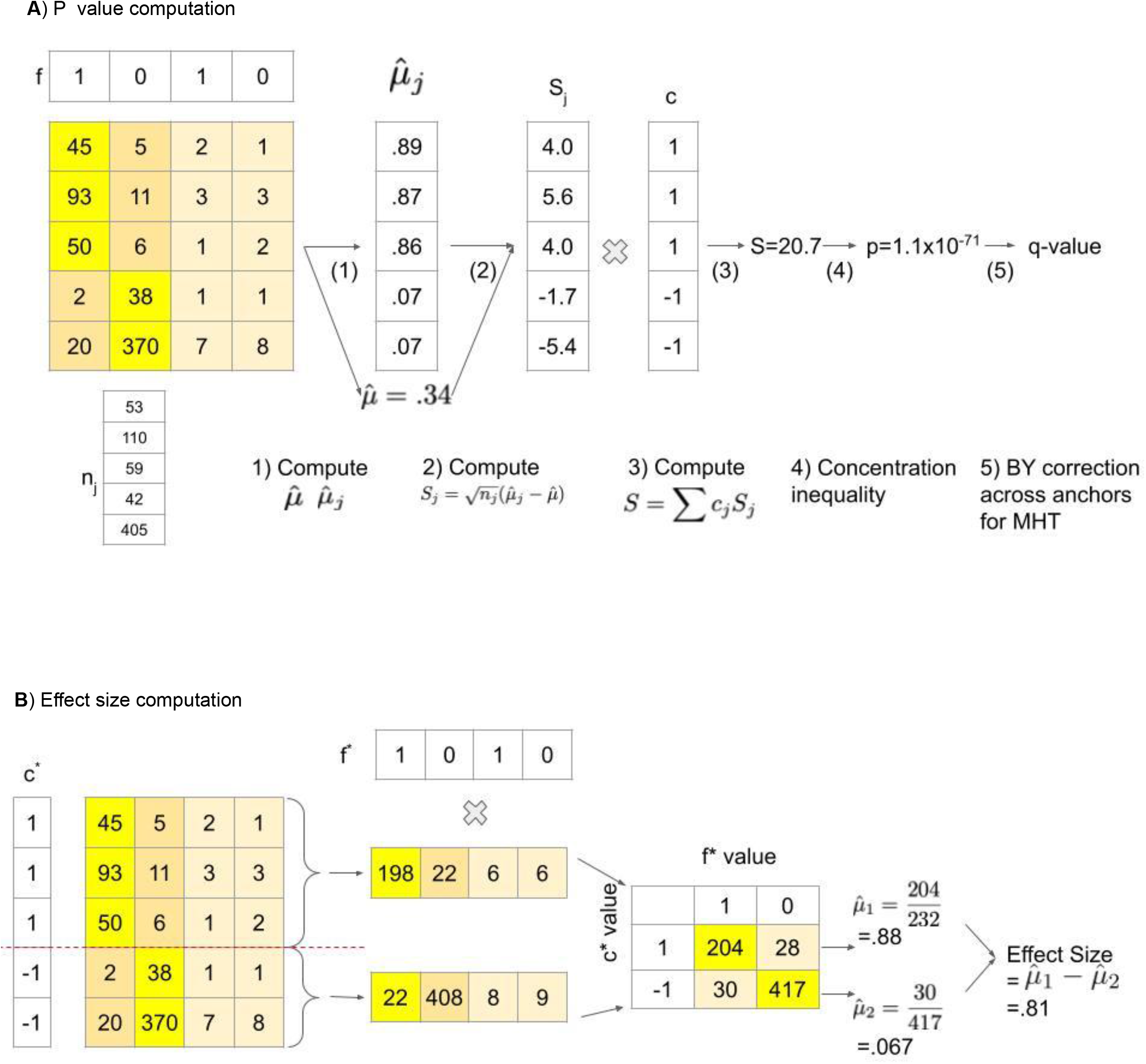
SPLASH computations. A. *p*-value computation for SPLASH for user specified vectors *f* and *c*. Contingency table transposed for visual convenience (rows are samples and columns are targets). Starting with a samples by targets counts matrix, SPLASH utilizes one (or several) functions *f* mapping targets to values within [0,1]. The mean with respect to *f* is taken over the targets in each row *j* to yield µ̂_*j*_, and an estimate for 𝑗 the mean over all target observations of *f* is taken, yielding µ̂. The anchor-sample scores *S_j_* are then constructed as the difference between the row mean µ̂_*j*_ and the overall mean µ, and is scaled by 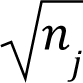. These anchor-sample scores are weighted by *c_j_* in [-1,1] and summed to yield the anchor statistic *S*. Finally, a value is computed utilizing classical concentration inequalities, which we correct for multiple hypothesis testing (with dependence) by constructing values using Benjamini-Yekutieli, a variant of Benjamini Hochberg testing which corrects for arbitrary dependence. B. Effect size computation for SPLASH. Reported effect size is calculated based on the random split *c* and random function *f* that yielded the most significant SPLASH *p*-value. Fixing these, the effect size is computed as the difference between the mean across targets (with respect to *f*) across those samples with *c_j_* = +1, and the mean across targets (with respect to *f*) across those samples with *c_j_* = -1. This should be thought of as studying an alternative where samples from *c_j_* = +1 have targets that are independent and identically distributed with mean (under *f*) of µ_1_, and samples with *c_j_* = -1 have targets that are independent 1 and identically distributed with mean (under *f*) of µ_2_. The total effect size is 2 estimated as µ_1_ − µ_2_.

**Figure S2.**
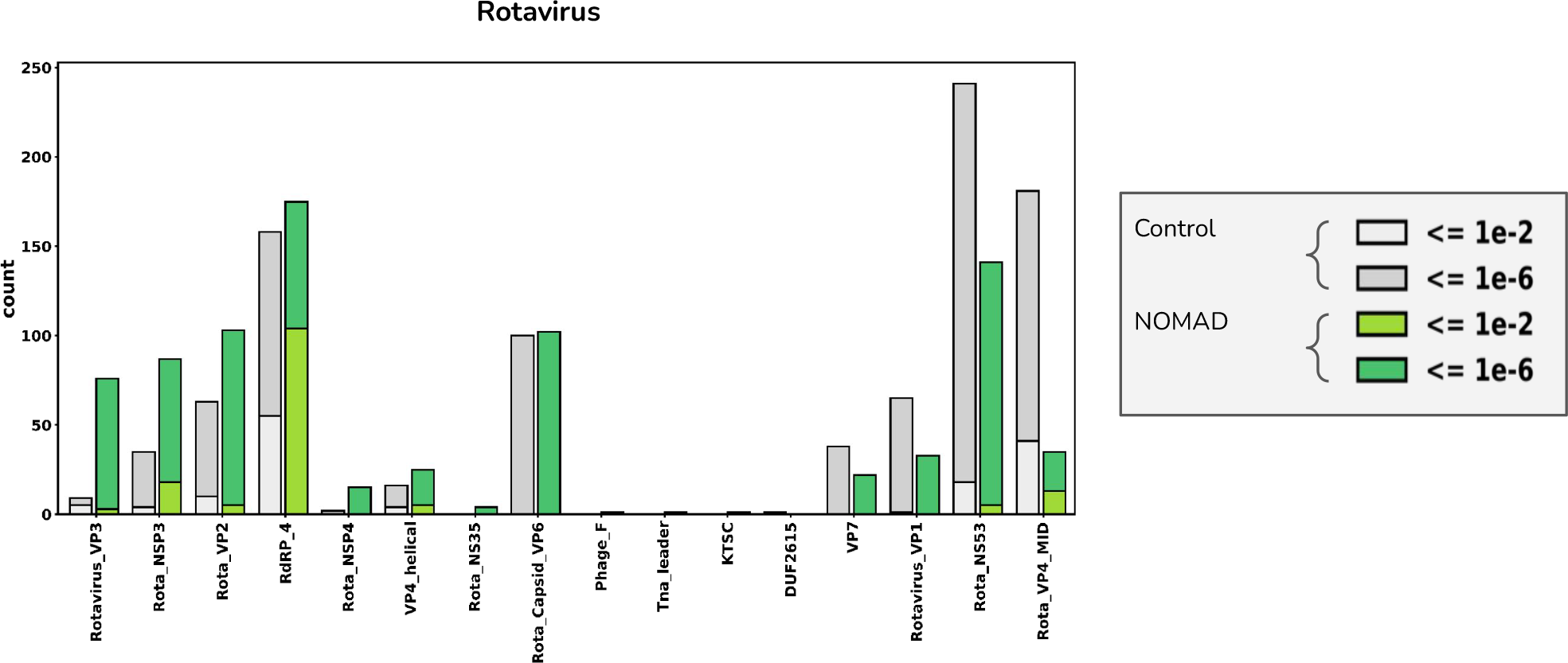
Rotavirus protein domain profiling. We performed SPLASH protein domain profiling in a dataset of virally enriched samples from breakthrough infections in patients that had been vaccinated against rotavirus; nearly all did have rotavirus infection, but some also had other coinfections^17^. The top domains were rotavirus VP3 (Rotavirus_VP3, 76 SPLASH hits vs 9 control hits) followed by rotavirus NSP3 (Rota_NSP3, 87 SPLASH vs 35 control hits), indicating that these rotavirus genes have sequences that vary significantly among these patients; they have roles in blocking host innate immunity^18^, and so may be under selective pressure for variation.

**Figure S3.**
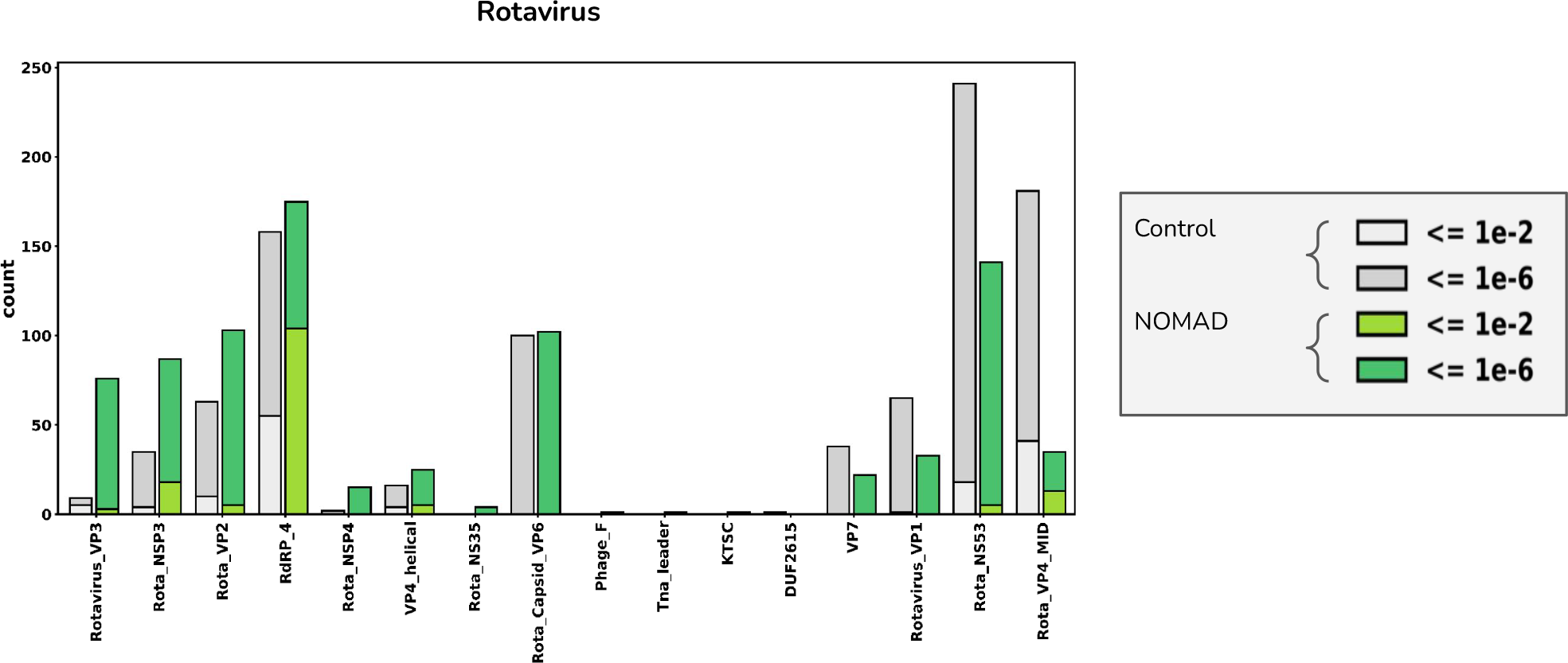
Effectiveness of SPLASH randomized sample splitting. Random *c*’s can recover samplesheet *c*’s. For the HLCA dataset, of the 3439 anchors (1384 genes) called by the input metadata (samplesheet *c*’s) in donor 1 (BY correction, alpha=.05), we have that 72% of the genes called were also called by SPLASH’s selection of random c’s (6287 called by anchors by random *c*’s, 2268 genes). Left plot indicates for each gene (dot) how many times it was called by samplesheet *c*’s vs random *c*’s. Red dots indicate those genes not called by random c’s. On the right plot we have the fraction of genes that are called at least *x* times by samplesheet *c*’s that are also called by random *c*’s. We see that for *x* = 2 (i.e. all genes hit by at least 2 anchors), random *c*’s call >94% of those genes called by samplesheet *c*’s. For donor 2 similar results are observed, with 3775 (5619) anchors from samplesheet *c*’s and 1125 (1844) genes for samplesheet *c*’s (random *c*’s) respectively. >90% of samplesheet c discoveries for *x* = 2, >94% for *x* = 3.

**Figure S4.**
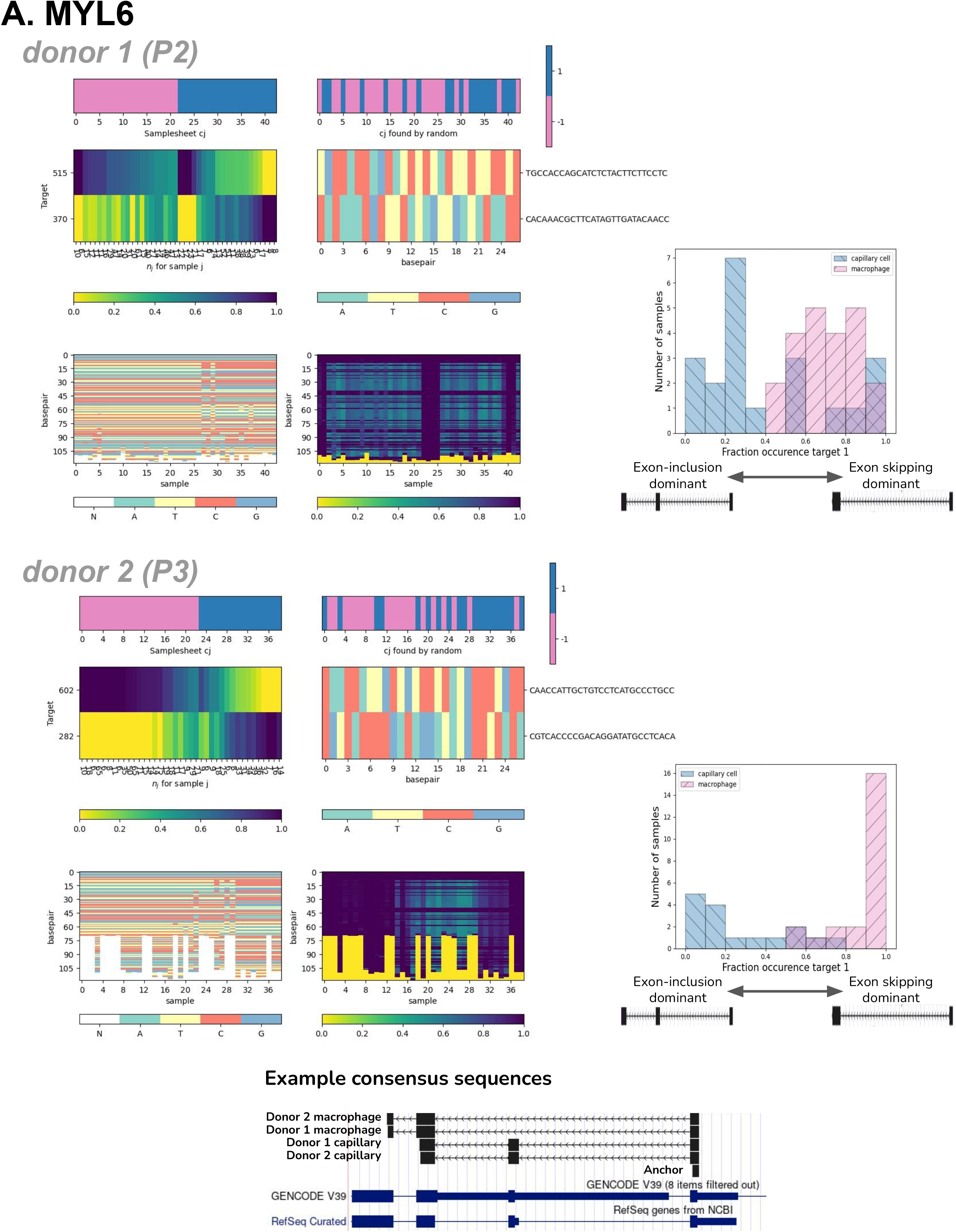
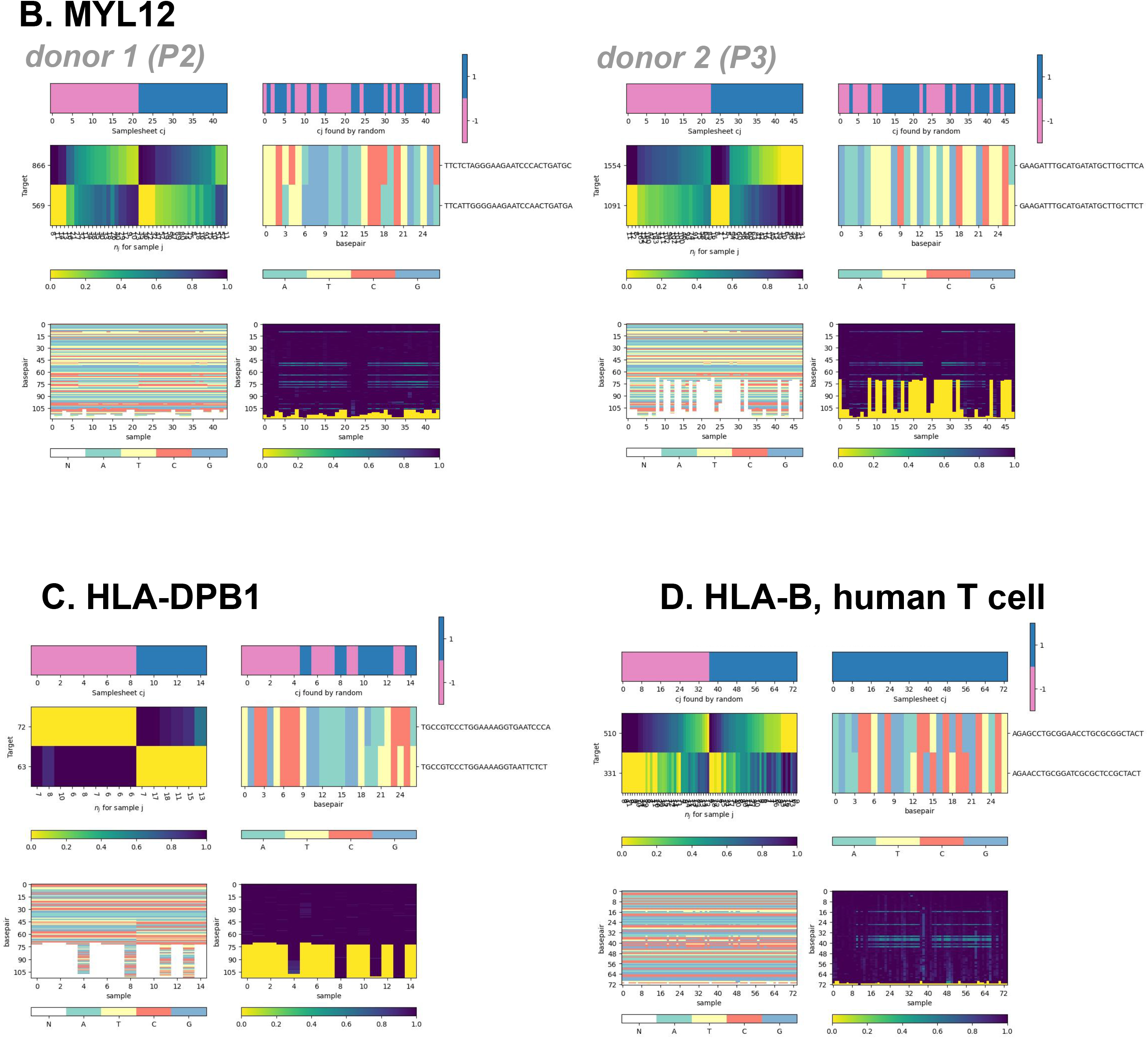
Additional details for single cell analyses. Heatmaps show the complete data for the called anchors. Each set of heatmaps is for one anchor sequence. The primary plot is the center left one, which shows the samples × targets contingency table. Each column represents a sample, and each row represents a unique target. The color indicates what fraction of the sample’s (column’s) targets come from the target corresponding to that row. The *x*-ticks correspond to *n_j_*, the number of times the anchor was observed in this sample. The *y*-ticks indicate the number of times this target appeared (following this anchor), and the targets are sorted by abundance. The two top plots indicate the *c_j_*’s used; when samplesheet *c_j_*s are available, they will be in the upper left, and the optimizing random *c_j_*s will be in the upper right. The middle left plot is used to visualize the targets that follow this anchor. Each row represents a target (sequence given in *y*-tick) corresponding to the row to the left of it in the contingency table. The columns are base pair positions along the sequence of each target. Each nucleotide is color-coded, to show the similarity of the targets (e.g. to indicate whether they differ by a SNP, deletion, alternative splicing, etc). The two bottom plots relate to the consensus sequences. The lower left plot shows the nucleotide sequence (same color scheme as the center right one for the targets). Each column corresponds to the consensus sequence for the sample of the same column above it in the contingency table. The rows are base pair positions along each consensus. These consensus sequences are variable length, and a value of -1 (yellow color) on the bottom of a sequence indicates that the consensus has ended. The bottom right plot shows the fraction agreement per nucleotide within a sample with its consensus sequence. We can see that for samples where only one isoform / SNP is expressed the consensus stays near 100%, while for samples with a diverse set of targets the consensus is less uniform. Panels A-C are macrophage and capillary cells from human lung (HLCA dataset). In the samplesheet *c_j_*s metadata heatmap (upper left) and histogram plots for panel A, pink is macrophage and blue is capillary cell. **A. MYL6.** SPLASH rediscovers alternative splicing in MYL6: capillary cells express more of the exon-inclusion variant, while macrophages express predominantly the exon-skipped variant. At the bottom is a UCSC Genome Browser screenshot, showing BLAT mapping of macrophage and capillary consensus sequences which fully capture the inclusion or exclusion of the short alternative exon. **B. MYL12**. This data is partly presented in Figure 3A. **C. HLA-DPB1**. This data is partly presented in Figure 3C. **D. HLA-B (human T cells, Tabula Sapiens)**. This data is partly presented in Figure 3D. In this case, all cells have the same metadata label (CD4+ T cell) so only the optimizing random *c_j_*s heatmap is relevant and so it is shown in the upper left. The best random *c_j_*s found divide the cells into two groups that differ in their relative distribution of targets (note that other *c_j_*s might give an even larger difference), indicating that HLA-B expression is not uniform across this set of cells.

**Figure S5.**
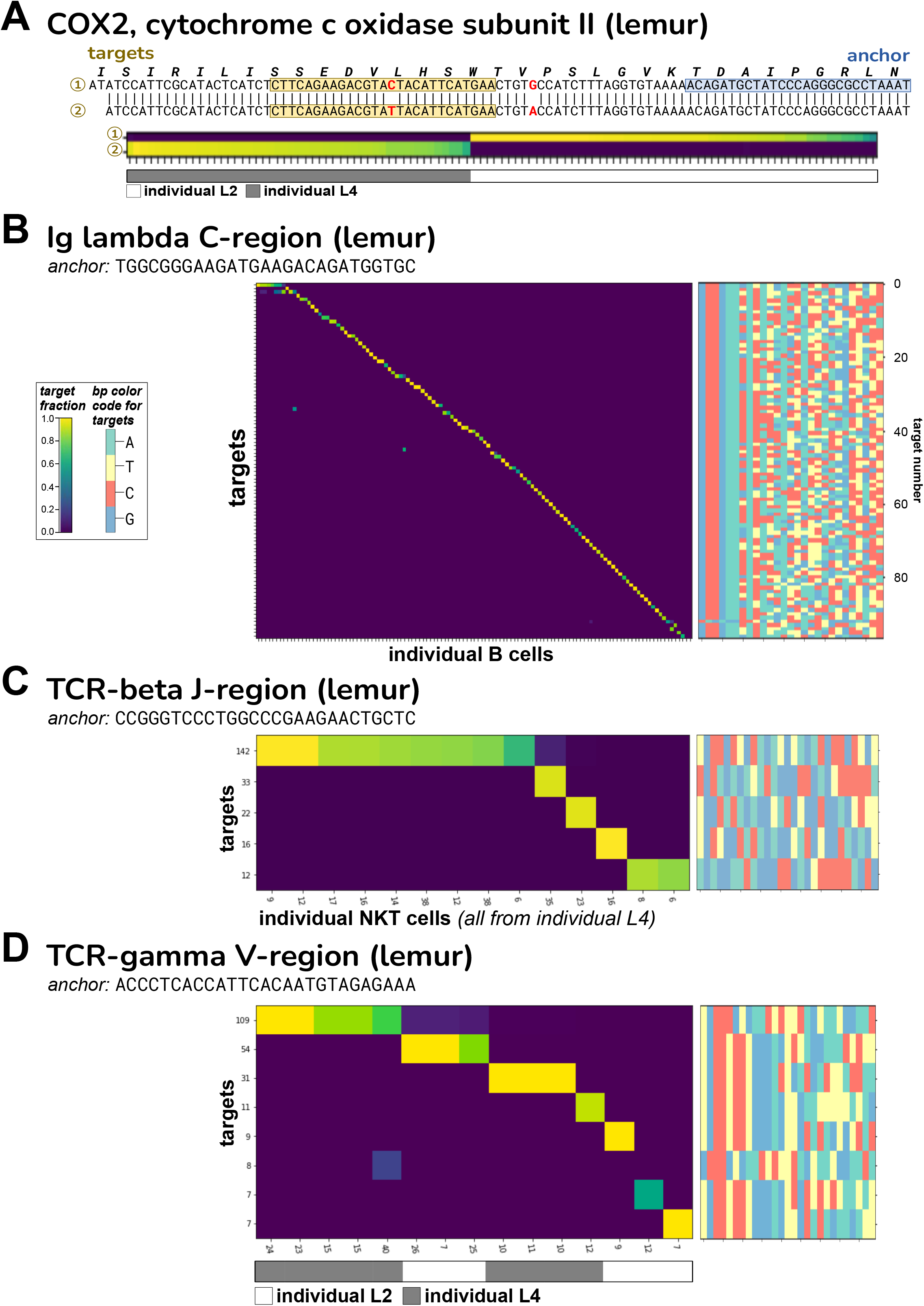
Lemur COX2 allelic detection, additional Ig/TCR anchors. For panels B-D, target × sample heatmaps are shown on the left (samples are individual cells), and bp color-maps on the right. The latter encode the target sequences with different colors for each type of base, and are only provided to give a quick visual impression of the sequence variability. The small numbers at the sides of the heatmaps are summed target counts (over rows or columns). **A. COX2**. SPLASH detects numerous anchors that have targets that correspond precisely to the identities of the two lemur individuals represented in the set of cells analyzed. One example is an anchor in COX2. The alignment of two consensuses (both from NKT cells) shows that targets 1 and 2 differ at a single silent position (C vs T) and another silent SNP is also present outside the target region (G vs A); these positions are highlighted in red. Consensuses 1 and 2 align perfectly to different lemur mitochondrial genome accessions (NC_028718.1 and KR911907.1). Targets 1 and 2 are found exclusively in individuals L4 and L2, respectively. **B. Ig-lambda C-region**. Depicted is a lemur anchor that maps to the 5’ end of the human Ig-lambda C3 segment. It has 97 different targets, which lie within the hypervariable CDR3 region. Nearly all targets are expressed clonotypically (i.e., cells do not share targets), except for targets 1 and 3 (at the top of the heatmap). **C. Ig-beta J-region**. We analyzed natural killer T (NKT) cells from lemur. In humans and mice, a subset of NKT cells express stereotypical or shared TCR genes; Figure 4B shows an example anchor in TCR-alpha. Here we show an anchor that maps to a human TCR-beta J-region (J2-1); targets reside in the V-region. A number of the NKT cells express a shared TCR-beta target (seen in the top row of the heatmap). All of these cells derive from individual L4. **D. Ig-gamma V-region**. There is evidence for shared TCR-gamma sequences in NKT cells as well. This anchor maps to a human TCR-gamma V-region (TRGV9); the targets lie at the V-J junction. Individual L4 has two different shared targets (rows 1 and 3) while individual L2 shows three cells expressing a shared target (row 2). Other NKT cells express unique targets, however.

**Figure S6.**
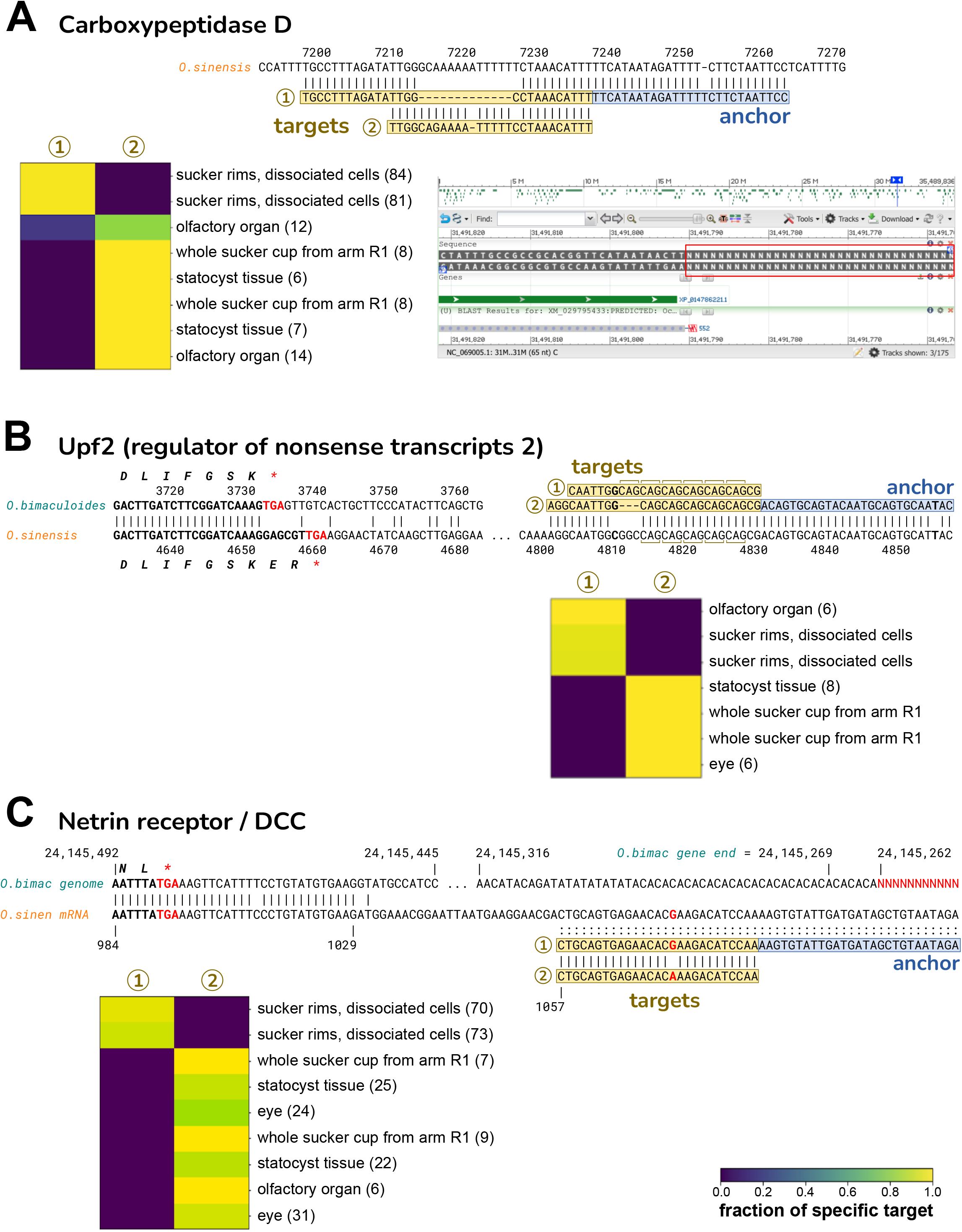
*O. bimaculoides* 3’ UTR anchors show tissue-specific expression. In the heatmap, the parenthetical numbers are summed anchor counts. **A. Carboxypeptidase D (CPD).** The anchor and targets align to the 3’ UTR of the *O. sinensis* CPD mRNA (XM_029795433.2), but are not found in the *O. bimaculoides* genome assembly. The NCBI Browser screenshot at lower-right shows that the 3’ UTR of the *O. bimaculoides* CPD gene (LOC106880679, Ch.25) is entirely missing from the genome: right after the end of the coding region, a run of Ns begins (red box). (There is a second *O. bimaculoides* CPD gene on Ch.16, LOC106873734, but has much lower identity to the sole *O. sinensis* CPD.) Target 2 is identical to *O. sinensis* except for two mismatches; target 1 has a 12-nt deletion relative to target 2. Target 1 is only expressed in dissociated cells from sucker rims, and at a low level in one olfactory organ sample. All other tissues express only target 2. **B. Upf2 (regulator of nonsense transcripts 2)**. The alignment of Upf2 mRNAs from *O. bimaculoides* (XM_014915650.2) and *O. sinensis* (XM_036513028.1) shows that they diverge just before the stop codon, so the 3’ UTRs are unrelated. Our anchor-targets from *O. bimaculoides* map only to *O. sinensis* Upf2; neither the anchor-targets nor the *O. sinensis* 3’ UTR match anywhere in the *O. bimaculoides* genome. The alignment also shows the downstream portion of the *O. sinensis* 3’ UTR where the anchor-targets map. The targets differ in the number of tandem repeats of CAG: target 1 and 2 have six and five repeats, respectively. Target 1 is expressed in dissociated cells from sucker rims, and in olfactory organ; the other tissues express target 2. **C. Netrin receptor/DCC**. The alignment of the *O. bimaculoides* genome (gene LOC106883766) and *O. sinensis* mRNA (XM_036508072.1) shows that the two diverge shortly after the stop codon. The *O. bimaculoides* gene ends in dinucleotide repeats just before the genome becomes a run of Ns (in red). Our anchor-targets from *O. bimaculoides* map only to *O. sinensis* netrin receptor 3’ UTR (also shown in the alignment); neither the anchor-targets nor the *O. sinensis* 3’ UTR match anywhere in the *O. bimaculoides* genome. The targets differ at a single nucleotide: target 1 and 2 have G and A, respectively; *O. sinensis* has a G in this position. If the *O. bimaculoides* genome encodes A, then target 1 is consistent with A-to-I RNA editing (inosine read as G during reverse transcription). The majority of tissues express target 2 only, while target 1 is only expressed in dissociated cells of sucker rims.

**Figure S7.**
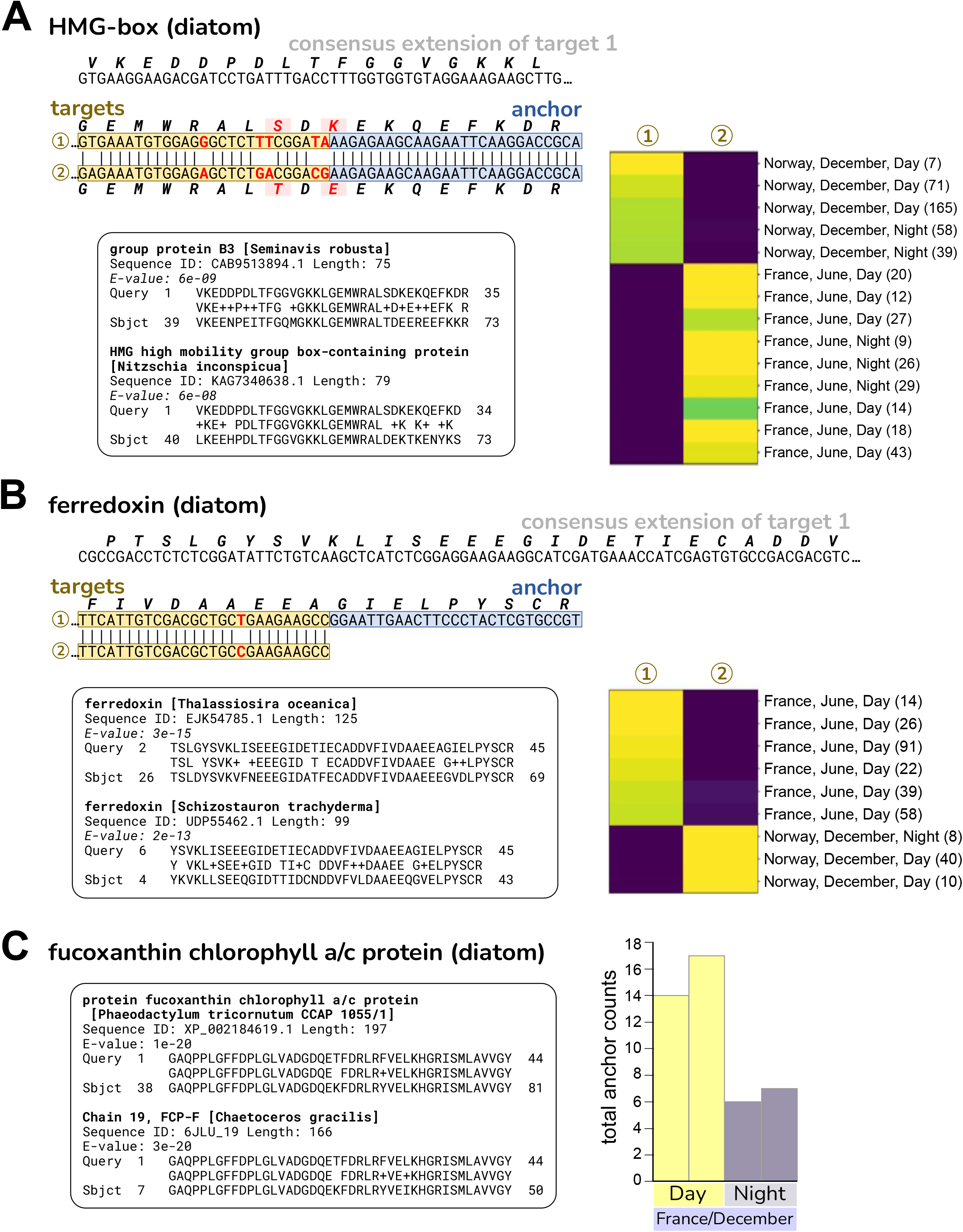
Diatom anchors in eelgrass samples show variation with location/season or Day vs Night. **A**. **HMG (high mobility group) box domain**. The two targets show several nucleotide differences that result in two coding differences. The translation of the consensus sequence has its best two protein BLAST matches to HMG box proteins from diatom species, shown in the inset. Its best Pfam match is also HMG_box. Target 1 is found only in Norway/December samples, while target 2 is found only in France/June samples; both targets are found in both Day and Night samples. **B. Ferredoxin**. The two targets show a silent single nucleotide polymorphism. The translation of the consensus sequence has its best protein BLAST matches to ferredoxin from several diatom species, the top two are shown in the inset. Its best Pfam match is also ferredoxin (Fer2 = 2Fe-2S iron-sulfur cluster binding domain). Target 1 is found only in France/June samples, while target 2 is found only in Norway/December samples. **C. Fucoxanthin chlorophyll a/c protein (FCP)**. This anchor and its targets are also presented in Figure 5C. At left, the translation of the consensus sequence has its best protein BLAST matches to several diatom species, two that are named as FCP are shown in the inset. The amino acid identity for *Phaeodactylum tricornutum* is 42/44 (95%). The consensus also BLASTs to the *P. tricornutum* genome, nucleotide identity 107/132 (81%) (not shown); this level of mismatch suggests the true origin species is not in the NCBI database. At right, histogram shows total anchor counts for Night are ∼60% lower than for Day, indicating circadian regulation of this gene. All are samples from France in December (the only location/season in which this anchor had both Day and Night representation).

## Note S1. Generality of SPLASH

In this work we focused our experimental results on identifying changes in viral strains and specific examples of RNA-seq analysis. SPLASH’s probabilistic formulation extends much further however, and subsumes a broad range of problems. Many other tasks, some described below, can also be framed under this unifying probabilistic formulation. Thus, SPLASH provides an efficient and general solution to disparate problems in genomics. We outline examples of SPLASH’s predicted application in various biological contexts, highlighting the anchors that would be flagged as significant:

- RNA splicing, even if not alternative or regulated, can be detected by comparing DNA-seq and RNA-seq
  - Examples of predicted significant anchors: sequences upstream of spliced or edited sequences including circular, linear, or gene fusions
- RNA editing can be detected by comparing RNA-seq and DNA-seq
  - Examples of predicted significant anchors: sequences preceding edited sites
- Liquid biopsy – reference free detection of SNPs, centromeric and telomeric expansions with mutations
  - Examples of predicted significant anchors: sequences in telomeres (resp. centromeres) preceding telomeric (resp. centromeric) sequence variants or chromosomal ends (telomeres) in cancer-specific chromosomal fragments
- Detecting MHC allelic diversity
  - Examples of predicted significant anchors: sequences flanking MHC allelic variants
- Detecting disease-specific or person-specific mutations and structural variation in DNA
  - Examples of predicted significant anchors: sequences preceding structural variants or mutations
- Cancer genomics e.g. BCR-ABL fusions
  - Examples of predicted significant anchors: sequences preceding fusion breakpoints
- Transposon or retrotransposon insertions or mobile DNA/RNA
  - Examples of predicted significant anchors: (retro)transposon arms or boundaries of mobile elements
- Adaptation
  - Examples of predicted significant anchors: sequences flanking regions of DNA with time-dependent variation
- Novel virus’ and bacteria; emerging resistance to human immunity or drugs
  - Examples of predicted significant anchors: sequences flanking rapidly evolving or recombined RNA/DNA
- Alternative 3’ UTR use
  - Examples of predicted significant anchors: 3’ sequences with targets including both the poly(A) or poly(U), or adapters in cases of libraries prepared by adapter ligation versus downstream transcript sequence
- Hi-C or any proximity ligation
  - Examples of predicted significant anchors: for Hi-C, DNA sequences with differential proximity to genomic loci as a function of sample; similarly, for other proximity ligation anchors would be predicted when the represented element has differential localization with other elements
- Finding combinatorially controlled genes e.g. V(D)J
  - Examples of predicted significant anchors sequences in the constant, D, J, or V domains

## Generality of SPLASH anchor, target and consensus construction

SPLASH can function on any biological sequence and does not need anchor-target pairs to take the form of gapped *k*-mers, and can take very general forms. For example, one could consider schemes that respect triplet codons: [X_1_X_2_Y_1_][X_3_X_4_Y_2_][X_5_X_6_Y_3_]… where X_i_ are bases in the anchor and Y_i_ are bases in the target, this would focus specifically on variation in the wobble position, the fastest to diverge; similar schemes might be appropriate for mechanisms with known patterns of diversity, such as diversity generating retroelements^79^. X and Y could also be amino acid sequences or other discrete variables considered in molecular biology. SPLASH consensus building can be developed into statistical *de novo* assemblies, including mobile genetic elements with and without circular topologies. Much more general forms of anchor-target pairs (or tensors) can be defined and analyzed, including other univariate or multivariate hash functions on targets or sample identity. SPLASH can also be further developed to analyze higher dimensional relationships between anchors, where statistical inference can be performed on tensors across anchors, targets, and samples. Similarly, hash functions can be optimized under natural maximization criterion, which is the subject of concurrent work. The hash functions can also be generalized to yield new new statistics, optimizing power against different alternatives.

## Note S2. SPLASH statistical inference

### Statistical Inference

In this section we discuss the statistics underlying our *p*-value computation. As discussed, detecting deviations from the global null, where the probability of observing a given target *k*-mer *t*, *R* bases downstream of an anchor *a*, is the same across samples, can be mapped to a statistical test on counts matrices (contingency tables).

### Probabilistic model

Formally, we study the null model posed below.

### Null model

Conditional on anchor *a*, each target is sampled independently from a common vector of (unknown) target probabilities not depending on the sample.

Despite its rich history, the field of statistical inference for contingency tables still has many open problems ^80^. The field’s primary focus has been on either small contingency tables (2×2, e.g. Fisher’s exact test ^81^), high counts settings where a chi-square test yields asymptotically valid *p*-values, or computationally intensive Markov-Chain Monte-Carlo (MCMC) methods. While contingency tables have been widely analyzed in the statistics community ^80, 82, 83^, to our knowledge no existing tests provide closed form, finite-sample valid statistical inference with desired power for the application at hand.

We note that even though we are not aware of directly applicable results, it may be theoretically possible to obtain finite-sample-valid p-values using likelihood ratio tests or a chi-squared statistic. However, even if this were possible, it would not allow for the modularity of our proposed method, where we can a) weight target discrepancies differently as a function of their sequences, to allow for power against different alternatives, b) reweight each sample’s contribution to normalize for unequal sequencing depths, and c) offer biological interpretability in the form of cluster detection and target partitioning. Overall, the statistics we develop for SPLASH are extremely flexible. Ongoing work is focused on further optimizing this general procedure, including application specific tuning of the functions *f* and robustification of the statistic against biological and technical noise.

### Test intuition

The test statistic used by SPLASH can be thought of as using a vector *f* to partition the rows (targets) of the contingency table into two groups, assigning them 1 and 0 respectively. Then, for each sample, we compute the expected value of its target (i.e. what fraction of the targets from this sample were assigned a 1). We construct a per sample score as the difference between its target expected value and the global target expected value (with respect to the target distribution across all samples), and scale this difference by the square root of the number of observations of this anchor. A vector *c* assigns each sample a scalar value, and the final test statistic is computed as the *c*-weighted sum of these per-sample scores. Due to the linear structure of the test statistic, finite-sample *p*-value bounds are possible through classical concentration inequalities.

For fixed vectors *f* and *c*, there are many alternatives that SPLASH would not have power against. Thus, to detect different alternatives, SPLASH utilizes several randomly chosen *f* and *c*, applying Bonferroni correction to the result. In subsequent work, more sophisticated (optimization-based) approaches to computing improved *f* and *c* have been developed, leveraging a linear algebraic perspective on the test statistic. Additional generalizations of *f* and *c* may be of interest.

### *p*-value computation

SPLASH’s *p*-value computation is performed independently on each anchor, and so statistical inference can be performed in parallel across all anchors. Our test statistic is based on a linear combination of row and column counts, giving valid false discovery rate (FDR) controlled *q*-values by classical concentration inequalities and multiple hypothesis correction (Figure S1A). To formalize our notation, we define *D_j,k_* as the sequence identity of the *k*-th target observed for the *j*-th sample. This ordering with respect to *k* that we assign is for analysis purposes only, it has no relation to the order in which targets are observed in the actual FASTQ files (can be thought of as randomly permuting the order in which we observe the targets). Under the null model, each *D_j,k_* is then an independent draw from the common target distribution.

SPLASH’s test statistic poses many exciting research directions, which we explore in a separate statistical work^55^. To construct *p*-values, we first estimate the expectation (unconditional on sample identity) of *f*(*D_j,k_*) as µ̂ by collapsing across samples. Next, we aggregate *f*(*D_j,k_*) across only sample *j* to compute µ̂_*j*_, constructing *S_j_* as the difference between the these two, normalizing by 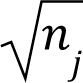 to ensure that each *S_j_* will have essentially constant variance (up to the correlation between µ̂, µ̂_*j*_). This is performed as below:

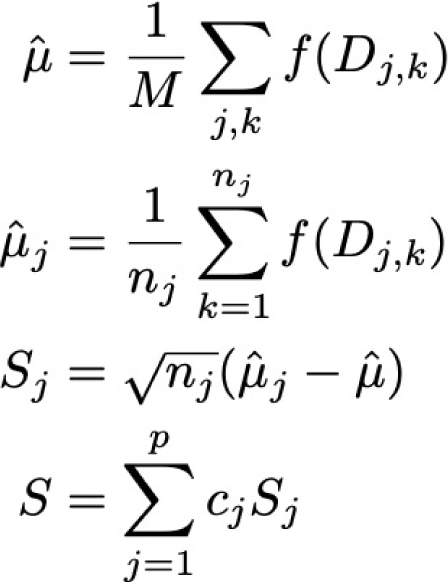

We see that *S_j_* is a signed measure of how different the target distribution of sample *j* is from the table average, when viewed under the expectation with respect to *f*. This function *f* is critical to obtain good statistical guarantees, and the choice of *f* determines the direction of statistical power, such as power to detect SNPs versus alternative splicing or other events. In this work we design a general probabilistic solution, utilizing several random functions *f* which take value 0 or 1 on targets, independently and with equal probability. In order to increase the probability that SPLASH identifies anchors with significant variation, several (*K* = 10 by default) random functions are utilized for each anchor, though more may be desired depending on the application.

After constructing these signed anchor-sample scores, they need to be reduced to a scalar valued test-statistic. Consider first the case where we are given sample metadata, i.e. we know that our samples come from two groups, and we want our test to detect whether the target distribution differs between the two groups. One natural way of performing such a test is to first aggregate the anchor-sample scores over each group, and then compute the difference between these group aggregates.

We formalize this by assigning a scalar *c_j_* to each sample, where in this two group comparison with metadata *c_j_* = ±1 encodes the sample’s identity, and construct the anchor statistic *S* as the inner product between the vector of *c_j_*’s and the anchor-sample scores. This statistic will have high expected magnitude if there is significant variation in target distribution between the two groups.

In many biologically important applications however, cell-type metadata is not available. In these cases, SPLASH detects heterogeneity within a dataset by performing several (*L* = 50 by default) random splits of the samples into two groups. For each of these *L* splits SPLASH assigns *c_j_* = ±1 independently and with equal probability for each sample, computes the test statistic for each split, and selects the split yielding the smallest *p*-value.

We now investigate the statistical properties of *S*. First, observe that *S* has mean 0 under the null hypothesis. This allows us to bound the probability that the random variable *S* is larger than our observed anchor statistic as follows. Since *f* and *c* are fixed, and are independent of the data, we have that since *f*(*D_j,k_*) are bounded between 0 and 1 we can apply Hoeffding’s inequality for bounded random variables. Defining µ as the expectation with respect to the common underlying distribution of *f*(*D_j,k_*) (unknown), we center our random variables by subtracting the sample mean µ, our estimate of the true mean µ. Standard bounds can now be applied to decompose this deviation probability into two intuitive and standard terms:

1) the probability that the statistic 𝑆̃, constructed with unavailable knowledge of the true µ, is large

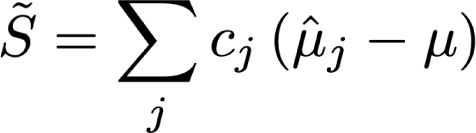

2) the probability that µ̂ is far from µ. Following this approach, we have that

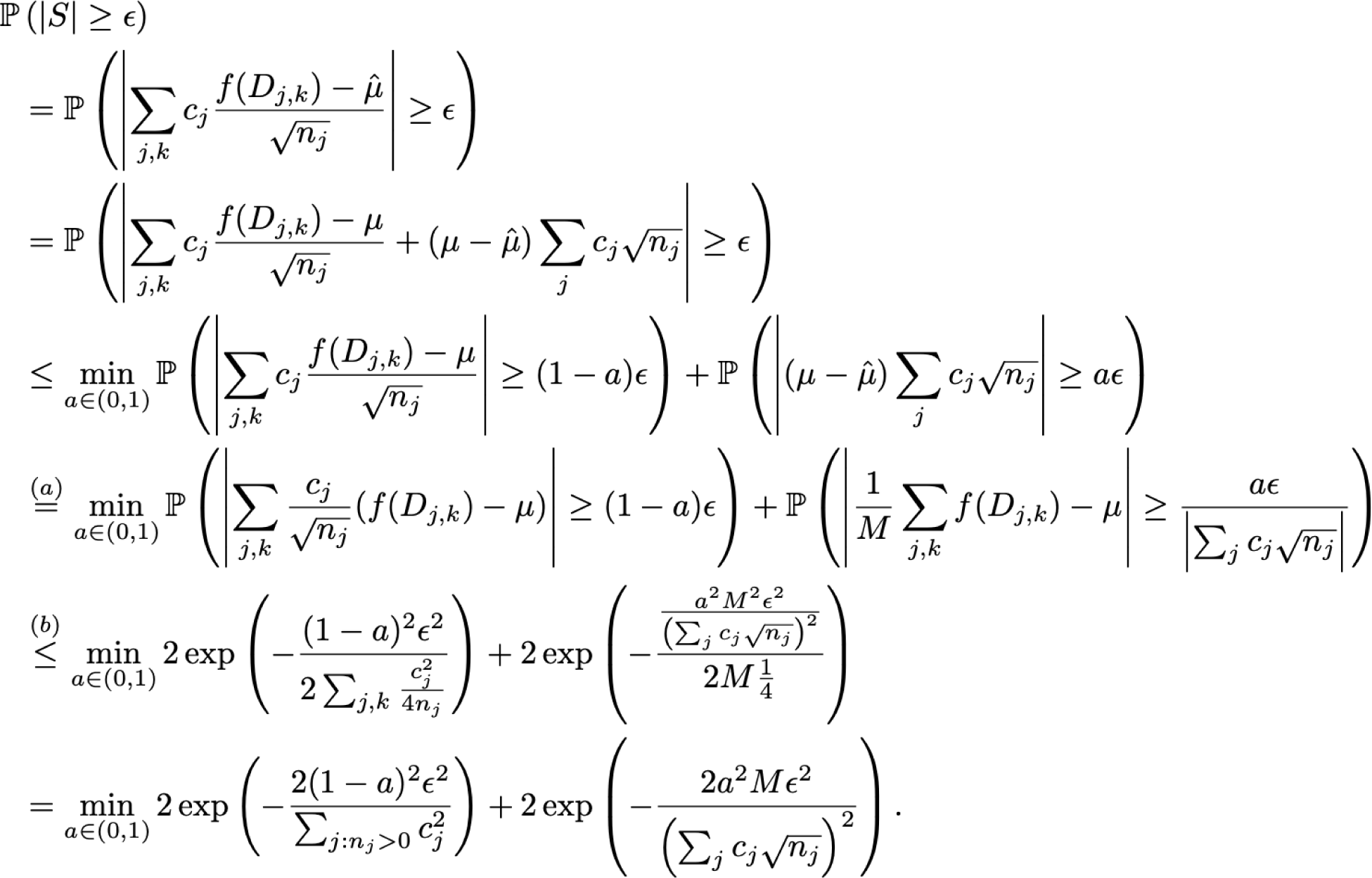

where (a) comes from the assumption that the sum in the denominator of the second term is nonzero, as otherwise this second term is 0 and we can essentially set *a* = 0. (b) utilizes Hoeffding’s inequality on each of these two terms. We can easily optimize this bound over *a* to within a factor of two of optimum by equating the two terms (as one is increasing in *a* and the other is decreasing), which is achieved when

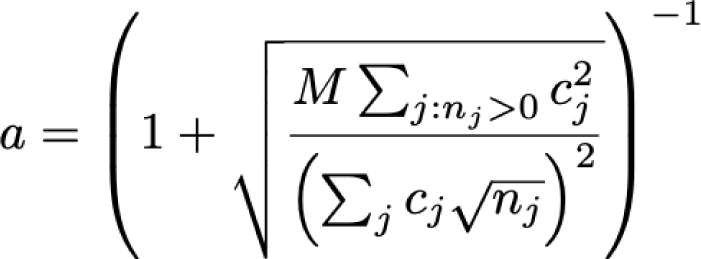

Thus, for an observed value of our test statistic *S*, we construct SPLASH’s finite-sample valid *p*-values as

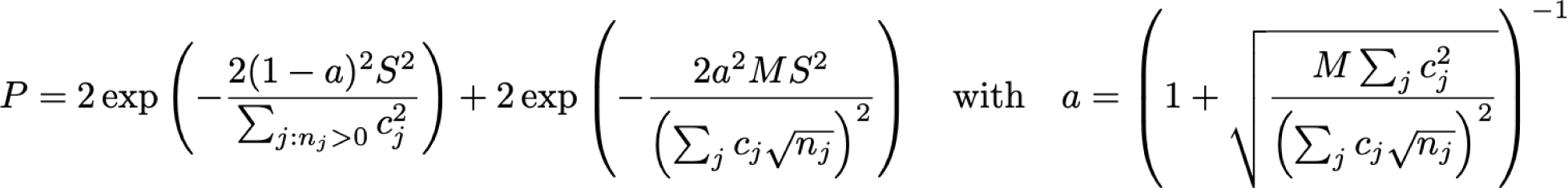

### *q*-value computation

*q*-values are computed using Benjamini Yekutieli correction ^58^ as

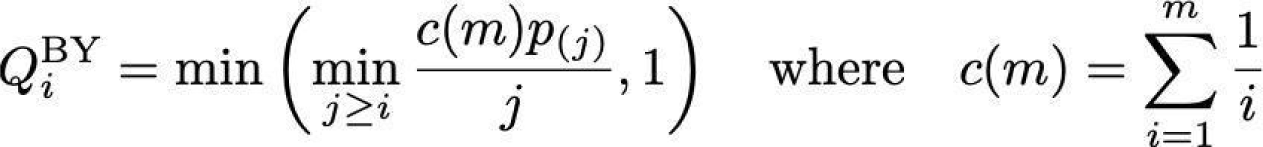

which enables SPLASH to control the FDR of the reported significant anchors.

### Effect size

SPLASH provides a measure of effect size when the *c_j_*’s used are ±1, to allow for prioritization of anchors with fewer counts but large inter-sample differences in target distributions. Effect size is calculated based on the split *c* and function *f* that yield the most significant SPLASH *p*-value. Fixing these, the effect size is computed as the difference between the mean over targets with respect to *f* across those samples with *c* = +1, and the mean over targets (with respect to *f*) across those samples with *c* = -1. This effect size is bounded between 0 and 1, with 0 indicating no effect (target distributions are identical when aggregated within each group), and 1 indicating disjoint supports. Defining *A_+_* as the set of *j* where *c_j_* > 0, and *A_-_* as the set of *j* where *c_j_* < 0 (generalizing beyond the case of *c_j_* = ±1), this is formally computed as:

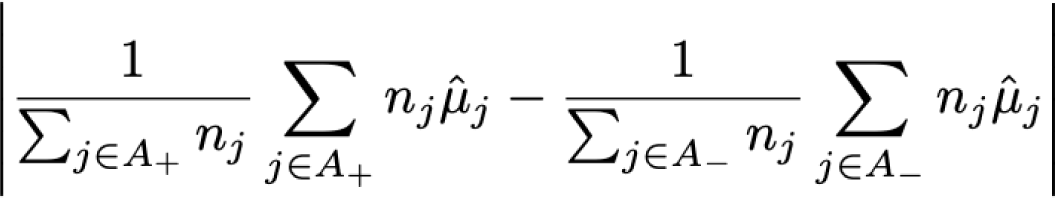

In this simple case of *c_j_* = ±1 and {0,1} valued *f*, this is simply a projection of the *T*×*p* table to a 2×2 table. Even considering more general *f*, there is an easy to understand alternative that SPLASH is designed to have power against. The effect size should be thought of under the alternative hypothesis where the columns follow multinomial distributions with probability vector *p_1_* or probability vector *p_2_*, depending on the group identity *c_j_*. The effect size we compute can be thought of in this scenario as measuring the difference between the expectation of *f* under *p_1_* and *p_2_*. In the case of maximizing the effect size over all possible {0,1}-valued *f*, the effect size will be equal to the total variation distance between the empirical distributions of the group *c_j_* = +1 and *c_j_* = -1. Thus, the effect size will be 1 if and only if the two sample groups partition targets into 2 disjoint sets on which the function *f* takes opposite values, as to be expected from the total variation distance interpretation (Figure S1B). This *f* will place a value of 1 on targets where the empirical frequency of the +1 group *p_1,t_* is larger than that of the -1 group *p_2,t_*. Since *p_1_* and *p_2_* are probability distributions, this ends up being exactly the total variation distance between them (i.e. half the vector *ℓ_1_* distance). Note that we can also consider a signed variant of this effect size measurement, where if we restrict ourselves to the same *c* and *f* for several anchors, the effect size sign gives us additional information about the direction of the effect.

## Note S3. SPLASH is robust to parameter choices and effective without metadata

### SPLASH is robust to parameter choices

We give examples of how choices of *k*, *R*, and tiling length impact results in France SARS-CoV-2 data as follows, showing that SPLASH yields similar results for a range of parameter choices. Default parameters shown in bold: we tested *k* = [25, **27**, 30]; Tile = [3, **5**, 7]; Lookahead = [0, 15, **23**]. For *k* = 25, 94.4% of anchors with default parameters contain at least one of the K=25 anchors as a substring. For *k* = 30, 93.8% of anchors with *k* = 30 contain at least one of the anchors with default parameters a substring. For tile size of 3, 85% of the anchors from the default run can be found in the significant anchors of tile size of 3. For tile size of 7, 85% of the anchors from the default run can be found in the significant anchors of tile size of 5. For lookahead distance of 0, 37% of the anchors from the default run can be found in the significant anchors of tile size of 3; for lookahead distance of 15, 76% of the anchors from the default run can be found in the significant anchors. Overall, as tile size decreases, anchor calls increase (4715, 5522, 7891 for [7, 5, 3] respectively). As *k* varies, anchor calls stay essentially the same (5875, 5522, and 5958 for *k* = [25, 27, 30] respectively). Finally, for lookahead distance, the total number of calls decrease as lookahead distance increases (13239, 8295, 5522 for *R* = [0, 15, 23] respectively).

### SPLASH is effective without metadata

As discussed, SPLASH can be run without any metadata. For the HLCA dataset, when run on the two donors without metadata, SPLASH calls 6287 anchors (2269 genes) as opposed to the 3439 anchors (1384 genes) called with metadata for donor 1. Filtering for genes hit by more than two anchors, SPLASH’s metadata free approach calls >94% of the genes called by the metadata-based approach (Figure S3). For donor 2, SPLASH calls 5619 anchors (1844) genes without any metadata as opposed to the 3775 anchors (1125) genes called with metadata. Filtering for genes hit by more than two anchors, SPLASH’s metadata free approach calls >90% of the genes called by the metadata-based approach, increasing to >94% for those genes hit by at least 3 anchors.

## Note S4. Lemur lambda light chain and surrogate light chains found by SPLASH

As detailed in Methods, we attempted to identify light chain variable regions in lemur B cells where the BASIC pipeline^38^ could not. We were successful in identifying a full variable region by extension (using ′grep′ of raw reads) of an initial seed consensus sequence found by SPLASH. Below we give the sequence identified, and the NCBI IgBlast report for it. IgBlast uses human Ig reference sequences (lemur Ig regions are not well annotated) so it is uncertain which differences are due to lemur and which to hypermutation, however there appears to be a high mutational load in this variable region, which may be why BASIC could not identify it.

This is the cell that had a full V-region:

## cell: MAA001400_B109012_I1_S193

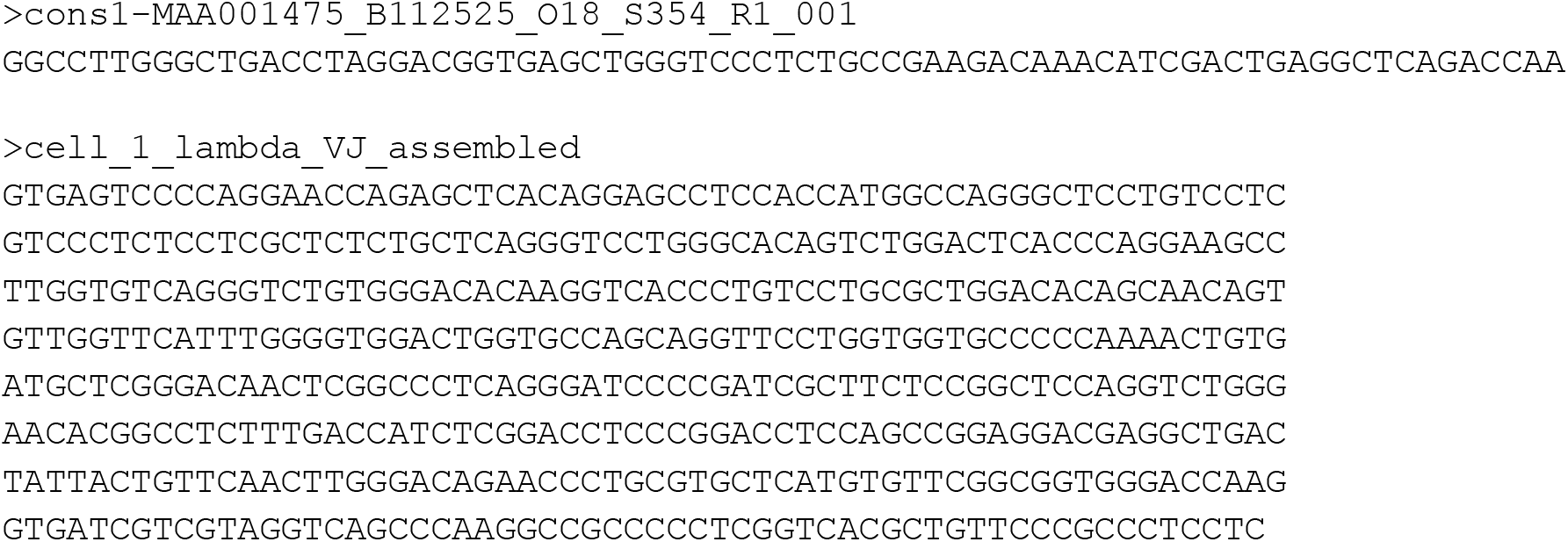

### Protein translation

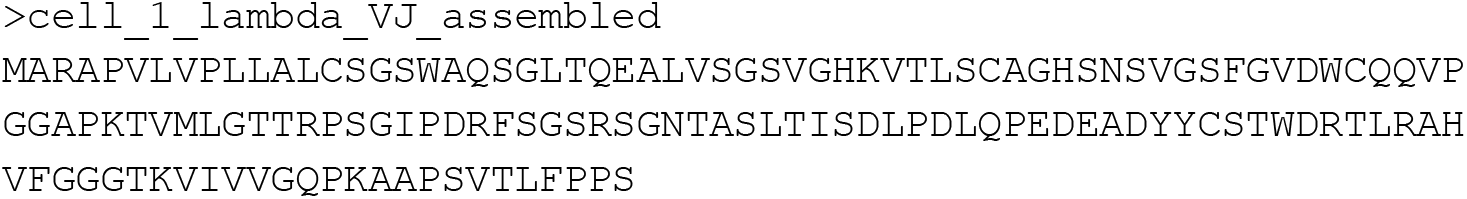

Below is the NCBI IgBlast report for the full V-region (FASTA sequence given at the end.) Coding differences from germline gene are shown in magenta.

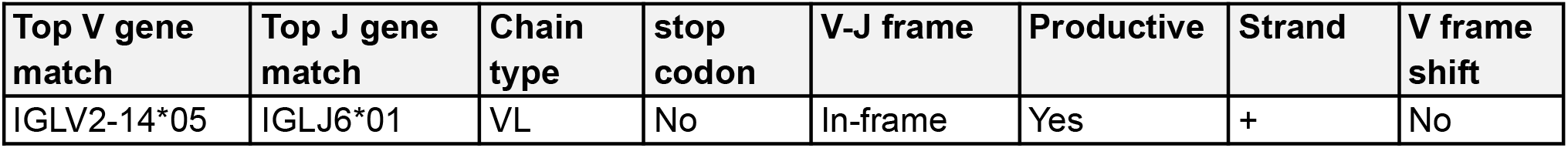

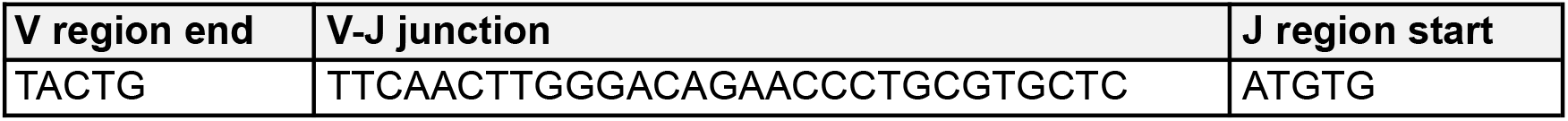

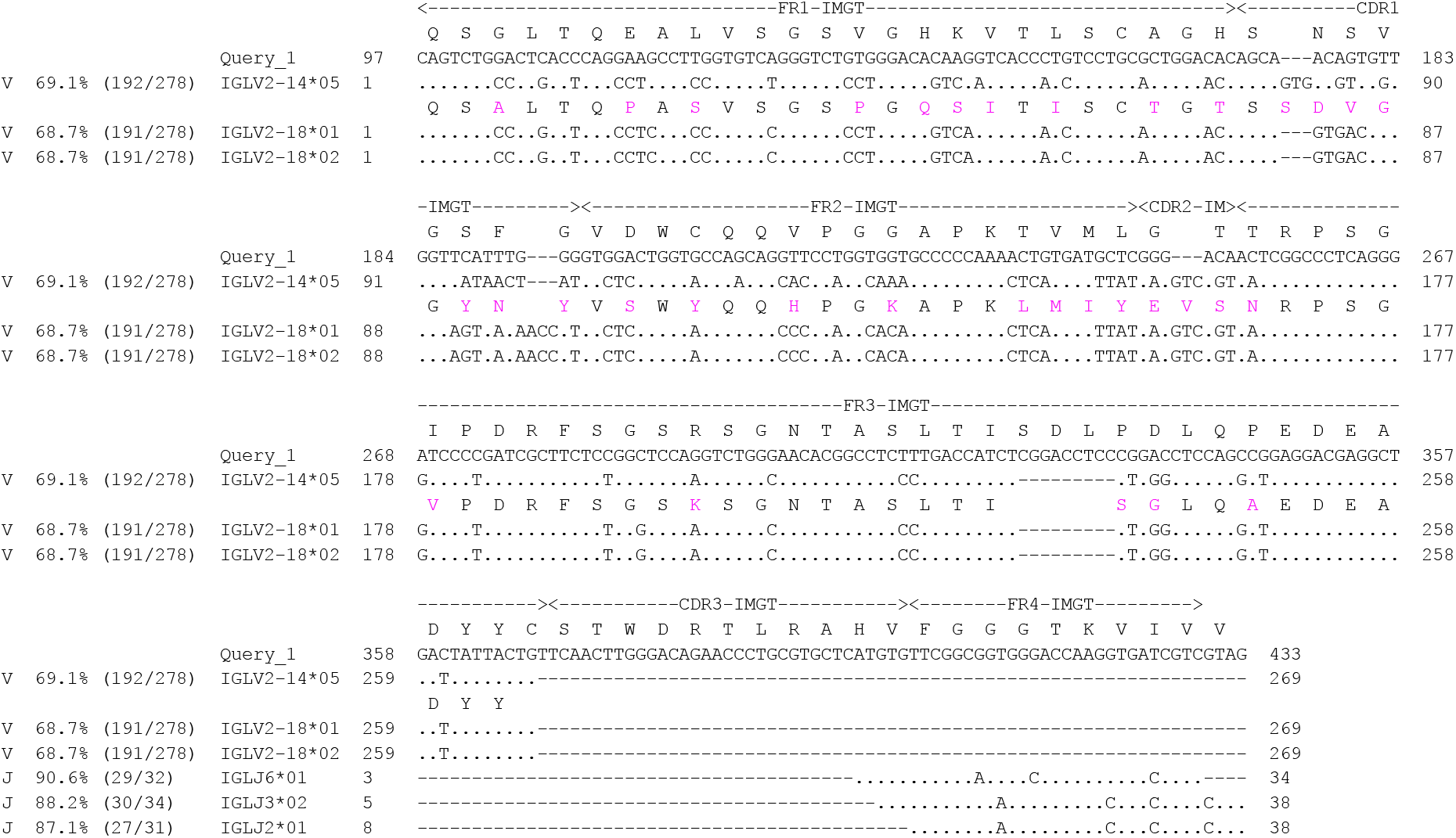

Here are the BLAST matches of consensuses from two cells to the unique region of IGLL1/5, one of the surrogate light chains. This suggests that these two cells have not yet rearranged their light chain.

### cell: MAA001475_B112525_C4_S52

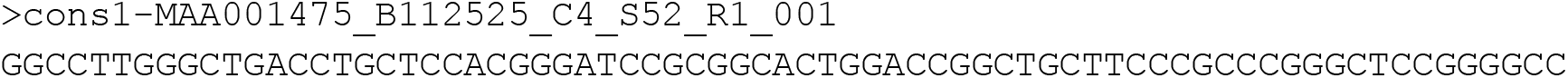

Consensus sequence above is the original strand, but BLAST was done with its reverse-complement, the sense strand (this is the Query). The alignment indicates a 1-nt insertion in the consensus relative to IGLL5 (which would cause a frameshift).

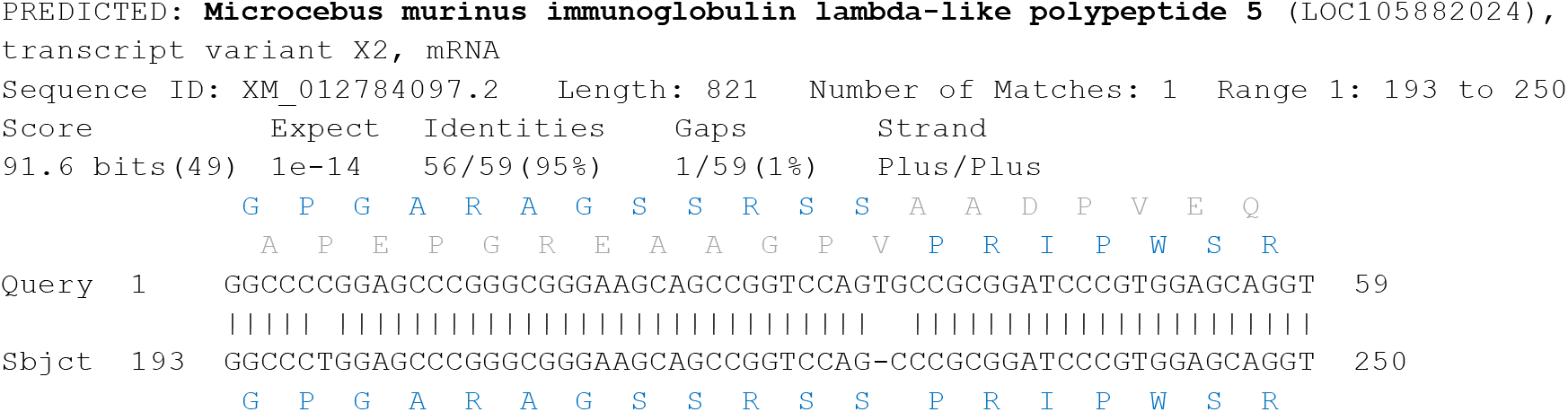

Protein comparison of the mouse lemur IGLL5 above to human IGLL5 (where domains are identified) shows that the above region clearly lies in the IGLL5 N-terminal unique region. The matching part above is shown in blue.

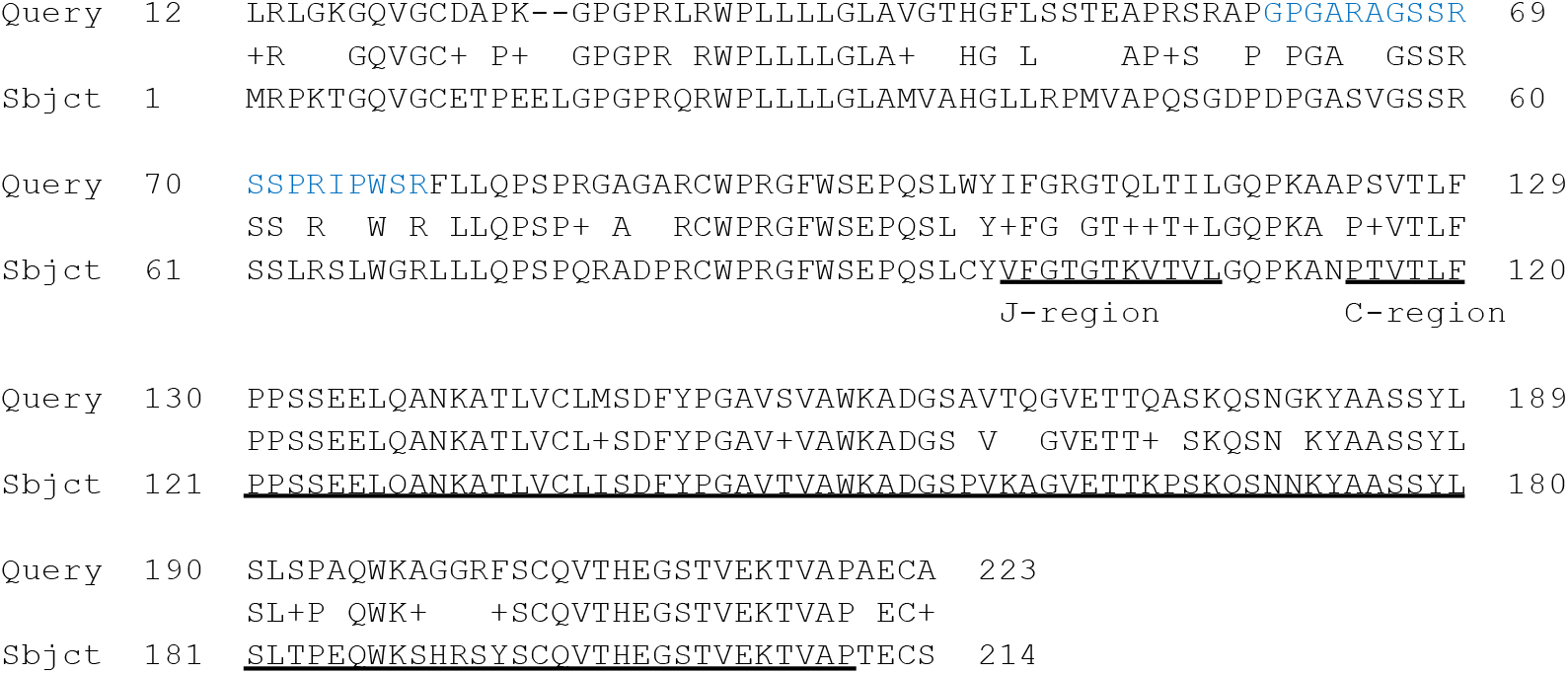

### cell: MAA001475_B112525_O18_S354

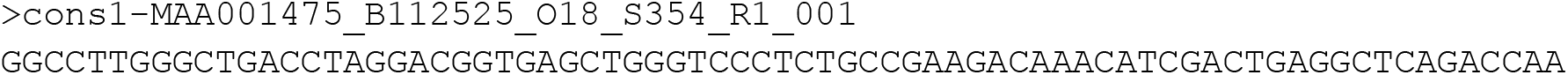

Consensus sequence above is the original strand, but BLAST was done with its reverse-complement, the sense strand (this is the Query). Here the match is to human IGLL5. The consensus includes regions upstream of the J-region, so part of the IGLL5-unique region.

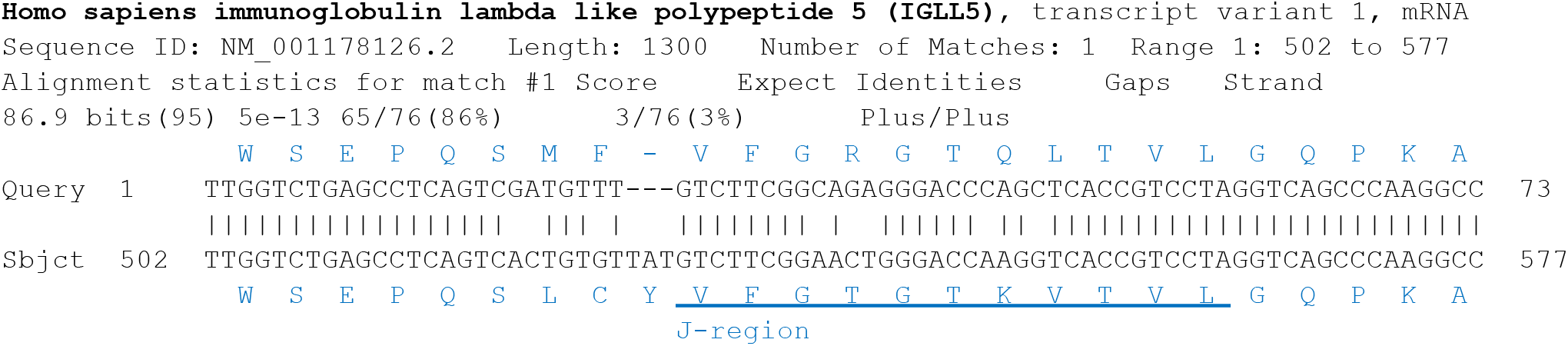

## Note S5. Octopus and eelgrass analyses, additional notes

### Octopus bimaculoides Myo-VIIa

We give more detailed alignment information for Myo-VIIa anchor and targets, to document the unannotated alternative first exon “1b” (that is expressed specifically in statocyst tissue) and the absence of sequence matching exon 2 in the *O. bimaculoides* genome. Note that we do not have information on the start of exon “1b”. We use reverse-complements of the anchor and targets (the sense strand).

### Alignment to O. sinensis genome

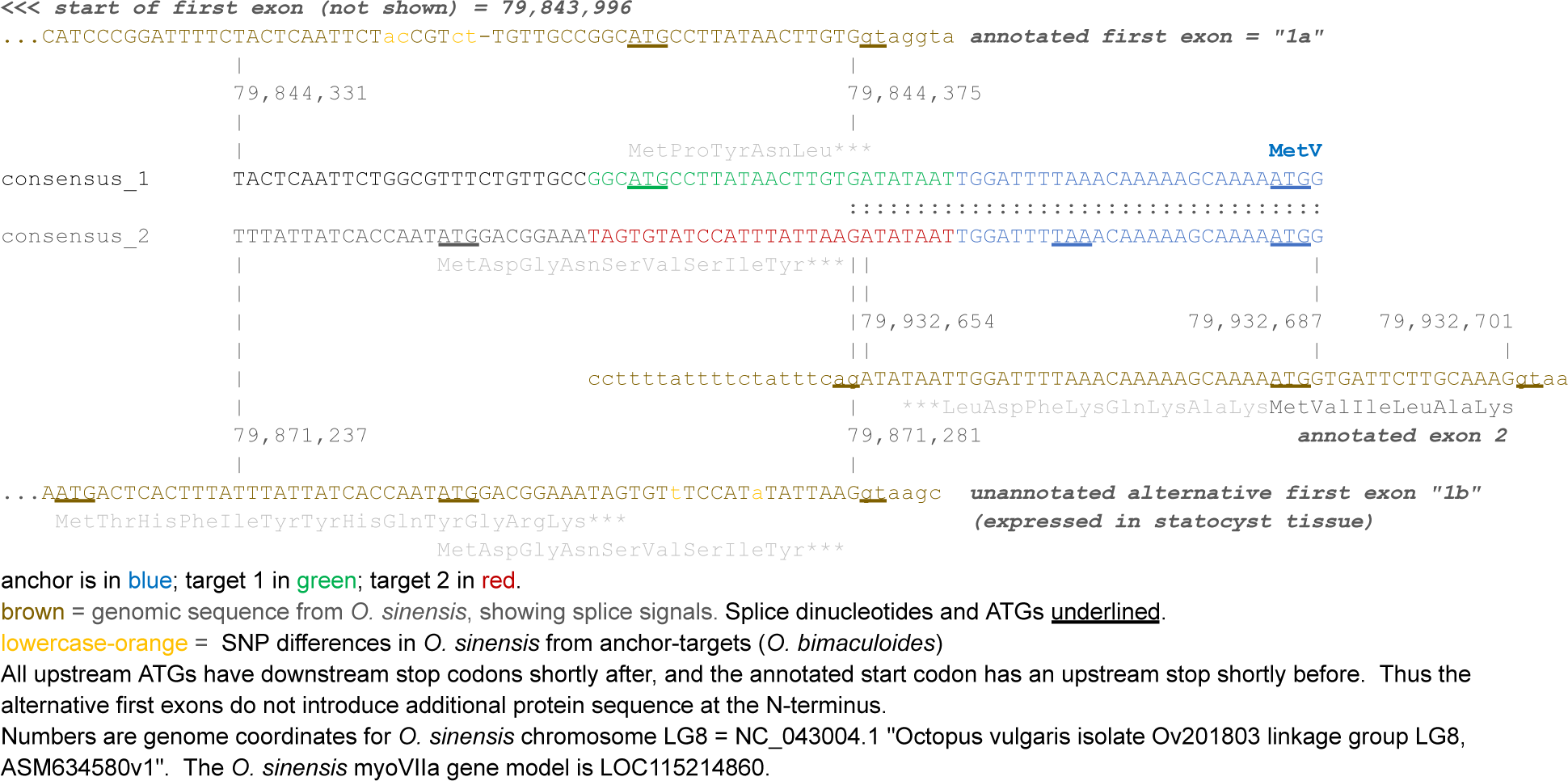

### Alignment to *O. bimaculoides* genome

There is no full perfect match for the relevant exon 2 portion nor for the anchor (underlined sequence) (ATATAAT<u>TGGATTTTAAACAAAAAGCAAAAATGG</u>) in *O. bimaculoides*. Sequences matching exons 1a and 1b are present in the *O. bimaculoides* genome. The splice donor for *O. sinensis* myoVIIa exon 1a matches perfectly to an *O. bimaculoides* splice donor in an annotated noncoding RNA XR_008264717.1 (gene LOC128248543) located upstream of the annotated *O. bimaculoides* myoVIIa transcript.

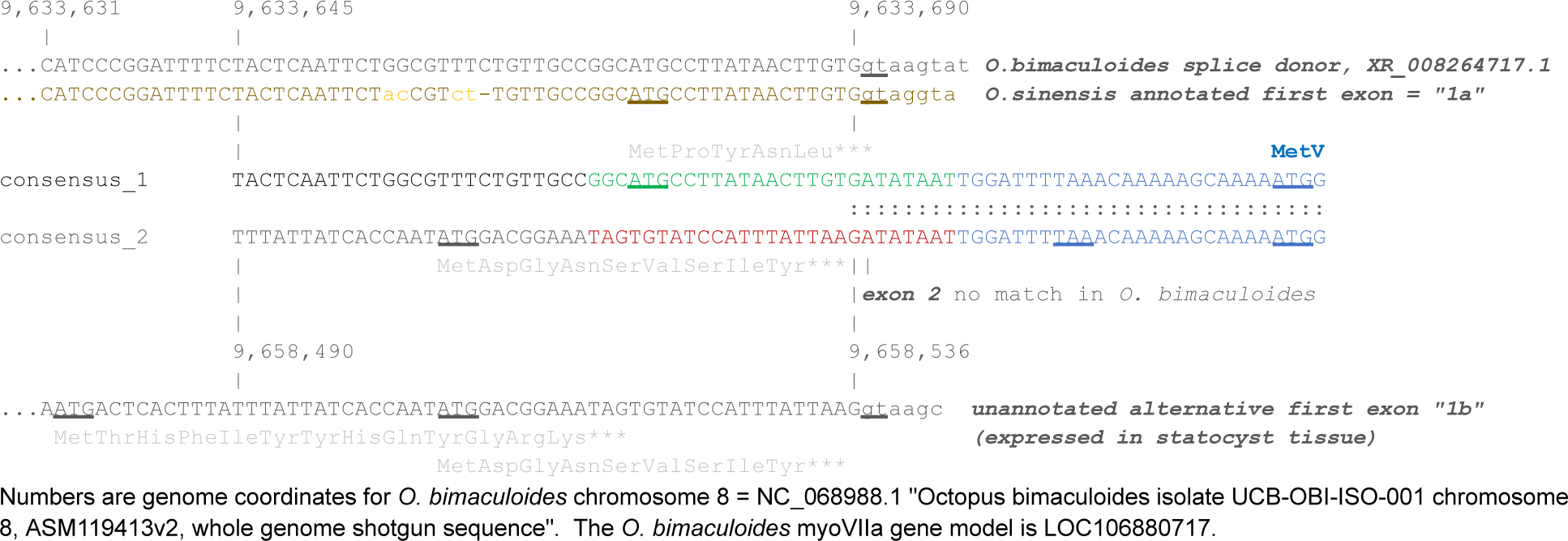

There appears to be an assembly issue for the *O. sinensis* Myo-VIIa gene on chromosome LG8 that matches our anchor-targets. There are several *O. sinensis* genes annotated with myosin-VIIa in the genome. We list them below, including their protein domain content.

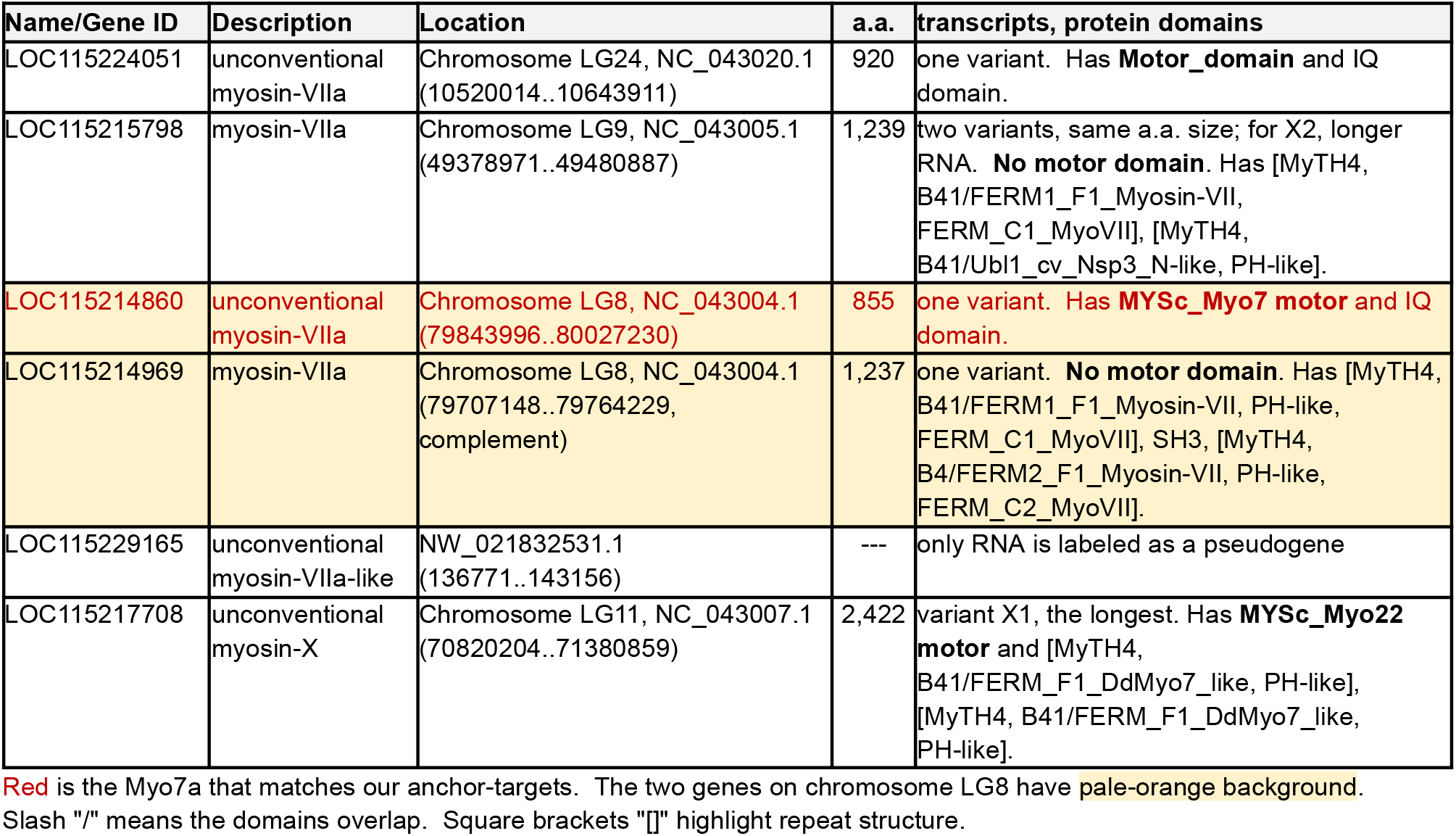

Neither of the two myo-VIIa genes on Ch. LG8 has a full complement of protein domains: LOC115214860 has the N-terminal myosin motor domain, but lacks the tail domains. LOC115214969 has all the tail domains but lacks the motor domain. The two genes are adjacent on the chromosome, but are in head-to-head orientation, as shown in this graphic screenshot from NCBI (LOC115214860 is the red arrow, LOC115214969 has been marked with a red box):

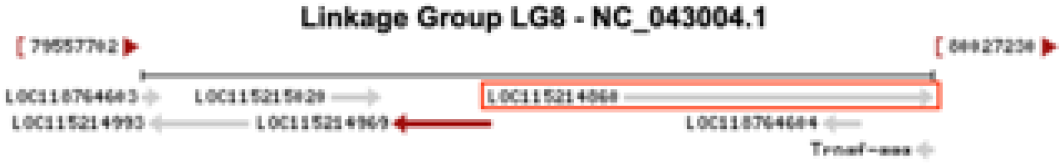

If LOC115214860 was inverted, then all domains would be present in the correct linear order.

## *Zostera marina* (eelgrass) NADPH quinone oxidoreductase subunit L (NdhL) intron retention

We have confirmed the intron retention event by sequence extension of target 4 from raw reads to reach the end of exon 2 (data not shown).

We show here the predicted translation of the intron retention isoform of NdhL (target 4 of Figure 5D). It causes a frameshift and termination shortly after the end of exon 3 (where the other targets are located). The anchor is in exon 4.

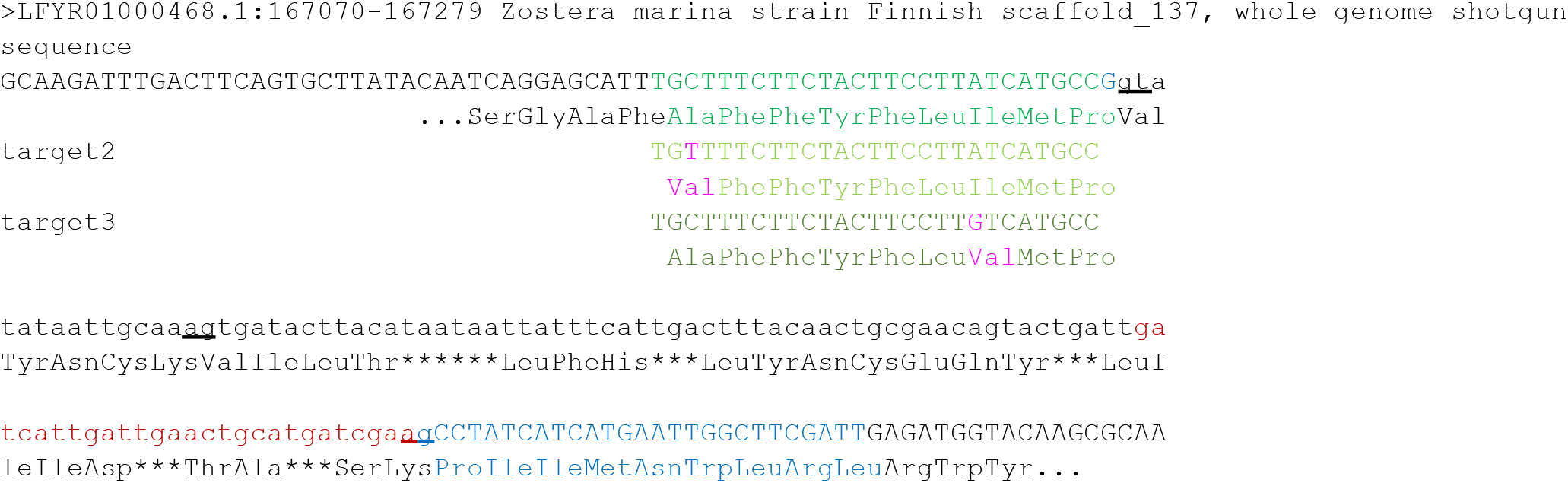

The frameshift and termination occur within the second transmembrane domain of the protein. (There is a third transmembrane domain, but it is not predicted by all programs.) The topology below was predicted by CCTOP (https://cctop.ttk.hu/)

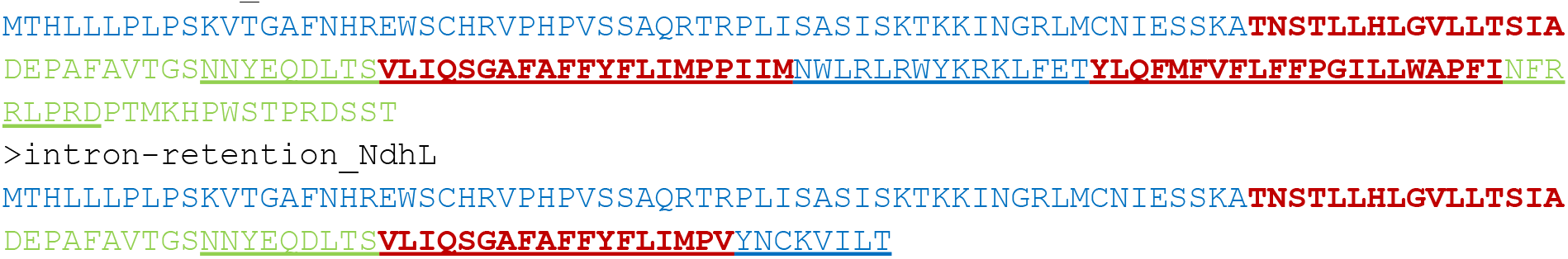

## Note S6. SPLASH runs on a laptop

### Computational benchmarking for SPLASH

SPLASH is computationally much more efficient than other approaches, due to its use of k-mers rather than reference alignment, and its closed-form statistics obviating compute-intensive significance testing. SPLASH is implemented as a fully containerized and parallelized workflow that requires only the FASTQ read files and no parameter tuning by the user. We ran SPLASH on a 2015 Intel laptop with an Intel® Core™ i7-6500U CPU @ 2.50GHz processor, generating significance calls for single cell RNA-seq totaling over 10 million reads in only 1 hour 45 min. When performed on a compute cluster, the same analysis is completed in an average of 22.8 minutes with 750 MB of memory for 10 million reads.

### Timing for SS2

Because code was run on a server with dynamic memory, we report summary statistics as follows. For the steps parallelized by FASTQ file, such as anchor and target retrieval, total time for dataset run, as reported by Nextflow, was parsed per cell. Thus, the average time per cell is reported. For the steps parallelized by 64 files (*q*-value calculations), total extracted times were summed and divided by number of cells. For steps that consisted of aggregating files, total run time was divided by number of cells. Thus, the total time and memory should be multiplied by the total number of cells to achieve an estimate of the pipeline time for this dataset.

### Laptop analysis details

#### Laptop specs

An Intel® Core™ i7-6500U CPU @ 2.50GHz (launched in 2015)

2 cores, total of 4 threads, 3 of which SPLASH was allowed to use. 8 GB DDR3 RAM

SODIMM DDR3 Synchronous 1600 MHz (0.6 ns)

### Laptop analysis dataset

Ten B and T cells from donor 2 blood sequenced by Smart-Seq2 were used for the laptop benchmarking. These files totalled 43,870,027 reads, averaging 4.3M reads per cell. The fastq files for the Tabula Sapiens data were downloaded from

https://tabula-sapiens-portal.ds.czbiohub.org/. Files used:

TSP2_Blood_NA_SS2_B114581_B133053_Lymphocytes_A13_S73_R1_001.fastq.gz

TSP2_Blood_NA_SS2_B114581_B133053_Lymphocytes_A18_S78_R1_001.fastq.gz

TSP2_Blood_NA_SS2_B114581_B133053_Lymphocytes_A19_S79_R1_001.fastq.gz

TSP2_Blood_NA_SS2_B114581_B133053_Lymphocytes_A21_S81_R1_001.fastq.gz

TSP2_Blood_NA_SS2_B114581_B133053_Lymphocytes_A3_S63_R1_001.fastq.gz

TSP2_Blood_NA_SS2_B114581_B133053_Lymphocytes_A5_S65_R1_001.fastq.gz

TSP2_Blood_NA_SS2_B114581_B133053_Lymphocytes_A6_S66_R1_001.fastq.gz

TSP2_Blood_NA_SS2_B114581_B133053_Lymphocytes_A8_S68_R1_001.fastq.gz

TSP2_Blood_NA_SS2_B114581_B133053_Lymphocytes_A9_S69_R1_001.fastq.gz

TSP2_Blood_NA_SS2_B114581_B133053_Lymphocytes_B10_S94_R1_001.fastq.gz

## Note S7. Anchor and target sequences, *q*-values, and binomial *p*-values

### Binomial *p*-value bound computation for plots depicting target fraction abundance

We provide *p*-values to quantify the visually striking nature of the plots depicting fraction abundance per a specific target (target 1 by default). Under a null model, where all samples are expressing this target with the same probability, the number of times each sample expresses target 1 is binomial(*n_j_*,p), for common p. As seen from the plots, many samples exhibit highly deviating occurrences (number of observations of target 1 that are far from the expected p*n_j_*. The *p*-values we provide to this effect are not used in any SPLASH discovery or analysis, and are just used to quantify the visuals.

*p*-values are constructed as follows: first, we compute p, the average occurrence of target 1 for this anchor (sum of counts of observations of target 1 divided by the total number of observations). Then, for all possible *n_j_*, we compute 1% and 99% quantiles (confidence bounds) for a binomial distribution with *n_j_* trials and heads probability p. If the fraction of target 1 in each sample was independent of sample identity, and were indeed binomially distributed, then each sample would have at least a 98% probability of falling within this confidence interval. Thus, we compute our test statistic *X* as the number of samples that fall outside of the [1,99] quantiles, and compute as our *p*-value the probability that a binomial random variable Bin(*m*,*q*)≥ *X*, where with *m* = number of samples and *q* = .02.

While intuitive, the above analysis is loose. Firstly, since binomials are discrete distributions, we will rarely be able to compute exact 1% and 99% quantiles. Thus, the probability that for any given *n_j_* a sample will fall outside of the [1,99] quantiles, which we denote *q_j_*, is almost always substantially less than .02. The true distribution of X is then poisson binomial, with this vector of probabilities (all at most .02), one for each sample. However, as this *p*-value is numerically difficult to compute, we bound this

*p*-value as the probability that Bin(*m*,*q’*)≥X, where m = number of samples with *q’*=max_j_ *q_j_*, where *q’* ≤.02.

### Anchor and target sequences, with *q*-values and binomial *p*-values

Targets are numbered by decreasing abundance, unless otherwise stated.

*q*-values are the BY-corrected *p*-values output by SPLASH, as detailed in Note S2. Binomial *p*-value calculations are described above, and are with respect to target 1, unless otherwise stated.

### SARS-CoV-2 mutation K417N (Figure 2A)

*q*-value: 9.4e-05

binomial *p*-value: 6.4e-07

>anchor ATTCATTTGTAATTAGAGGTGATGAAG

>target_1_Delta ACTGGAAAGATTGCTGATTATAATTAT

>target_2_K417N_Omicron ACTGGAAATATTGCTGATTATAATTAT

### SARS-CoV-2 mutations V213G, NL211I, R214REPE (Figure 2B)

*q*-value: 8.3e-08 binomial *p*-value: 1e-13

>anchor TTTAAGAATATTGATGGTTATTTTAAA

>target_1_Delta TAATTTAGTGCGTGATCTCCCTCAGGG

>target_2_V213G_BA.2 TAATTTAGGGCGTGATCTCCCTCAGGG

>target_3_NL211I-R214REPE_BA.1 TATAGTGCGTGAGCCAGAAGATCTCCC

SARS-CoV-2 mutations P681R, N679K, P681H (Figure 2C)

*q*-value: 1.2e-04

binomial *p*-value: 4.9e-12

(reverse-complements are shown in Figure 1C)

>anchor GTGACATAGTGTAGGCAATGATGGATT

>target_1_P681R_Delta (abundance order = 1) CGACGAGAATTAGTCTGAGTCTGATAA

>target_2_P681R-Q677H (abundance order = 3) CGACGAGAATTAGTATGAGTCTGATAA

>target_3_P681R-Q677H (abundance order = 4) CGACGAGAATTAGTGTGAGTCTGATAA

>target_4_N679K-P681H_Omicron (abundance order = 2) CGATGAGACTTAGTCTGAGTCTGATAA

### MYL12A / MYL12B (Figure 3A, S4B)

P2 *q*-value: 2.5e-08

P2 binomial *p*-value: 9.9e-37 (with respect to target 2) P3 *q*-value: 2.3E-42

P3 binomial *p*-value: 2.2e-45 (with respect to target 1)

>P2_anchor AAGAGGCCTTCAACATGATTGATCAGA

>P2_target_2_MYL12A TTCATTGGGGAAGAATCCAACTGATGA

>P2_target_1_MYL12B TTCTCTAGGGAAGAATCCCACTGATGC

>P2_consensus_MYL12A_macrophage ACAGAGATGGTTTCATCGACAAGGAAGATTTGCATGATATGCTTGCTTCATTGGGGAAGAATCCAACTGATGAGTATCTAGATGCCATGATGAATGAGGCTCCAGGCCCCATCAATT

>P2_consensus_MYL12B_capillary ACAGAGATGGCTTCATCGACAAGGAAGATTTGCATGATATGCTTGCTTCTCTAGGGAAGAATCCCACTGATGCATACCTTGATGCCATGATGAATGAGGCCCCAGGGCCCATCA

>P3_anchor AAGAGGCCTTCAACATGATTGATCAGA

>P3_target_1_MYL12A GAAGATTTGCATGATATGCTTGCTTCA

>P3_target_2_MYL12B GAAGATTTGCATGATATGCTTGCTTCT

>p3_consensus_MYL12A_macrophage ACAGAGATGGTTTCATCGACAAGGAAGATTTGCATGATATGCTTGCTTCATTGGGGAAGAATCCAACTGATGAGTATCTAGATGCCATGATGAATGAGGCTCCAGGCC

>p3_cons_MYL12B_capillary ACAGAGATGGCTTCATCGACAAGGAAGATTTGCATGATATGCTTGCTTCTCTAGGGAAGAATCCCACTGATGCATACCTTGATGCCATGATGAATGAGGCCCCAGGGCCCATCAATTT

### HLA-DRB1 / HLA-DRB4 (Figure 3B)

P2 *q*-value: 4.0e-10

P2 binomial *p*-value: 2e-17 P3 *q*-value: 1.2e-4

P2 binomial *p*-value: 1.6e-08

(reverse-complements are shown in Figure 3B)

>P2_anchor GGAAGCCACAAGGGAGGACATTTTCTG

>P2_target_1_DRB1 GTGGAAGAATAACTGCCAAGCAGGAAA

>P2_target_2_DRB4 GGAAGAATAAGAGCCAAGTGGGAAAGC

>P2_consensus_DRB1_macrophage GGAAGCCACAAGGGAGGACATTTTCTG

CAGTTGCCGAACCAGTAGCAACCAGGTCCTGAGAAAGCCCTCTCTTGTGGAAGAATAACTGCCAAGCAG GAAAGCTTTTCATTCTGCAAAGCTGGGACAGAAGGTTCTTCCTTGAATGT

>P2_consensus_DRB4_capillary CAGAGTTGCTGAACCAGTAACAACCTGGTCCTGACAAAGCTCTTGTGGAAGAATAAGAGCCAAGTGGGAAAGCTTTTCATCTTGCAAAGCTGGGGCAGAAGGTTCTTCCTTGAATGT

>P3_anchor (same sequence as P2_anchor) GGAAGCCACAAGGGAGGACATTTTCTG

>P3_target_1_DRB1 AGGTCCTGAGAAAGCCCTCTCTTGTGG

>P3_target_3_DRB4 CCTGGTCCTGACAAAGCTCTTGTGGAA

>P3_consensus_DRB1_macrophage CAGTTGCTGAACCAGTAGCAACCAGGTCCTGAGAAAGCCCTCTCTTGTGGAAGAATAACAGCCAGGAGGGAAAGCTTTTCATCCTGCAAAGCTGGGGCAGAAAGTTCTTCT

>P3_consensus_DRB4_capillary GGAAGCCACAAGGGAGGACATTTTCTG

CAGAGTTGCTGAACCAGTAACAACCTGGTCCTGACAAAGCTCTTGTGGAAGAATAAGAGCCAAGTGGGA AAGCCTTTCATCTTGCAAAGCTGGGGCAGAAGGTTCTTCCTTGA

### HLA-DPA1 / HLA-DPB1 (Figure 3C, S4C)

P3 *q*-value: 7.9e-22

P3 binomial *p*-value: 9.15e-18

(anchor as given here is sense strand for DPA1, antisense strand for DPB1)

>P3_anchor AGATGTATCTCTCCAGGAAGCGCTGTG

>P3_target_1_DPA1 TGCCGTCCCTGGAAAAGGTGAATCCCA

>P3_target_2_DPB1 TGCCGTCCCTGGAAAAGGTAATTCTCT

>P3_consensus_DPB1_macrophage TCCCATTAAACGCGTAGCATTCCTGCCGTCCCTGGAAAAGGTAATTCTCTGGAGTGGCCCTGCCCTGGACCACAGATGTGAGCAGCACCATCAGTAACGCCGTCAGAGCCACT

>P3_consensus_DPA1_capillary TCCCATTAAACGCGTAGCATTCCTGCCGTCCCTGGAAAAGGTGAATCCCAGCCATGCTGATTCCTCTCCACCCATTTCCAGTGCTAGAGGCCCACAGTTTCAGTCTCATCTGC

### HLA-B (Figure 3D, S4D)

*q*-value: 2.7e-05

binomial *p*-value: 1.7e-25

>anchor TTGGGACCGGAACACACAGATCTTCAA

>target_1_HLA-B AGAGCCTGCGGAACCTGCGCGGCTACT

>target_2_HLA-B AGAACCTGCGGATCGCGCTCCGCTACT

>consensus_1_HLA-B

TTGGGACCGGAACACACAGATCTTCAAGACCAACACACAGACTGACCGAGAGAGCCTGCGGAACCTGCGCGGCTACTACAACCAGAGCGAGGCCGGGTC

>consensus_2_HLA-B TTGGGACCGGAACACACAGATCTTCAAGACCAACACACAGACTTACCGAGAGAACCTGCGGATCGCGCTCCGCTACTACAACCAGAGCGAGGCCGGGTC

### human Ig-kappa C-region (Figure 4B)

*q*-value = 1.6E-35

>anchor TGGCGGGAAGATGAAGACAGATGGTGC

>Targ0 GCTTGGTCCCCTGGCCAAAAGTCCCGG

>Targ1 GCTTGGTCCCCTGGCCAAAAGGGCTAC

>Targ2 GCTTGGTCCCCTGGCCAAAAGTGTACG

>Targ3 CCTTGGTCCCTCCGCCGAAAGAAGGTG

>Targ4 GCTTGGTCCCCTGGCCAAAAGTGTCGT

>Targ5 GCTTGGTCCCCTGGCCAAAAGTGCCCG

>Targ6 CTTTGGTCCCAGGGCCGAAAGTGAATA

>Targ7 CCTTGGTCCCTTGGCCGAACGTCCACC

### human TCR-alpha C-region (Figure 4B)

*q*-value = 3.4E-5

>anchor GTACACGGCAGGGTCAGGGTTCTGGAT

>Targ1 TGCCTTTGCCGAAGTTGAGTGCATACC

>Targ2 TCCCTGATCCAAAGATTATCTTGGAAG

>Targ3 TGCCTGTCCCAAAGGTGAGTTTGTTTC

>Targ4 TCCCAGCGCCCCAGATTAACTGATAGT

>Targ5 TCCCCCTTGCAAAGAGCAGCTTCTGGC

>Targ6 TTCCTCCTCCAAAAGTTAGCTTGTTGC

>Targ7

TCCCTGTCCCAAAATAGAACTGGTTAC

>Targ8 TTCCTCTTCCAAAGTATAGCCTCCCCA

>Targ9 TTCCCTTTCCAAAGACCAGCTTTTCAG

>Targ10 TTCCCTGTCCGAAGATAAGCTTTCCTC

>Targ11 TCCCTGCTCCAAAGCGCATGTCATTGT

>Targ12 TTCCCTTCCCAAAGATCAGAGCAGTTC

>Targ13 TCCCAGATCCAAAGTAAAATTTGTTGA

>Targ14 TCCCTTGCCCAAAGATTAGTTTGCCTG

>Targ15 TTCCTCTTCCAAATGTAGGTATGTAGC

>Targ16 TTCCATCTCCAAACATGAGTCTGGCAT

>Targ17 TTCCACTCCCAAAAGTAAGTGCTCTCC

>Targ18 TTCCTTTTCCAAATGTCAGTTTATAGT

>Targ19 TGCCTGTTCCAAAGATGTATTTGTAGG

>Targ20 TTCCAGTTCCAAAGGTAACTTTCTGGT

>Targ21 TCCCTTGTCCAAATGTCAGCTTTCCAT

>Targ22 TCCCCTTCCCGAAAGTGAGTTGGTAAC

>Targ23 TGCCAGTTCCAAAGATGAGCTTGTTTG

### lemur Ig-heavy V-region (Figure 4B)

*q*-value = 1.3E-11

>anchor AGCCTGGGGGGTCCCTGAGACTCTCCT

>Targ0 AGTGACTACTACATGAGCTGGGTCCGC

>Targ1 AGCAGCTATGGGATGAACTGGGTCCGC

>Targ2 AGCAACTACTGGATGAGCTGGGTCCGC

>Targ3

AAGAACTATGAGATAAACTGGGTCCGC

>Targ4 AGCAGCTACTACATGCACTGGGTCCGC

>Targ5 AGCAGCTACGATATGAACTGGGTCCGA

>Targ6 AGTGACTACTACATGAACTGGGTCCGC

>Targ7 AGCAGCCATGGAATGCACTGGGTCCGC

>Targ8 AGCAGCTACGATATGAACTGGGTCCGC

>Targ9 AGCAGCTATGATATGCATTGGGTCCGC

>Targ10 AGTGACCACCACATGAGCTGGGTCCGC

>Targ11 GATGACTACCTCATGCACTGGATCCGC

>Targ12 AGCAGCTATGCCATGAGCTGGGTCCGC

>Targ13 AGTAGTTACTGGATGAACTGGGTCCGC

>Targ14 GATTACTATGGCATGAACTGGGTCCGC

>Targ15 ACCAATTTTGGGATGAACTGGGTCCGC

>Targ16 AGCAGCTATGGGATGCACTGGGTCCGC

>Targ17 ACCAGTTATGGGATGAACTGGGTCCGC

### lemur TCR-alpha C-region (Figure 4B)

*q*-value = 4.1E-7

>anchor TCAGCTGGTACACGGCGGGGTCAGGGT

>Targ0 AGTCTGGTCCCTGCTCCAAAGCGCAGA

>Targ1 AGCCTGGTCCCTGCTCCAAAAATCAAC

>Targ2 AGCAGAGTGCCAGTCCCAAAGATGAGC

>Targ3 ACGGTGGTTCCTTTCCCAAAGATCAAC

>Targ4 AGTTGGGTGCCAGTTCCAAACACGGGT

>Targ5

AACTGGGTCCCGGATCCAAAGGTCAGT

>Targ6 AGTTGTGTCCCTTTTCCAAAGGTGACT

>Targ7 AGTTTGGTCCCAGATCCAAAGTAAAAT

>Targ8 AATCTGGTCCCAGTCCCAAAGATGAGC

>Targ9 AGTCTGGTCCCTGATCCAAAGATTAGC

### Octopus bimaculoides Myo-VIIa (Figure 5A)

*q*-value = 4.0e-03

(reverse-complements shown in Figure 5A)

>anchor CCATTTTTGCTTTTTGTTTAAAATCCA

>target_1 ATTATATCACAAGTTATAAGGCATGCC

>target_2 ATTATATCTTAATAAATGGATACACTA

### fucoxanthin chlorophyll a/c protein, diatom (Figure 5C)

*q*-value = 6.0e-08

(reverse-complements shown in Figure 5C)

>anchor AAGTATCCAACAACGGCAAGCATGGAG

>target_1 (abundance order = 1) ATACGTCCGTGCTTGAGCTCGACAAAT

>target_2 (abundance order = 6) ATACGGCCGTGCTTGAGCTCGACAAAT

>target_3 (abundance order = 2) ATACGTCCGTGCTTGATCTCGACGTAT

>target_4 (abundance order = 4) ATACGTCCGTGCTTGATCTCAACGTAT

>target_5 (abundance order = 5) ATACGTCCGTGCTTGATCTCGACGTAC

>target_6 (abundance order = 3) ACACGTCCATGCTTAATTTCGACATAT

### *Zostera marina* NADPH quinone oxidoreductase subunit L (NdhL) (Figure 5D)

*q*-value = 6.5e-56

(reverse-complements shown in Figure 5D)

>anchor AATCGAAGCCAATTCATGATGATAGGC

>target1

GGCATGATAAGGAAGTAGAAGAAAGCA

>target2 GGCATGATAAGGAAGTAGAAGAAAACA

>target3 GGCATGACAAGGAAGTAGAAGAAAGCA

>target4 TTCGATCATGCAGTTCAATCAATGATC

### human MYL6 (Figure S4A)

>anchor AAGGTCCTCAGCCATTCAGCACCATGC

>P2_consensus1_macrophage GGACGAGCTCTTCATAGTTGATACAACCATTGCTGTCCTCATGCCCTGCCACCAGCATCTCTACTTCTTCCTCTGTCATCTTCTCACCCAGTGTGACAAGAACATGCCGGATTTC

>P2_consensus2_capillary GGACGAGCTCCGCCCCATGGGCCCGTCACCCCGACAGGATATGCCTCACAAACGCTTCATAGTTGATACAACCATTGCTGTCCTCATGCCCTGCCACCAGCATCTCTACTTCTTCC

>P2_target1 TGCCACCAGCATCTCTACTTCTTCCTC

>P2_target2 CACAAACGCTTCATAGTTGATACAACC

>P3_consensus_macrophage AAGGTCCTCAGCCATTCAGCACCATGCGGACGAGCTCTTCATAGTTGATACAACCATTGCTGTCCTCAT GCCCTGCCACCAGCATCTCTACTTCTTCCTCTGTCATCTTCTCACCCAGTGTGACAAGAACATGCCGGA

>P3_consensus_capillary AAGGTCCTCAGCCATTCAGCACCATGCGGACGAGCTCCGCCCCATGGGCCCGTCACCCCGACAGGATAT GCCTCACAAACGCTTCATAGTTGATACAACCATTGCTGTCCTCATGCCCTGCCACCAGCATCTCTACTT CTTCCT

### mouse lemur COX2 (cytochrome c oxidase subunit II) (Figure S5A)

(reverse-complements are shown in Figure S5A)

>anchor ATTTAGGCGCCCTGGGATAGCATCTGT

>target_1 TTCATGAATGTAGTACGTCTTCTGAAG

>target_2 TTCATGAATGTAATACGTCTTCTGAAG

### lemur IGLC3 with 97 targets (Figure S5B)

>anchor ACCGAGGGGGCGGCCTTGGGCTGACCT

>Targ0 GCCGAACACCCCAGTGCCACCACTCCT

>Targ1

GCCGAAGATATGACCACTCAGGCTGTC

>Targ2 GCCGAACACATGATTGTAGCTGCCATC

>Targ3 GCCGAATACATTAACACCACTGTTGTC

>Targ4 GCCGAACACATAACCATATGAATCACC

>Targ5 GCCGAACACACCACCACTGCTGTCCCC

>Targ6 GCCGAACACATTAACACCACCGTCCCA

>Targ7 GCCGAATACAGCACTGTTGTGCCACAC

>Targ8 GCCGAAGATATAAGTGTTCCTGCCCGC

>Targ9 GCCGAACACACCAACACCACTGCTGTC

>Targ10 GCCGAACACACCAACACCAGTTTCCCA

>Targ11 GCCGAAGATAACACCACTGTTGTCCCA

>Targ12 GCCGAACACACTGTAGCTGCCATCATA

>Targ13 GCCGAACACATAACCATATGAACCACC

>Targ14 GCCGAAGATATACTGAATGCTGCTCCC

>Targ15 GCCGAAGATATAAGTATTAGAGCTGCC

>Targ16 GCCGAACACCCGAGCATCAAGACTGCT

>Targ17 GCCGAATACATAAGCACTCAGGCTTTT

>Targ18 GCCGAACACCCGACCATTCAGGCTGCT

>Targ19 GCCGAATACATAAGTGCCACTGTTGGC

>Targ20 GCCGAAGATATACGCACTCAGGCTACT

>Targ21 GCCGAACACCTGACCACTCAGGCTACT

>Targ22 GCCGAACACACCAACACCACTGTTGTC

>Targ23 GCCGAACACCCAACTAGCACTGGCATC

>Targ24 GCCGAACACACCAGCACGTAGGCTGCT

>Targ25 GCCGAACACATGACCACTCAGGCTACT

>Targ26 GCCGAACACATGAGCACTCAGGCTTCT

>Targ27 GCCGAACACCCGACTGTAGCTGCCATC

>Targ28 GCCGAAGATATTAACACCACTGTTGTC

>Targ29 GCCGAAGATATCACTCAGGCTACTGTC

>Targ30 GCCGAACACCCAACTCTTAGAGCTGCC

>Targ31 GCCGAACACATCAGCACTGTTGTGCCA

>Targ32 GCCGAACACAAGATTGTAGCTGCCATC

>Targ33 GCCGAACACATAACTCTTAGAGCTGCC

>Targ34 GCCGAACACCCCAGTGCCACCACTCTT

>Targ35 GCCGAACACATCACCACTCAGGCTACT

>Targ36 GCCGAACACCCTGCTGTCATAGGACTG

>Targ37 GCCGAACACCCAATTAACACCACTGCT

>Targ38 GCCGAACACCCAAGCATCAAGACTGGT

>Targ39 GCCGAACACACGAGCATCAAGACTGCT

>Targ40 GCCGAACACCCAACCATATGAATCACC

>Targ41 GCCGAACACACCATGACCACTCAGGCT

>Targ42 GCCGAACACACCATAGTTTCCATAACC

>Targ43 GCCGAACACCGCATTAAGACTGCTGTC

>Targ44 GCCGAAGATATACTGGTTGCTGAACCA

>Targ45 GCCGAACACACCATGAGTACCAGTGCT

>Targ46

GCCGAATACATGACCACTCAGGCTGTC

>Targ47 GCCGAACACACCATCAAGACTGCTGTC

>Targ48 GCCGAAGATATAAGTGCCGCTGCCCGC

>Targ49 GCCGAACACATGACCACTCAGGCTTCT

>Targ50 GCCGAACACACCAGCATCAAGACTGCT

>Targ51 GCCGAAGATATAAGTGTTGCTGCCCGC

>Targ52 GCCGAACACCCAAGCATCAAGACTGCT

>Targ53 GCCGAACACACCATGACTCAGGCTGCT

>Targ54 GCCGAACACCCAAACACCACTGTTGTC

>Targ55 GCCGAACACATGAGCACTCAGGCTACT

>Targ56 GCCGAACAGACCACTCAGGCTACTATC

>Targ57 GCCGAAGATATACCCATATGAACCACC

>Targ58 GCCGAAGATATGACCACTCAGGCTACT

>Targ59 GCCGAACACCCAACCATATGAACCACC

>Targ60 GCCGAATACATAATTGTAGCTGTCATC

>Targ61 GCCGAACACACCACCACTCAGGCTGTC

>Targ62 GCCGAACACAAAATTAACACCACTGCT

>Targ63 GCCGAACACAGCACGCAGACTGCTGTC

>Targ64 GCCGAACACCCAAGTGCCGCTGCCCGC

>Targ65 GCCGAACACCCAGCACTGTTGTGCCAC

>Targ66 GCCGAAAACATAAGTCTTAGACCTGCC

>Targ67 GCCGAAGATATACGTATCAAGACTGCT

>Targ68 GCCGAAGATATTGTTTTCACTAACCCA

>Targ69 GCCGAAGATAGCACTGTTGTGCCACAC

>Targ70 GCCGAACACACGAGCACCCAGACTACT

>Targ71 GCCGAATACATGACCATTCAGGCTGCT

>Targ72 GCCGAATATATAACTCTTAGAACTGCC

>Targ73 GCCGAACACAAAACGGTTGCTGAACCA

>Targ74 GCCGAACATCCAACTCTTAGAGCTGCC

>Targ75 GCCGAACACCCAAGTCTTAGAGCTGCC

>Targ76 GCCGAACACATGACTGTAGCTGTCATC

>Targ77 GCCGAACACCCAATGGTTGCTGAACCA

>Targ78 GCCGAACACCCAAAGTGCCGCTGCCCG

>Targ79 GCCGAACACACCAGTCTTAGAGCTGCC

>Targ80 GCCGAAGATATTAACACCAGTTTCCCA

>Targ81 GCCGAACACACTGTAGCTGTCATCATA

>Targ82 GCCGAATACAAATGGTTGCTGAACCAC

>Targ83 GCCGAACACCCTATTAACACCACTGCT

>Targ84 GCCGAACACAGCATCAAGACTGCTGTC

>Targ85 GCCGAATACATAATCAAGACTGCTGTC

>Targ86 GCCGAACACACCACTCAGGCTACTATC

>Targ87 GCCGAAGATAGCATGAGTACCAGTATT

>Targ88 GCCGAAGATAAGACCACTCAGGCTACT

>Targ89 GCCGAACACAATAGCTGCCATCATAAG

>Targ90 GCCGAACACCTGATTGTAGCTGTCATC

>Targ91

GCCGAACACAAGACTAACACTGTCATC

>Targ92 CAGAGGCCTGTGTCCACCTGGGGAGCC

>Targ93 GCCGAACACACCTAGAGCTGCCATTCC

>Targ94 GCCGAATACATTAACACCACTGCTGTC

>Targ95 GCCGAATACATAATTGTAGCTGCCATC

>Targ96 GCCGAAGACAAACATCGACTGAGGCTC

### lemur TCR-beta J-region (Figure S5C)

>anchor CCGGGTCCCTGGCCCGAAGAACTGCTC

>Targ0 TGCCGCTGCAGATGTAGACGCCGCTGT

>Targ1 CGCAGAGATACAGGGCCGAGTCCCCCA

>Targ2 TGGCACAGAGGTACGTGGCGGAGTCTT

>Targ3 TGCTGGCACAGAGGTACGTGGCAGAGT

>Targ4 AGAGGAACAGGGCCGAGTCCCCCAGCG

### lemur TCR-gamma V-region (Figure S5D)

>anchor ACCCTCACCATTCACAATGTAGAGAAA

>Targ0 TGCCCGTGAACTCTTCAGTAATGGAAC

>Targ1 TGCCTCCTGGGAGTCTAGGAAACTCTT

>Targ2 TGCCTCCTGGGACTGACGACTTACCAA

>Targ3 TGCCTCCTGGGAGTTGAATTTTTATAG

>Targ4 TGCCTCCTGGGAGTTGCACAGTGTCAC

>Targ5 GCCCGTGAACTCTTCAGTAATGGAACA

>Targ6 TGCCTCCTGGGAGTCGCTCTCTAATAT

>Targ7 TGCCTCCTGGGAGTTGCACAGAAGATT

### *Octopus bimaculoides* carboxypeptidase D (Figure S6A)

(reverse-complements are shown in Figure S6A)

>anchor GGAATTAGAAGAAAAATCTATTATGAA

>target_1 AAATGTTTAGGCCAATATCTAAAGGCA

>target_2 AAATGTTTAGGAAAAATTTTCTGCCAA

### *Octopus bimaculoides* Upf2 (regulator of nonsense transcripts 2) (Figure S6B)

(reverse-complements are shown in Figure S6B)

>anchor GTATTGCACTGCATTGTACTGCACTGT

>target_1 CGCTGCTGCTGCTGCTGCTGCCAATTG

>target_2 CGCTGCTGCTGCTGCTGCCAATTGCCT

### *Octopus bimaculoides* netrin receptor / DCC (Figure S6C)

(reverse-complements are shown in Figure S6C)

>anchor TCTATTACAGCTATCATCAATACACTT

>target_1 TTGGATGTCTTCGTGTTCTCACTGCAG

>target_2 TTGGATGTCTTTGTGTTCTCACTGCAG

### HMG-box (diatom) (Figure S7A)

(reverse-complements are shown in Figure S7A)

>anchor TGCGGTCCTTGAATTCTTGCTTCTCTT

>target_1 TATCCGAAAGAGCCCTCCACATTTCAC

>target_2 CGTCCGTCAGAGCTCTCCACATTTCTC

### ferredoxin (diatom) (Figure S7B)

(reverse-complements are shown in Figure S7B)

>anchor ACGGCACGAGTAGGGAAGTTCAATTCC

>target_1 GGCTTCTTCAGCAGCGTCGACAATGAA

>target_2 GGCTTCTTCGGCAGCGTCGACAATGAA

