## Supplemental tables 1-5 for "SPLASH: a statistical, reference-free genomic algorithm unifies biological discovery": run.sh

```
datasets="
lemur_B_cell_spleen
lemur_natural_killer_T_cell_spleen
SS2_TSP1_B_cell
SS2_TSP2_T_cell
rotavirus_gut_metagenome
metatranscriptome_influenza_A
"
base_dir=/oak/stanford/groups/horence/zheludev/preprint_060722/figs
for dataset in ${datasets}
do
    nomad=${base_dir}/${dataset}/hits_filt.Pfam
    control=${base_dir}/${dataset}/decoys_filt.Pfam

    ls ${nomad}
    ls ${control}

    cp ${nomad} ${dataset}_nomad.Pfam
    if [[ -f ${control} ]]
    then
        cp ${control} ${dataset}_control.Pfam
    else
        touch ${dataset}_control.Pfam 
    fi
done

datasets="
france_coinfection
south_africa_covid_omnicron
stanford_preomicron
"

dir=/oak/stanford/groups/horence/kaitlin/nomad_paper/protein_domains/plotting/all_anchors

for dataset in ${datasets}
do 
    ## all anchors
    nomad=${dir}/protein_domains_nomad_${dataset}.tsv
    control=${dir}/protein_domains_control_${dataset}.tsv
    
    cp ${nomad} ${dataset}_nomad.Pfam
    cp ${control} ${dataset}_control.Pfam
done

cp /oak/stanford/groups/horence/kaitlin/nomad_paper/protein_domains/plotting/all_anchors/protein_domains_control_PRJNA817806_dehosted_NPS_france_coinfection_patient.tsv france_coinfection_control.Pfam
cp /oak/stanford/groups/horence/kaitlin/nomad_paper/protein_domains/plotting/all_anchors/protein_domains_nomad_PRJNA817806_dehosted_NPS_france_coinfection_patient.tsv france_coinfection_nomad.Pfam

dir=/oak/stanford/groups/horence/kaitlin/nomad_paper/protein_domains/plotting/anchors_effect_size_0.5
```
