## Supplemental tables 1-5 for "SPLASH: a statistical, reference-free genomic algorithm unifies biological discovery": run.sh

```
datasets="
HLCA_SS2_P2_macrophage_capillary
HLCA_SS2_P3_macrophage_capillary
lemur_B_cell_spleen
lemur_natural_killer_T_cell_spleen
lemur_B_cell_spleen
france_coinfection
south_africa_covid_omnicron
SS2_TSP1_B_cell
SS2_TSP2_T_cell
stanford_preomicron
rotavirus_gut_metagenome
metatranscriptome_influenza_A
"

base_dir=/oak/stanford/groups/horence/kaitlin/results_nomad/bulk_RNAseq/paper

for dataset in ${datasets}
do 
    file=${base_dir}/${dataset}/anchors_pvals.tsv
    echo ${dataset}_anchors_pvals.tsv
    cp ${file} ${dataset}_anchors_pvals.tsv
done
```
