## Supplemental tables 1-5 for "SPLASH: a statistical, reference-free genomic algorithm unifies biological discovery": run.sh

```
datasets="
HLCA_SS2_P2_macrophage_capillary
HLCA_SS2_P3_macrophage_capillary
"

base_dir=/oak/stanford/groups/horence/kaitlin/results_nomad/bulk_RNAseq/paper

for dataset in ${datasets}
do 
    file=${base_dir}/${dataset}/additional_summary.tsv
    echo ${dataset}_additional_summary.tsv
    cp ${file} ${dataset}_additional_summary.tsv
done
```
