## Supplemental tables 1-5 for "SPLASH: a statistical, reference-free genomic algorithm unifies biological discovery": run.sh

```
datasets="
lemur_B_cell_spleen
lemur_natural_killer_T_cell_spleen
SS2_TSP1_B_cell
SS2_TSP2_T_cell
"

base_dir=/oak/stanford/groups/horence/kaitlin/results_nomad/bulk_RNAseq/paper

for dataset in ${datasets}
do 
    file=${base_dir}/${dataset}/genome_annotations/genome_annotations_anchor.tsv
    echo ${dataset}_genome_annotations_anchor.tsv
    cp ${file} ${dataset}_anchor_genome_annotations.tsv
done
```
